# Spatial remodeling of the thymic stroma with age disrupts niches for T cell selection and tolerance

**DOI:** 10.64898/2026.07.29.741472

**Authors:** Andreas Tarcevski, Fatima Dhalla, Joshua W. Moore, Saulius Zuklys, Anja Kusch, Liliana Tchernev, Joshua Grier, Stefano Maio, Amir Khan, Thomas Barthlott, Mary Deadman, Adam E. Handel, Helen M. Byrne, Georg A. Holländer

## Abstract

Age-associated thymic involution is a major driver of immunosenescence, yet the cellular and spatial mechanisms coordinating age-related thymic remodeling remain incompletely understood. Combining single-cell transcriptomics, chromatin accessibility profiling, and spatial transcriptomics, we generated a spatially resolved multi-omic atlas of the aging mouse thymus. We show that thymic aging is not simply a process of epithelial loss, but a spatial reorganization of the stroma into new microenvironments, including age-associated epithelial states, a fibroblast-supported epithelial progenitor niche, and tertiary lymphoid structures. This remodeling displaces niches supporting positive and negative thymocyte selection and coincides with an intrinsic decline in cortical thymic epithelial cell function. Ligand-receptor mapping identifies medullary fibroblasts as a signaling hub sustaining epithelial progenitors and promoting tertiary lymphoid structure neogenesis, linking these hallmarks of thymic aging. Together, our findings reframe thymic involution as spatial stromal reorganization that links stromal remodeling to impaired thymopoiesis, central tolerance, and immune aging.

## Introduction

Aging is a time-dependent, multi-scale process marked by progressive functional decline across cells, tissues and organs, driven by hallmarks including telomere attrition, somatic mutation, epigenetic drift, altered intercellular communication and immunosenescence.^1,2^ Immunosenescence itself comprises thymic involution, chronic low-grade inflammation (‘inflammaging’), metabolic adaptation and hematopoietic change, collectively compromising immune organization and adaptability with age.^3,4^ The resulting decline in adaptive and innate immunity increases the susceptibility to infection, blunts vaccine responses, impairs tumor surveillance and promotes autoimmunity, while systemic inflammaging further elevates the risk of cardiovascular disease, dementia and frailty.^5–9^ Inflammaging is itself sustained by persistent senescent cells, oxidative tissue damage, latent infection, shifts in the effector– regulatory cell balance, and the accumulation of autoreactive T cells.^10–13^

The thymus is the primary lymphoid organ responsible for generating a functionally competent T cell compartment, yet it is also the first organ to undergo age-related decline.^14,15^ This process, termed thymic involution, begins in humans after the second year of life and in mice after postnatal week 6.^16–18^ Initially evident by a slow decline in thymic cellularity, thymic involution later leads to progressively compromised T cell maturation and selection.

Thymocyte selection depends on the spatial organization of hematopoietic and non-hematopoietic stromal cells that arrange within distinct niches and collectively generate a T cell repertoire responsive to foreign antigens yet tolerant to the body’s own “self” antigens.^19,20^ Cortical thymic epithelial cells (cTEC) attract hematopoietic precursors, commit them to the T cell lineage and mediate their positive selection, permitting the survival of thymocytes bearing T cell receptors (TCR) with sufficient affinity for self-peptide-MHC complexes (pMHC). Whereas, thymocytes that bind too strongly to self-pMHC, presented mainly by medullary epithelial cells (mTEC) and hematopoietic antigen-presenting cells (APC), are eliminated by negative selection. This occurs in two sequential waves that purge autoreactive specificities from the repertoire.^21^ Aging alters thymic cellularity and cell-type composition and generates novel age-associated cell states,^17,18,22,23^ correlating with reduced naïve T cell output, a contracted TCR repertoire, impaired central tolerance, and the emergence of intrathymic B Cell aggregates.^24–28^

A comprehensive understanding of thymic aging requires resolving how senescence alters the composition, spatial organization and functional interactions of the cell types that sustain thymopoiesis. Progress has been constrained by a trade-off between dissociative approaches, which enable deep molecular profiling at the cost of tissue architecture, and spatial methods, which preserve anatomical context but sacrifice molecular resolution. Here, we computationally combine single-cell and spatial multi-omics to resolve, at single-cell resolution, how aging affects higher-order thymic architecture and show that this reorganization, rather than epithelial loss alone, disrupts lympho-stromal crosstalk and impairs thymopoiesis.

## Results

### Aging generates novel epithelial states and remodels thymic stromal composition

To resolve how aging reshapes the thymic stroma, we compared thymi from young (4 to 8-week-old) and old (65 to 77-week-old) mice (precise ages used for each assay are provided in the methods). We first assessed the effect of aging on the cellular composition of the thymus using flow cytometry and single-nucleus (sn) multi-omics. Using flow cytometry, we determined the cellularity and phenotypes of hematopoietic (CD45+) thymic cells, thymic epithelial cells (TEC), and non-epithelial thymic stroma (NETS) (Extended Data Fig. 1a; Supplementary Fig. 1.1-1.2). Although total thymus cellularity was greatly reduced in old mice, TEC showed an age-related relative expansion. This increase was driven by intertypical TEC (ItTEC), whereas both mature mTEC and tuft-like TEC (TlTEC) declined. ItTEC comprise phenotypically and transcriptionally distinct TEC subsets that co-express features of cortical and medullary TEC and represent intermediate states within TEC differentiation trajectories.^17^ Medullary fibroblasts also increased in frequency, while other non-epithelial stromal populations remained unchanged.

To resolve these compositional shifts at molecular resolution, we profiled the stroma of young and old mice by combined snRNAseq and snATACseq, identifying 32 stromal subtypes and states originating from all three germ layers (Fig. 1a; Extended Data Fig. 1b; Supplementary Data DGE). Four TEC subtypes were selectively enriched in aged thymi (Fig. 1b,c). Consistent with prior nomenclature, we classified these as age-associated TEC (aaTEC1-4; Supplementary Fig. 1.3a).^23^ Although each aaTEC subset displayed a distinct transcriptional identity, all four shared signatures of cellular senescence, inflammaging, a senescence-associated secretory phenotype (SASP) and features of epithelial-to-mesenchymal transition (EMT) (Extended Data Fig. 1c).^29–32^ aaTEC1 expressed *Eya1*, a regulator of cell-fate specification and genomic stability,^33^ together with *Tmod1*, *Fam107a* and *Gas2*, which govern actin cytoskeletal organization and senescence-associated growth arrest (Fig. 1c).^34–37^ aaTEC2 preferentially expressed *Ncam1* and *Sema3a*, mediators of epithelial-mesenchymal cell-cell interactions,^38,39^ and *Notch3*, which promotes TGF-β-driven fibroblast persistence and fibrosis.^40^ aaTEC3 and aaTEC4 expressed *Rbfox1*, *Ank3* and *Ctnnd2*, regulators of cell polarity and mesenchymal remodeling,^41,42^ and both aaTEC subtypes expressed *Egfr* (Extended Data Fig. 1c), which sustains TEC progenitor proliferation and injury-induced regeneration.^43^ All aaTEC populations additionally expressed senescence-associated effectors: aaTEC1 and aaTEC3 expressed *Tgfb2* and *Rb1*, mediators of programed cell death and cell-cycle arrest,^44–46^ while aaTEC2 and aaTEC4 preferentially expressed SASP- and EMT-associated genes (Extended Data Fig. 1c). Beyond the aaTEC populations, most other TEC subtypes acquired age-associated features to varying degrees, including induction of transcription of the EMT regulators *Zeb1* and *Zeb2*, and of extracellular matrix and senescence-associated genes such as *Col1a2*, *Col3a1*, *Igfbp4* and *Igfbp7,*^47,48^ linking transcriptional aging signatures to structural remodeling of the thymic microenvironment (Supplementary Fig. 1.3b).

**Fig. 1:**
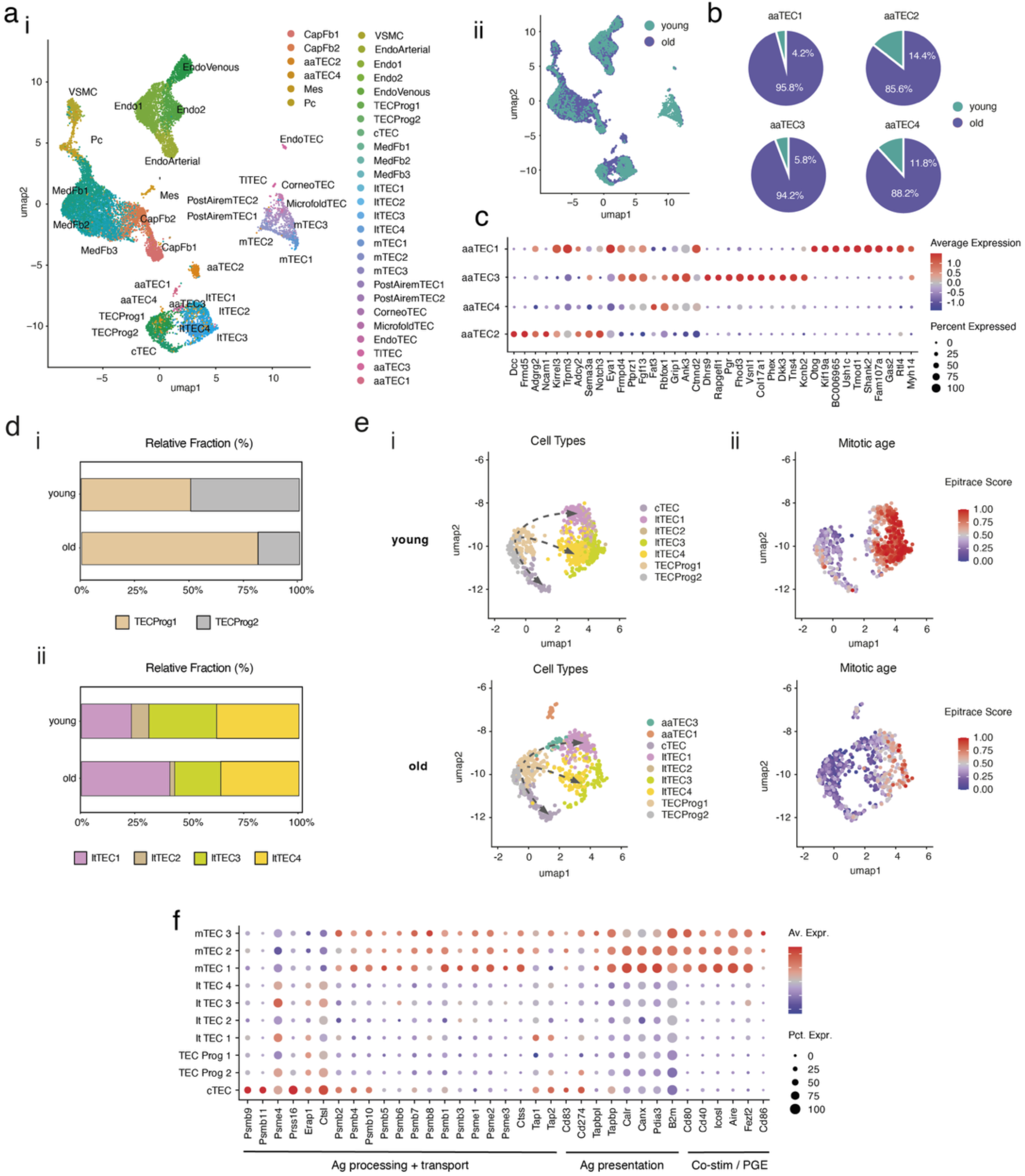
Aging generates novel epithelial cell states and alters developmental dynamics of TEC progenitors. a) Joint UMAP projection of the annotated thymic stromal cell compartment profiled via snRNA/ATACseq in young and old mice. (i) Cells grouped by cell type annotation. CapFb = capsular fibroblasts; aaTEC = age-associated TEC; cTEC = cortical thymic epithelial cells; Endo = endothelial cells; ItTEC = intertypical TEC; MedFb = medullary fibroblasts; mTEC = medullary TEC; Pc = pericytes; TECProg = TEC progenitors; TlTEC = tuft-like TEC; VSMC = vascular smooth muscle cells. (ii) Cells grouped by sample age; teal = young, purple = old. b) Relative proportion of young (teal) and old (purple) cells within age-associated TEC subtypes (n_tot_: aaTEC1 = 48, aaTEC2 = 197, aaTEC3 = 52, aaTEC4 = 209). c) Top differentially expressed genes of age-associated TEC populations defined against the full stromal cell compartment. d) Comparison of the relative proportions of (i) TEC progenitor and (ii) intertypical TEC subtypes in young and old thymi. e) (i) UMAP projection and (ii) mitotic age representation of selected TEC subtypes from young vs old thymi; arrows represent the gradual increase in mitotic age. f) Heatmap showing expression of genes associated with antigen (Ag) presentation, processing, transport along with co-stimulatory molecules (Co-stim) and molecules involved in promiscuous gene expression (PGE) in TEC progenitors, intertypical TEC, cortical and medullary TEC subtypes; Av. Expr. = average expression, Pct. Expr = percentage of cells expressing.

Integration of transcriptomic and chromatin accessibility data further resolved two TEC progenitor (TECProg) subtypes exhibiting distinct transcriptomic profiles yet retaining signatures of previously identified progenitor populations (Extended Data Fig. 1d; Supplementary Fig. 1.3c,d)^49^ along with several ItTEC populations marked by the expression of *Ccl21a*, *Krt5* and *Lifr* (Extended Data Fig. 1b,e). TECProg1 and TECProg2 were equally abundant in young thymi, but only the frequency of TECProg1 increased with age (Fig. 1d i). Among the ItTEC populations, the relative frequency of ItTEC1 expanded in old mice while ItTEC2 and ItTEC3 declined and ItTEC4 remained stable (Fig. 1d ii). To define the developmental relationships between these populations, we applied EpiTrace, which infers replicative age from chromatin accessibility and cross-confirmed using established trajectory inference tools (Fig. 1e; Extended Data Fig. 1f).^50,51^ TECProg1 exhibited the lowest replicative age, consistent with a progenitor identity, while TECProg2, displayed a higher replicative age and marked transcriptional similarity to cTEC regardless of chronological age (Extended Data Fig. 1g; Supplementary Fig. 1.3e). Trajectory and replicative-age analyses together further indicated that TECProg1 gives rise to ItTEC1 and ItTEC4 (Fig. 1e; Extended Data Fig. 1g), which mature sequentially into ItTEC2 and ItTEC3, respectively, given the transcriptomic similarity between these TEC subtypes (Extended Data Fig. 1e). Notably, aaTEC3 occupied a transcriptional space between TECProg1 and ItTEC1 in old thymi, suggesting an age-associated intermediate that preferentially accumulates along this trajectory (Fig. 1e). Neither the TEC progenitor populations nor the ItTEC subsets expressed the antigen-processing and presentation machinery characteristic of mature cortical and medullary TEC (Fig. 1f), implying that they remain functionally immature despite their expansion with age. Taken together, the comparison of the transcriptomes of young and old thymi at single-cell resolution identified previously unrecognized age-related stromal subtypes and further enabled the characterization of differentiation trajectories for various TEC subtypes.

### A spatial multi-omic atlas anchors stromal cell states to thymic anatomy

To determine where these populations reside within the thymus, we performed single-cell-resolved spatial transcriptomics on thymic sections of young and old mice and computationally integrated the resulting data with our stromal snRNAseq/snATACseq atlas and a published thymocyte CITEseq dataset,^52^ imputing genome-wide transcriptional and chromatin profiles onto each spatially resolved cell (Extended Data Fig. 2.1a,b). This integration was validated by the concordance of cell-type marker genes identified separately within the spatial and single-cell datasets, and by correct anatomical reconstruction of differentially expressed mTEC markers (e.g., *Cadm2*, *Syt1*, *Adgrg1*) absent from the Xenium panel (Extended Data Fig. 2.1c,d; Supplementary Fig. 2.1a,b), confirming that cross-modal integration reliably enhanced molecular resolution without compromising spatial accuracy.

Mapping the integrated dataset onto the tissue architecture enabled high-confidence anatomical localization of distinct cell subtypes previously defined only by transcriptional profiles (Fig. 2a; Supplementary Fig. 2.2-2.9). Comparing young and aged sections confirmed a broad depletion of medullary epithelial populations, including mimetic TEC, alongside an expansion of B cells and mesenchymal populations including fibroblasts (Fig. 2b). In young mice, both progenitor populations exhibited subcapsular and (cortico-)medullary localization, with TECProg2 showing slight predominance within the medullary compartment. With age, TECProg1 expanded markedly in the inner thymus, accumulating adjacent to the corticomedullary junction (CMJ), whereas TECProg2 progressively declined in frequency. (Fig. 2c). All four ItTEC populations localized to the CMJ and central medulla irrespective of subtype and retained this distribution across both ages, a finding indicative of a spatially conserved function, such as CCL21 production (Fig. 2d). Two aaTEC populations were detectable in sufficient numbers for spatial quantification in old mice and localized to the deep cortex adjacent to the CMJ. This spatial atlas therefore assigned precise anatomical addresses to age-associated and progenitor epithelial states, revealing that aging altered their abundance and relocated a key progenitor population towards the CMJ, a finding we return to when defining the fibroblast-rich niche that sustains them.

**Fig. 2:**
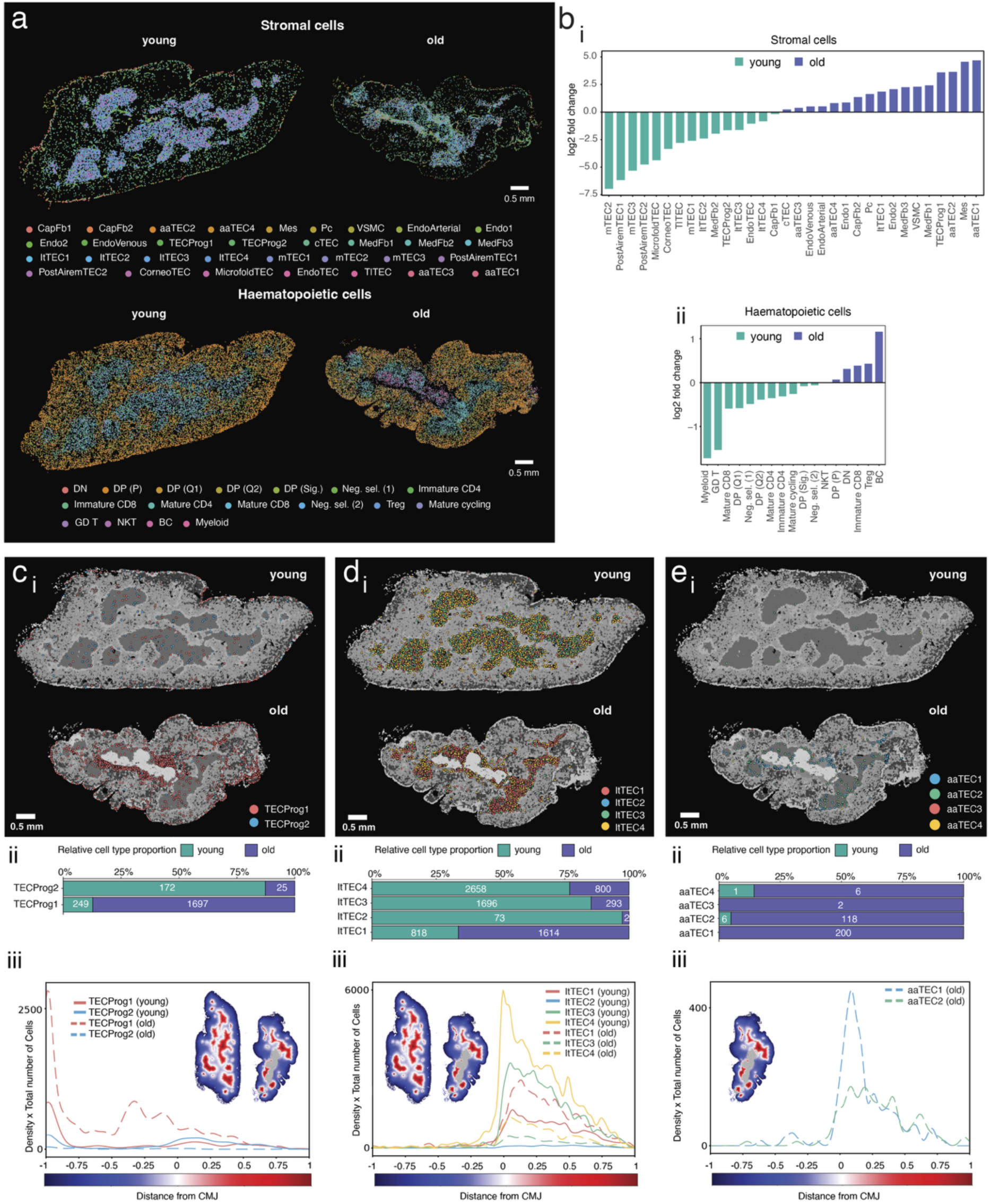
Cross-modal integration of single-cell and spatial transcriptomics data reveals in-situ location of heterogeneous cell sub-types. snRNA/ATACseq and CITEseq data were cross-modally integrated with single-cell spatial transcriptomics (Xenium) of young and old thymus sections a) Spatial projections of cross-modal integration informed stromal and hematopoietic cell type locations in the young and old thymus section. Cell type abbreviations: DN = double negative thymocytes; DP = double positive thymocytes; Neg. sel. = negatively selected thymocytes; immature/mature CD4/8 = immature/mature, single positive thymocytes for either CD4 or CD8; Treg = regulatory T cells; GD T = gamma-delta T cells; NKT = natural killer T cells; BC = B cells. b) Bar chart showing changes in proportions of (i) stromal cell and (ii) hematopoietic cell subtypes within the spatial transcriptomics dataset; expressed as Log 2-fold change of relative proportions in old vs young thymus tissue; teal bars indicate cell types relatively increased in young and purple bars indicate cell types relatively increased in old. c-e) Analysis of spatial position and abundance of c) TEC progenitors, d) intertypical TEC and e) age-associated (aa) TEC in young and old thymus tissue sections, showing (i) their in situ positions (ii) absolute cell counts and relative representation in young vs old, and (iii) cell count adjusted probability density functions, which quantify their position and abundance with respect to the CMJ.

### Mimetic TEC form discrete medullary niches that contract with age

We next asked whether the medulla itself, the principal site of tolerance induction, is spatially organized into discrete niches, and how this organization is affected by age. Mimetic TEC are a heterogeneous population of medullary epithelial cells that mimic peripheral cell types, recapitulating their gene expression programs to display tissue-restricted antigens (TRAs) and thus support central tolerance induction.^53,54^ Because mimetic TEC were markedly depleted in aged thymi, we first defined their baseline spatial organization in young mice. CorneoTEC, microfold TEC, endocrine TEC (endoTEC) and tuft-like TEC (TlTEC) localized predominantly to the deep medulla, with modest enrichment at the CMJ (Fig. 3a), forming discrete cellular aggregates rather than being uniformly dispersed (Fig. 3b; Supplementary Fig. 3a). Quantitative spatial analysis confirmed that mimetic TEC were non-randomly distributed within the medulla, forming localized clusters (Moran’s *I*, *Z* = 3.04, *P* = 0.0024) that defined discrete medullary hotspots of mimetic cells (Fig. 3c). Individual mimetic TEC subtypes exhibited strong short-range homotypic co-localization (Fig. 3d). In addition, most mimetic TEC subtypes showed significant heterotypic co-localization over distances of up to 100μm, except for microfold and endo TEC (Fig. 3e). These findings demonstrate that mimetic TEC are organized into discrete multicellular niches rather than randomly distributed throughout the medulla.

**Fig. 3:**
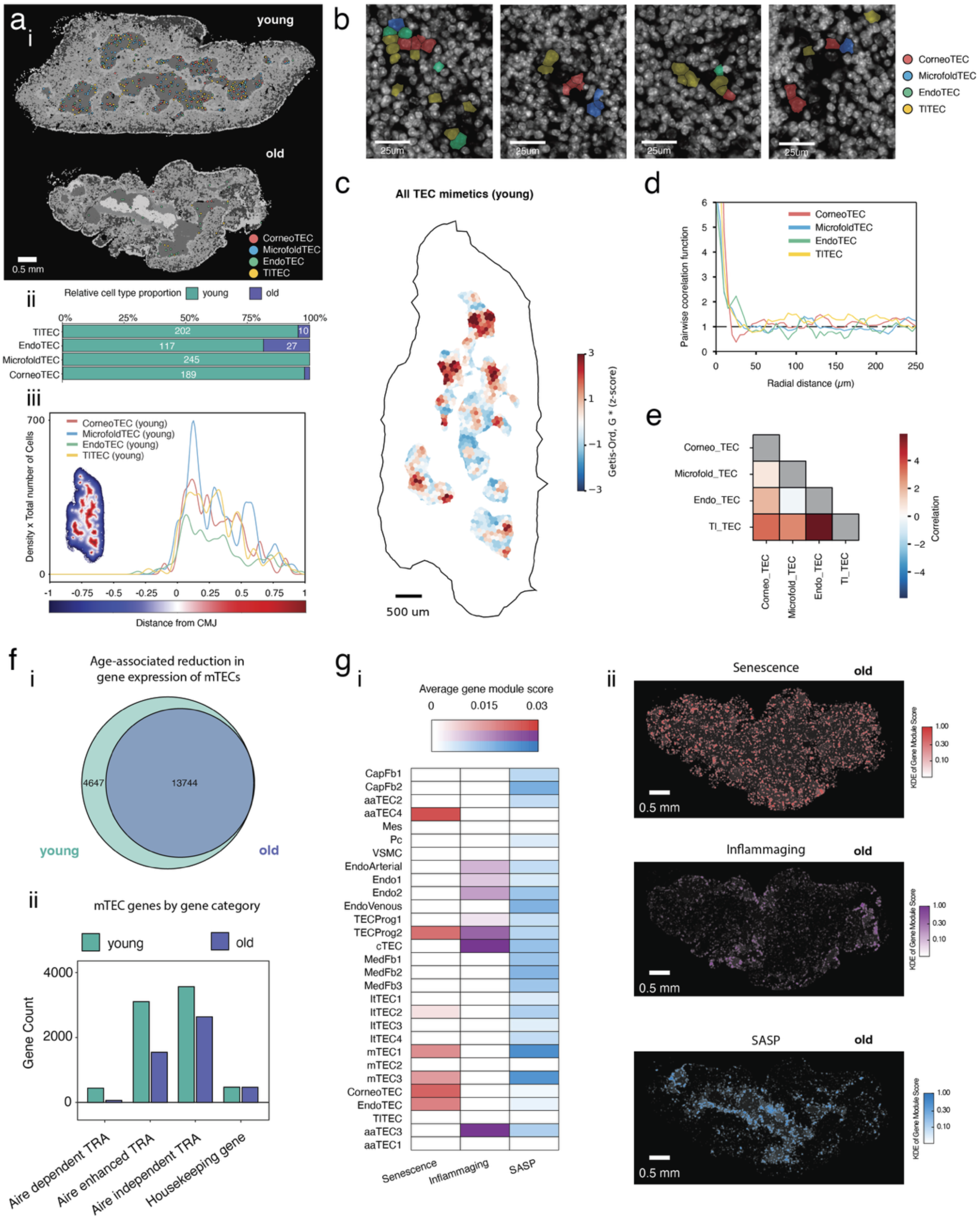
Mimetic TECs cluster spatially and the thymus microenvironment is differentially impacted by signatures of aging. a) (i) Projected locations of mimetic TEC, (ii) absolute cell counts and relative representation in young vs old thymus tissue sections, and (iii) in the young sample only cell count adjusted probability density functions, which quantify their normalized position with respect to the CMJ. b) Qualitative examples of mimetic TEC locations in the young thymus tissue section. c) Spatial distribution of mimetic TEC populations quantified using the local Getis–Ord Gi* statistic across hexagonal regions within the medulla. Gi* statistics were computed relative to a null model of complete spatial randomness (CSR). Red indicates regions of significant local enrichment (hotspots) of mimetic TEC, whereas blue indicates regions of significant local depletion (coldspots). d) Homotypic spatial association of individual mimetic TEC subtypes quantified using the pair correlation function (PCF). The dashed grey line indicates the expectation under CSR. e) Heterotypic spatial co-occurrence of mimetic TEC subtypes quantified using the quadrat correlation matrix (QCM) computed over the same hexagonal lattice shown in **c**, with pairwise correlations evaluated relative to CSR. f) snRNAseq-based quantification of gene expression in all mTEC populations (including mTEC1-3, Post-Aire mTEC1-2 and mimetic TEC) in young vs old thymi showing (i) the total number of genes detected and (ii) the number of genes encoding Aire-dependent, Aire-enhanced, and Aire-independent tissue restricted antigens (TRAs), compared to housekeeping genes.^55^ g) Quantification and localization of aging signatures associated with cellular senescence, inflammaging, and senescence-associated secretory pathways (SASP) in the old thymus section only, showing (i) average gene module scores for each aging-associated process across stromal cell subtypes within the old snRNAseq dataset.^29,30^ (ii) Kernel density map of gene module scores for each aging-associated process across all cells of the old spatial transcriptomics dataset.

Consistent with the age-associated reduction in mTEC abundance (Extended Data Fig. 1a iii), mTEC also exhibited a reduced expression of tissue-restricted antigens (TRAs) with age, particularly those dependent on AIRE for transcription (Fig. 3f).^55^ To determine how aging is manifested across the tissue, we next examined the spatial distribution of distinct hallmarks of cellular aging. Gene module scoring for cellular senescence, inflammaging and SASP showed that nearly all epithelial and mesenchymal populations expressed at least one aging hallmark, but that these hallmarks were not uniformly distributed (Fig. 3g). Instead, they formed discrete spatial domains, leaving substantial regions of the aged thymus comparatively spared (Fig. 3g ii).

Together, these findings indicate that the medullary compartment is organized into functionally distinct spatial niches that support immune tolerance and are differentially marked by molecular features of aging, suggesting that aging not only erodes niche function but also reshapes their higher-order spatial organization.

### Aging generates new higher-order microenvironments, including a fibroblast niche and B cell aggregates

To define this higher-order spatial organization systematically, we integrated the spatial transcriptomic data for stromal and hematopoietic cells and performed unsupervised neighborhood clustering using Multiscale Spatial Analysis (MuSpAn).^56^ This analysis groups cells into recurrent neighborhoods based on local composition and quantifies their spatial relationships (Supplementary Fig. 4a). Across young and aged thymi, we identified 15 microenvironmental regions common to both ages, comprising three capsular/subcapsular, five cortical, two CMJ, and five medullary regions (Fig. 4a,b). Several regions occupied the same histological compartment yet differed in cellular composition, indicating that cortex and medulla are each composed of multiple functionally distinct stromal niches rather than single homogeneous domains. Strikingly, two additional regions were detected exclusively in aged thymi. Region 16 (‘Fibrous region I’, Fibr I) was enriched for medullary fibroblasts together with TECProg1 and ItTEC populations but relatively depleted of thymocytes. Region 17 comprised prominent B cell aggregates (BCaggr), likewise markedly depleted of thymocytes, and was largely surrounded by region 16. These two regions represented microenvironments that arose specifically during thymic aging, rather than age-related contraction or expansion of pre-existing niches.

**Fig. 4:**
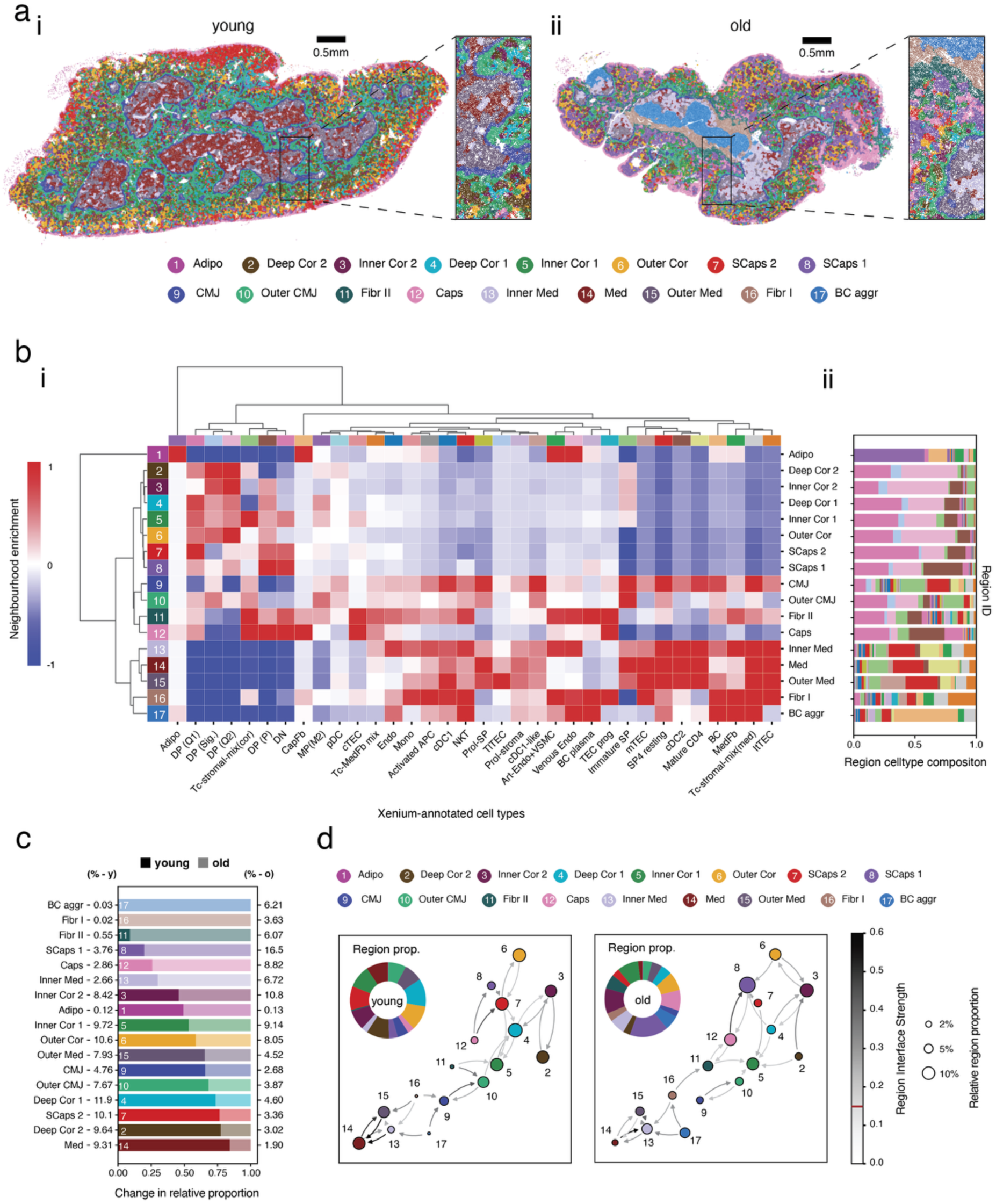
Aging remodels the thymic microenvironment and generates fibroblast- and B cell-rich niches. a) Representative young (i) and aged (ii) thymus sections with cell centers colored by spatial neighborhood identity. Neighborhoods were identified by unsupervised clustering of local cell type composition within three-hop neighborhoods defined on a Delaunay triangulation of cell centroids. b) (i) Cell type enrichment heat map and (ii) cell type composition of the identified spatial neighborhoods. d) Relative abundance of spatial neighborhoods in the young and aged thymus. e) Directed spatial neighborhood interaction network summarizing shared interfaces between neighborhoods across both age groups. Node size is proportional to neighborhood abundance, and edge color indicates the relative interface strength (proportion of shared boundary) between adjacent neighborhoods. Only interfaces with strength >0.15 are shown. Pie chart insets indicate the relative abundance of neighborhoods in each sample.

Aging also substantially altered the relative abundance of the conserved niches: deep cortical and some medullary regions contracted while capsular/subcapsular domains and other medullary regions expanded and exhibited subtle changes in the underlying cell type signatures (Fig. 4b,c). These changes also profoundly remodeled the adjacencies of distinct niches leading to the formation of novel interfaces that range from central aspects of the tissue such as the BCaggr (Region 17) via the fibrous regions I and II (Regions 16 and 11, respectively) into (sub)capsular areas (Regions 7 and 8) (Fig. 4d). These alterations coincided with an increased abundance of mesenchymal cells across aged medullary regions and a redistribution of areas with subcapsular signatures reaching deep into the cortex and occasionally close to the CMJ (Fig. 4a,b; Supplementary Fig. 4b,c).

Thymic aging therefore involves not only altered cellular composition but a fundamental reorganization of tissue architecture: existing niches contract, redraw their boundaries, and are joined by two entirely new microenvironments. Because this reorganization directly alters the structural scaffold along which thymocytes migrate and mature, we next asked whether the developmental checkpoints of thymocyte selection are themselves displaced within this altered landscape.

### Aging displaces the anatomical and functional checkpoints of positive and negative selection

To link the remodeled architecture to thymocyte development, we reconstructed the developmental trajectory of double-positive (DP) and single-positive (SP) thymocytes by combining a public CITEseq dataset^52^ of 4 to 8 week-old mice with pseudo-time analysis (Fig. 5a i,ii).^51^ Cross-modal integration with our spatial data faithfully recapitulated the canonical centripetal migration of thymocytes beyond the initial stages of intrathymic maturation. This developmental journey was initiated in the cortex and progressed across the CMJ into the medulla independent of the age of the thymi (Fig. 5a iii). This analysis enabled us to map the principal checkpoints of thymopoiesis -positive and negative selection-onto the tissue according to both pseudo-time and their spatial position relative to the CMJ (Fig. 5b,c).

**Fig. 5:**
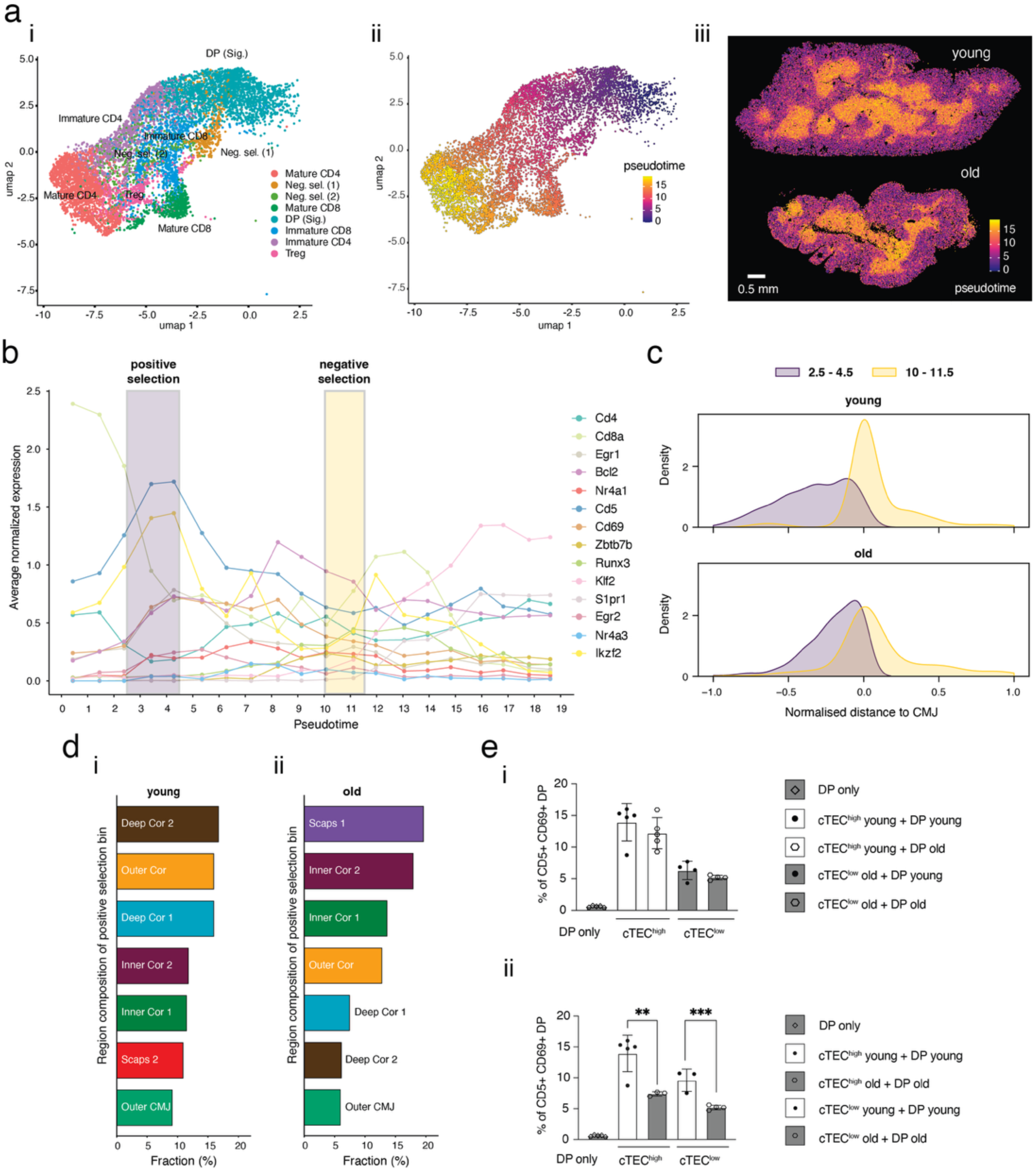
Aging alters the sites of T cell selection checkpoints and impairs the ability of cTEC to mediate positive selection. a) (i) UMAP projection of a literature-adapted, CITEseq-profiled, developing thymocyte dataset at 4 weeks of age showing the original cell type annotations^52^ and (ii) monocle3-based pseudo-time trajectory. (iii) Projection of the individual pseudo-time scores onto thymocytes within the young and old thymus tissue sections. b) Binned average expression profiles of selected genes with relevance to thymocyte maturation and selection plotted against their pseudo-time scores. Highlighted are positive (purple; 2.5-4.5) and negative (yellow; 10-11.5) selection windows. c) Quantification of the spatial position of thymocytes falling into the positive and negative selection windows, expressed as normalized distance to the CMJ in the young (upper) vs old (lower) thymus. d) Relative neighborhood composition of thymocytes (according to previous neighborhood definitions in Fig. 4a) within the positive selection bin. e) Thymic epithelial cell cultures, performed by co-culturing cTEC and pre-selection DP thymocytes from young and old thymi. Bar graphs show the up-regulation of CD5 and CD69 within young and old DP thymocytes after 48 hours of culture in the absence (DPs only) or presence of cTEC expressing high levels of MHCII (cTEC high) or low levels of MHCII (cTEC low). Data are represented as mean +/-SD. Unpaired student’s t-test with * p ≤0.05, ** p ≤0.01, *** p ≤0.005.

In young mice, thymocytes undergoing positive selection (pseudo-time 2.5–4.5), identified by induction of *Cd5*, *Cd69*, *Egr1* and *Egr2* transcription downstream of TCR signaling, were distributed throughout the cortex with increasing density toward the CMJ. In aged thymi, positive selection was displaced further toward the CMJ (Fig. 5c), corresponding to altered use of the underlying stromal niches: whereas in young mice positive selection occurred predominantly in outer and deep cortical regions, in aged mice a substantial fraction of this selection relocated to subcapsular and inner cortical regions (Fig. 5d; Fig. 4a). Nearly half of all positive-selection events occurred within a different stromal microenvironment in aged compared with young thymi, demonstrating extensive spatial displacement of this checkpoint. Additionally, a targeted spatial analysis of cTEC proficient at performing positive selection (perinatal cTEC, or PS+ cTEC), identified by markers such as *Cd83, Cd274, Cd40, Tnfrsf14,* and *Icam1*^57^, revealed 6 distinct, local cTEC neighborhoods along with age-related changes in both their relative abundance and global spatial arrangement (Fig. 4a; Extended Data Fig. 5a-c).

We next tested whether this spatial displacement was accompanied by an intrinsic functional decline in cTEC. Comparative transcriptomics identified age-dependent changes in cTEC gene expression (Extended Data Fig. 5b), and functional co-culture of cTEC (EpCAM+Ly51+UEA-) from young or aged mice with pre-selection DP thymocytes (CD4+CD8+CD69-CD5-) showed that induction of CD69 and CD5 after 48 hours was consistently more efficient with young cTEC, regardless of the age of the responding thymocytes (Fig. 5e i). Age-matched co-cultures further confirmed that both cTEC^hi^ and cTEC^lo^ from young mice induced significantly greater CD69/CD5 expression than their aged counterparts (Fig. 5e ii). These findings show that aging both relocates the anatomical niche supporting positive selection and intrinsically impairs the epithelial cells that execute it.

Negative selection, identified by induction of *Egr2*, *Nr4a1*, and *Ikzf2* following lineage commitment (pseudo-time 10-11.5), showed a broadly conserved cortico-medullary localization between young and aged thymi but extended further into central aspects of the medulla with age (Fig. 5b,c). Notably, the specific global and local microenvironments hosting thymocytes undergoing negative selection differed substantially between the age groups (Fig. 6a,b), demonstrating that the anatomical position of a checkpoint and the cellular neighborhood executing it are not equivalent, and prompting us to directly examine how the cellular partners of maturing thymocytes change with age.

**Fig. 6:**
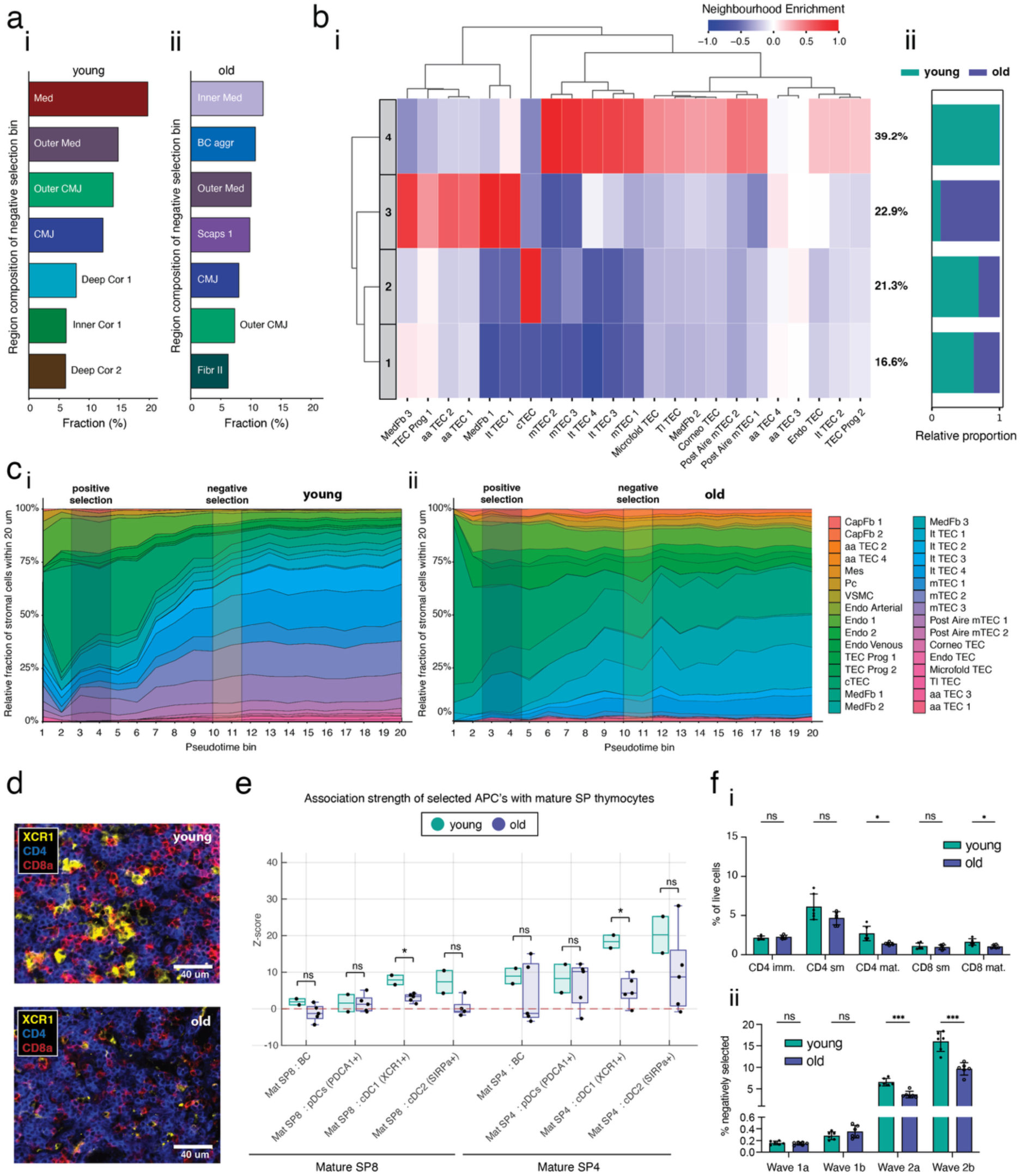
Aging remodels the cellular neighborhoods surrounding T cells at distinct stages of maturation. a) Relative neighborhood composition of thymocytes (according to previous neighborhood definitions in Fig. 4a) within the negative selection bin (Fig. 5b) for young (i) and old (ii) mice. b) (i) Heatmap showing the local enrichment of stromal cell populations surrounding thymocytes undergoing negative selection, quantified within 30μm neighborhoods and clustered by k-means (k = 4). (ii) Bar chart depicts the relative contribution of young and old neighborhoods. c) Relative proportion of stromal cells within a 20µm proximity of thymocytes with distinct pseudo-time scores in the (i) young and (ii) old (right panel) spatial transcriptomic thymus tissue sections. d) Multiplex-Immunohistochemical (mIHC) images of young (upper panel) and old (lower panel) thymus tissue sections (XCR1 in yellow, dendritic cells (DCs); CD4 in blue, mature single positive CD4 T cells (Mat. SP4) and CD8a in red, mature single positive CD8 T cells (Mat. SP8)); images are representative of 2 independent experiments. e) Quantification of pairwise spatial association strength in young (n = 2) and old (n = 5) thymus sections profiled by mIHC. Z-scores indicate a spatial enrichment relative to random distribution; * indicates p < 0.05 (exact two-sided permutation test). f) Young and old mice thymi were analyzed flow cytometrically for (i) relative frequencies of different singe positive CD4 or CD8 T cell populations, and (ii) negatively selected thymocytes stratified into first and second waves respectively (wave 1a = TCRb^int/hi^CCR7-DNs+DPsHelios+PD1+; wave 1b = CD4immature Helios+PD1+; wave 2a = CD4semimature Helios+; wave 2b = CD4mature Helios+, see Supplementary Fig. 1.2 for gating strategy); data is represented as mean +/- standard deviation. Statistical analyses were performed on the flow cytometric data using a two-tailed, unpaired student’s t-test for singular comparisons, and multiple unpaired t-tests with Holm-Sidak correction for multiple comparisons; * p ≤0.05, ** p ≤0.01, *** p ≤0.005. Abbreviations: imm. = immature, sm = semi-mature, mat. = mature.

### Maturing thymocytes are progressively reassigned from epithelial to fibroblast neighbors

To resolve these changing cellular partnerships, we quantified the stromal cells within a 20µm radius of thymocytes at each successive pseudo-time interval (Fig. 6c). In both age groups, immature DP thymocytes remained closely associated with cTEC, consistent with their role in positive selection. As thymocytes matured, however, their stromal neighborhood diverged sharply with age. In young mice, maturing thymocytes progressively associated with medullary epithelial populations, including, Aire+ mTEC1-3, post-Aire mTEC and multiple mimetic TEC populations, whereas in aged mice, maturing thymocytes instead increasingly localized near medullary fibroblasts (MedFb1, MedFb3), together with only a restricted subset of epithelial cells (ItTEC1, ItTEC4). Because ItTEC express substantially lower levels of antigen- rocessing and presentation genes than mature cortical and medullary TEC (Fig. 1f),^57^ this shift implies that maturing thymocytes in the aged thymus increasingly encounter cells with a diminished intrinsic capacity for antigen presentation. Furthermore, multiplex imaging of XCR1+ dendritic cells in young and aged thymi revealed age-associated changes in the spatial association between type-1 conventional dendritic cells and single positive CD4 and CD8 thymocytes (exact two-sided permutation test, P < 0.05; Fig. 6d,e), suggesting that antigen-presenting partnerships beyond the epithelial compartment are also reconfigured with age. Consistent with this reorganization of cellular neighborhoods important for medullary negative selection, flow cytometric analysis revealed a reduction in single positive CD4 thymocytes undergoing negative selection in the aged thymus (Fig. 6f).

Together, these results show that aging remodels not only the anatomical location of thymocyte selection but the cellular identity of the niche in which it occurs, raising the question of what becomes of the medullary space vacated by mature epithelial cells, and what draws thymocytes and stroma into the fibroblast-rich territory identified in the neighborhood analysis (Fig. 4a). We address this question in the sections that follow.

### Fibroblast-adjacent B Cell aggregates in the aged thymus constitute tertiary lymphoid structures

Among the age-specific niches identified above, Region 17 comprised *Cd19*-positive B Cell aggregates (BCaggr) of variable size (Fig. 4a; Fig. 7a i). Independently, immunofluorescence analysis mapped those B cell aggregates throughout the cortex and medulla and frequently located them adjacent to fibrous structures (Extended Data Fig. 7a,b), a spatial relationship suggesting a mechanistic link between fibroblast remodeling and B Cell accumulation that we tested directly.

**Fig. 7:**
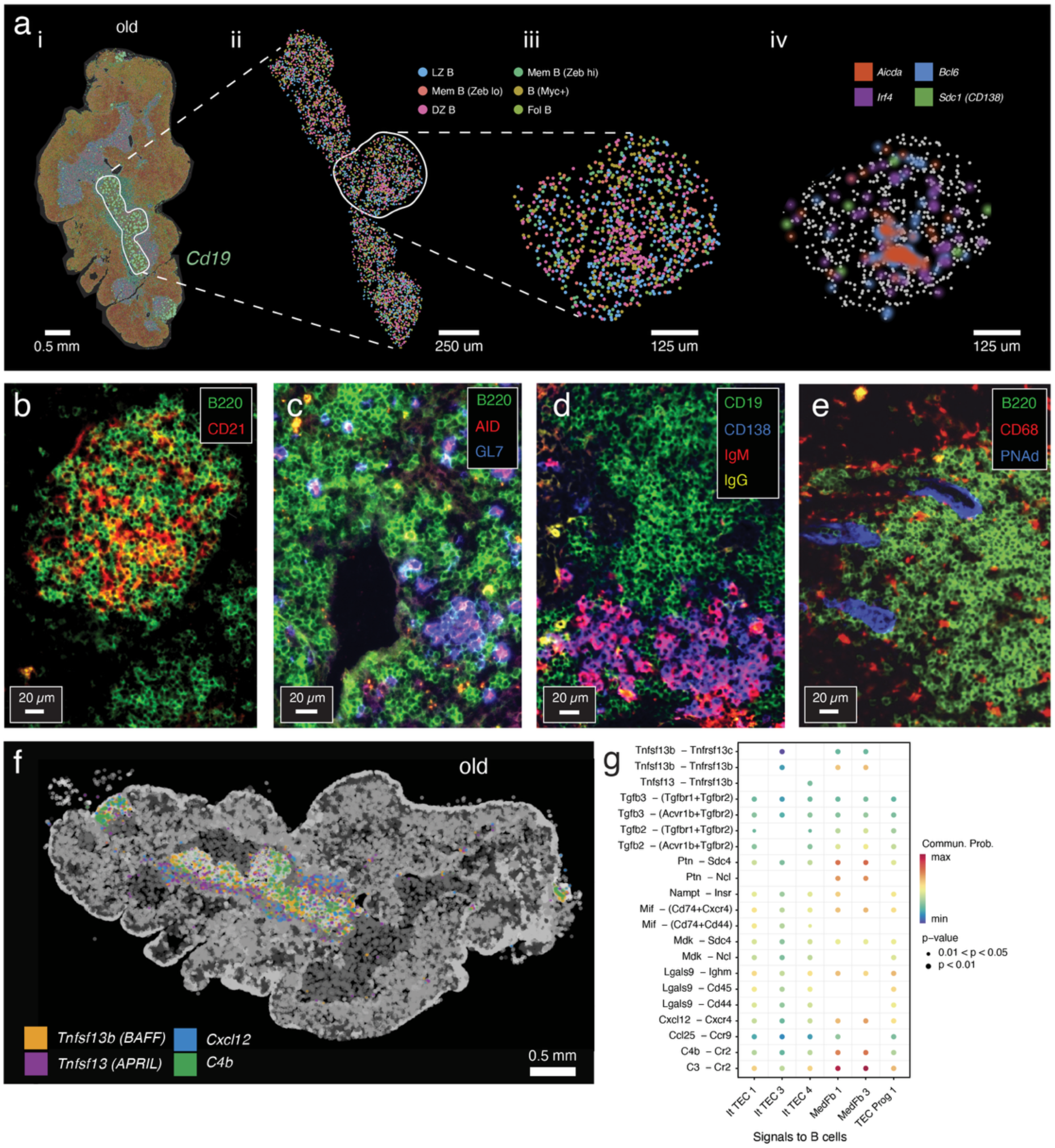
Medullary fibroblasts contribute to the formation of thymic TLS that show germinal-center-like features. a) (i) Spatial transcriptomics image of the old thymus lobe to illustrate the location of B cell aggregates; distinct colors mark distinct cell clusters, green spots mark location of *Cd19* transcripts, the white selection demarcates a centrally located B cell aggregate. (ii) Zoomed in representation of the central B cell aggregate; distinct colors correspond to individual B cell sub types which were annotated using a splenic B cell, scRNAseq reference^58^; LZ B = light zone B cell, Mem B = memory B cell, DZ B = dark zone B cell, Fol B = follicular B cell. (iii-iv) Further magnified area of the central B cell aggregate displaying spatial gene expression density maps of selected genes relevant during germinal center (GC) reactions. b-e) Immunofluorescence images of aged thymic B Cell aggregates stained for markers including B220, CD19, CD21, CD68, GL7, AID, CD138, IgM, IgG and PNAd illustrating spatial organization of different cell subtypes. Images are representative of 2 independent experiments. f) Spatial gene expression density map illustrating the localization of transcripts encoding selected B cell relevant cytokines, chemokines and complement compounds. The representation of spatial gene expression is restricted to the B cell and fibrous area. g) Spatially-restricted ligand:receptor analysis quantifying the communication probability via distinct paracrine signaling avenues (y-axis) for selected stromal cell types (x-axis; source cells) and B cells (target cells) localized within the B cell-rich area (BC aggr, 17), and the surrounding fibroblast-rich area (Fibr I, 16); Commun. prob = communication probability; p-values were calculated in CellChat^94^ using a permutation test.

Integrating the spatial data with a public splenic scRNAseq reference containing germinal-center (GC) B Cell populations^58^ identified transcriptionally distinct light-zone and dark-zone GC B cells, follicular B cells, two memory B Cell populations differing in *Zeb2* expression, and a *Myc*-expressing population associated with light-zone expansion within the thymic aggregates (Fig. 7a, ii-iii). Co-localized expression of *Bcl6*, the master regulator of GC formation, and *Aicda*, encoding the somatic hypermutation and class-switch enzyme activation-induced cytidine deaminase (AID), supported active GC-like reactions within these structures (Fig. 7a iv).^59,60^ Immunohistological analyses further confirmed the presence of CD21+ cells, likely follicular dendritic cells (Fig. 7b), in cytokeratin-free spaces of the B cell aggregates, supporting their organized lymphoid architecture. These aggregates contained CD4+, CD8+, and FoxP3+ T cells, while CD206+ macrophages and CD11c+ dendritic cells were predominantly localized at their periphery (Extended Data Fig. 7c–f). Furthermore, the presence in these aggregates of GL7+ germinal center B cells, CD138+ plasma cells, both IgM+ and IgG+ B cells, CD4+CXCR5+ T follicular helper cells, and PNAd+ high endothelial venules (HEVs) were confirmed by immunohistology supporting the formation of late-stage TLS (Fig. 7c–e, Extended Data Fig. 7g).^61^ Comparable κ/λ light-chain usage in thymus and spleen from the same animals argued against a dominant clonal expansion as the origin of these aggregates (Extended Data Fig. 7h–j).

Together, the cellular composition, molecular signature, and stromal context of these age-associated B Cell aggregates fulfil the defining criteria of tertiary lymphoid structures (TLS) bearing germinal-center-like reactions,^62^ identifying TLS formation as a feature of physiological thymic aging. Because these structures consistently arose adjacent to fibrous structures, we next asked whether the same fibroblast populations directly supply the molecular signals required for TLS formation and whether this population also plays an analogous supportive role for the epithelial progenitors that also accumulate there.

### Medullary fibroblasts form a shared signaling hub that sustains both TLS neogenesis and epithelial progenitors

We next interrogated spatially restricted ligand-receptor signaling within the BCaggr (region 17) and adjacent Fibr I (region 16) niches for factors that regulate B Cell recruitment and epithelial progenitor maintenance. B Cell aggregates and their surrounding areas were enriched for transcripts encoding BAFF (*Tnfsf13b*), APRIL (*Tnfsf13)*, *Cxcl12*, and complement component *C4b* (Fig. 7e), all promoting B Cell recruitment, survival, and differentiation.^63–65^ Mapping these ligands to their cellular source identified medullary fibroblasts (MedFb1, MedFb3) as the principal producers of BAFF, CXCL12, pleiotrophin, midkine and complement components C3/C4b, with cognate receptors expressed on neighboring B cells (Fig. 7f). ItTEC1, ItTEC3, ItTEC4 and TECProg1 contributed additional but quantitatively minor B Cell-supportive signals (Fig. 7f; Extended Data Fig. 7k). These findings position medullary fibroblasts as the principal architects of the molecular niche sustaining local germinal-center-like responses, consistent with the emerging role of fibroblastic stroma in TLS formation and maintenance.^66^

Independently, spatial quantification showed that TEC progenitors accumulated within this same fibroblast-rich territory, with TECProg1 as the predominant epithelial population enriched along the borders of the B Cell aggregates (Fig. 2c i; Fig. 8a; Supplementary Fig. 4c). A spatially-restricted CellChat^67^ analysis of the Fibr I/BCaggr niche identified a communication network dominated by FGF, BMP, WNT, IGF, TGF-β, SEMA3 and midkine (MK) signaling, pathways with established roles in TEC specification, proliferation, differentiation and regeneration (Fig 8b).^68–71^ Several of these ligands (PTN, IGF, TGF-β, SEMA3) were also linked to extracellular matrix remodeling and age-associated epithelial-mesenchymal crosstalk.^72–74^ The dominant signaling axis in this network ran between medullary fibroblasts (MedFb1, MedFb3), TECProg1 and the ItTEC1/ItTEC4 populations (Fig. 8c; Supplementary Fig. 8): fibroblasts supplied the principal trophic input to TECProg1 via PTN, FGF, IGF and TGF-β signaling, while ItTEC populations contributed complementary BMP- and CCL-family signals (Fig. 8d). TECProg1 in turn signaled reciprocally to fibroblasts via MK and SEMA3, to ItTEC via FGF, MK and WNT, and displayed prominent FGF, MK and WNT4 autocrine signaling suggestive of intrinsic reinforcement of its progenitor state (Fig. 8e).

**Fig. 8:**
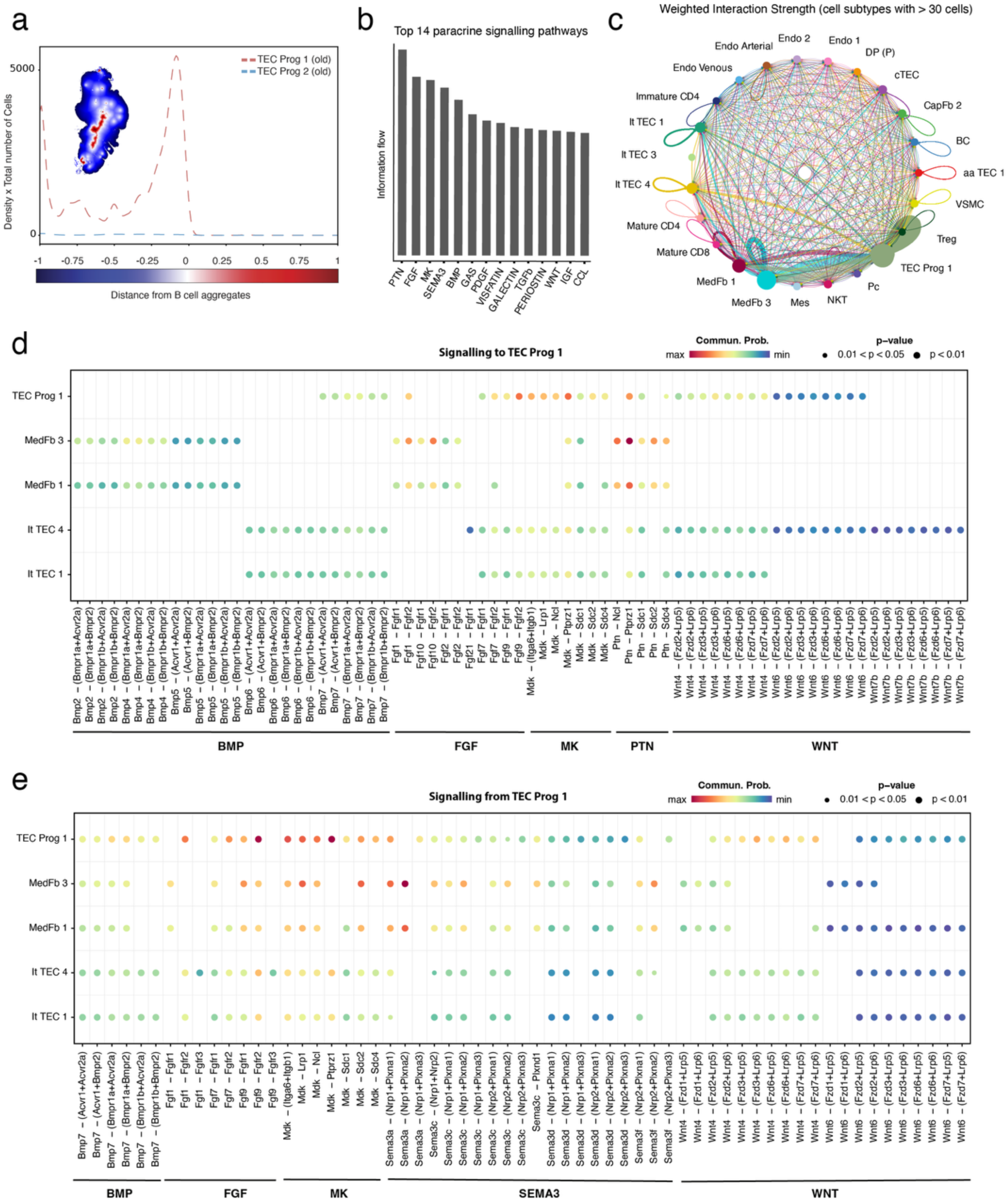
TLS-adjacent stromal cells provide growth signals contributing to the expansion of TEC progenitors. a) Cell count-adjusted probability density functions quantifying the normalized position and abundance of TEC progenitor populations relative to the boundaries of the B cell aggregate region (BC aggr, 17) in the old spatial transcriptomics dataset. b) Information-flow ranked paracrine signaling pathways between all cell types of the fibroblast-rich region (Fibr I, 16). c) Weighted interaction strength network via all available paracrine signaling pathways for cell types within the fibroblast-rich region (Fibr I, 16); only cell types with more than 30 cells were considered; line thickness is proportional to strength of interaction. d-e) Heatmaps of paracrine communication pathways d) to TEC Prog 1 from the stromal cell types indicated on the y-axis and e) from TEC Prog 1 to the stromal cell types indicated on the y-axis. Commun. Prob = communication probability; BMP = bone morphogenetic protein, FGF = fibroblast growth factor, MK = midkine, PTN = pleiotrophin, SEMA3 = semaphorin 3A, WNT = wingless-related integration site; p-values were computed using a permutation test.

Together, these results converge on a single mechanistic theme: the same fibroblast-rich niche that emerges in the aged thymus is predicted to provide trophic signals to the epithelial progenitor compartment while also supplying survival and recruitment cues that support TLS formation. Medullary fibroblasts therefore define a shared signaling hub associated with two of the most striking architectural consequences of thymic aging: the emergence of a spatially restricted epithelial progenitor niche and the neogenesis of tertiary lymphoid structures.

## Discussion

Thymic involution is a principal driver of immune aging, yet the spatial organization of the aging thymic microenvironment and its consequences for thymopoiesis have remained largely undefined. By integrating single-nucleus transcriptomics, chromatin accessibility, CITEseq and single-cell spatial transcriptomics, we generated a spatially resolved multi-omic atlas of the aging thymus that anatomically localizes previously defined stromal populations and reconstructs their communication with developing thymocytes in functionally defined regions. These analyses establish that thymic aging is not simply an epithelial loss, but an extensive spatial reorganization of the cellular niches that support T cell development.

Several age-associated changes identified here, namely the decline of mature mTEC, shifting intertypical TEC abundance, and the expansion of medullary fibroblasts, corroborate previous transcriptomic studies of thymic aging^17,23^, providing orthogonal validation of our approach. Beyond confirmation, this work assigns precise anatomical addresses to populations previously defined largely by their molecular signature, including TEC progenitors, intertypical TEC, mimetic TEC and the newly identified age-associated epithelial states. Spatial context thus constitutes biological information that is not recoverable from dissociated transcriptomic data alone.

A central observation concerns the epithelial progenitor compartment. We localize two transcriptionally distinct TEC progenitor populations to the subcapsular and (cortico-) medullary area and show that TECProg1 expands with age specifically within fibroblast-rich regions adjacent to tertiary lymphoid structures. Together with trajectory inference, this supports a developmental relationship linking TECProg1 to intertypical TEC populations, consistent with descriptions of postnatal thymic epithelial progenitors.^49^ The preferential expansion of ItTEC1 alongside the decline of downstream intertypical populations and mature mTEC further suggests that epithelial differentiation becomes partially arrested during aging, consistent with previous findings.^17^ This inference rests on pseudo-time and replicative-age analyses rather than direct lineage tracing, which will be required to formally establish whether it reflects a genuine developmental block. These experiments are, however, technically not yet possible because suitable cell-surface markers for these populations are still lacking.

A second central finding is that aging displaces the stromal microenvironments in which thymocyte selection occurs, and not merely their cellular composition. During early life, thymocytes migrate along a precisely sequenced series of epithelial niches coordinating positive and negative selection. We show that these checkpoints relocate into different stromal neighborhoods with age, and that this spatial displacement is compounded by a measurable, intrinsic decline in the capacity of aged cortical TEC to mediate positive selection in reaggregate cultures. Hence, both the availability and the functional quality of epithelial support deteriorate with age. Recent transcriptomic studies confirm that thymocyte-intrinsic programs also change with age^75,76^; our functional data indicates that epithelial aging remains a dominant, and tractable, determinant of impaired positive selection, consistent with earlier transplantation and regeneration studies.^77,78^

The expansion of medullary fibroblasts is a further prominent feature of aging thymic stroma. Rather than a passive by-product of fibrosis, these cells establish a signaling-rich microenvironment around TEC progenitors, engaging BMP, FGF, WNT, IGF and MK pathways with established roles in thymic epithelial maintenance and regeneration.^68–71^ Whether this signaling represents a maladaptive response to chronic inflammation or a compensatory attempt to preserve epithelial homeostasis cannot be resolved from our data. The fibroblasts’ coordinated enrichment around TEC progenitors nonetheless indicates that they actively participate in remodeling -rather than simply replacing- the aging epithelial compartment.

We also provide a detailed spatial and molecular characterization of thymic tertiary lymphoid structures arising during physiological aging. Ectopic germinal-center-like reactions in the thymus have previously been described predominantly in context of autoimmune conditions.^27,28,79,80^ The cellular organization, molecular signature and stromal support shown here meet the defining criteria of germinal center B cell-containing TLS, and the enrichment of BAFF, APRIL and CXCL12 in the surrounding fibroblasts and intertypical TEC suggests that stromal remodeling actively contributes to their formation rather than merely accommodating pre-existing lymphoid aggregates.

The functional significance of these thymic TLS remains open. Prior work links analogous structures to both protective local antibody responses^81^ and, in other contexts, to autoreactive B Cell generation.^79^ Our data identifies the stromal architecture capable of sustaining thymic

TLS but does not distinguish between these possibilities. Determining whether these niches compromise central tolerance, support local immune surveillance, or both, is an important direction for future work.

More broadly, these findings support a revised view of thymic involution: rather than uniform epithelial atrophy, aging reorganizes epithelial, mesenchymal and immune compartments into new functional microenvironments that alter thymocyte developmental niches, promote tertiary lymphoid neogenesis, and reroute epithelial-mesenchymal signaling through a common fibroblast hub. Because this hub simultaneously sustains a regenerative epithelial progenitor population and licenses ectopic lymphoid structures, it represents a specific, spatially defined target for interventions aimed at preserving epithelial regeneration while limiting maladaptive lymphoid neogenesis. This distinction will be essential for designing thymic rejuvenation strategies that restore T cell output without exacerbating age-associated autoimmunity.

## Methods

### Mice

C57BL/6 wildtype mice were maintained under specific pathogen-free conditions in the Universities of Oxford and Basel according to their respective local and national regulations or obtained from Charles River Laboratories and rested for at least one week prior to use in experiments. The experimental protocols were approved by the United Kingdom Home Office and the Cantonal Veterinary Office of Basal-Stadt. Young adult (4 to 8-week-old) and aged (65 to 77-week-old) mice were used for the downstream protocols listed below except for the TEC cultures where young animals (2 to 3-week-old) were compared to older animals (6 to 38- week-old). Exact ages, sex and numbers of animals are specified for each section below.

### Thymic stromal cell isolation for flow cytometric comparisons

Thymic stromal cells were isolated from freshly harvested thymic lobes of young (5 to 7-week- old, n=6) and old (75 to 77-week-old, n=6), female C57BL/6 wild-type mice via enzymatic digestion using 75μg/ml liberase TM (Roche) and 300μg/ml DNaseI (Roche) or 500μg/ml Papain (Sigma Aldrich) and 200μg/ml liberase TM (Roche) and 200μg/ml DNaseI (Roche) in a 37°C water bath for a total of 30min, with pipetting at 5-min intervals. Cells were collected into FACS buffer consisting of PBS containing 2% FCS (Sigma Aldrich), filtered through a 100-μm cell strainer and then counted using a hemocytometer. After counting, the cell suspensions were either processed directly for flow cytometry, depleted of CD45+ cells or enriched for EpCAM+ cells using anti-Bio beads and an AutoMACS magnetic cell separator (Miltenyi Biotec).

### Thymocyte isolation for flow cytometric comparisons

Thymocytes were isolated from freshly harvested thymic lobes of young (5 to 7-week-old, n=6) and old (75 to 77-week-old, n=6), female C57BL/6 wild-type mice via mechanical disruption between frosted glass slides. After washing, straining through a 100μm cell strainer and counting, thymocytes were processed for analysis and sorted flow cytometrically.

### Flow cytometry

Cell suspensions of thymic stromal cells or thymocytes were first stained with antibodies directed against cell surface molecules for 30-45 mins at 4°C. For a list of antibodies used see Supplementary Table 1. For assessment of cell viability, DAPI and the fixable viability dyes Zombie Red (Biolegend) and LIVE/DEAD fixable Aqua Dead Cell Stain Kit were used. For panels requiring the detection of intracellular markers, samples were next fixed and permeabilized using the eBioscience FOXP3/Transcription Factor Staining Buffer Set (Invitrogen), before staining for intracellular proteins for 1-16 hours at 4°C. Samples were acquired using a Sony ID7000 spectral cell analyzer or sorted using a BD FACSAria cell sorter. Flow cytometry data were analyzed using FlowJo (v10). GraphPad Prism (v9) was used to perform statistical analyses of flow cytometry data. The statistical tests used are described in the figure legends.

### Spatial transcriptomics

Thymic lobes were harvested from young (8-week-old, n = 1) and aged (65-week-old, n = 1) female C57BL/6 wild-type mice, cleaned of adherent fat, and fixed in 10% neutral buffered formalin (Sigma Aldrich) for 24 hours at 4°C, before ethanol dehydration, clearing with xylene, wax infiltration, and paraffin embedding. FFPE blocks were stored at 4°C for 4 weeks before sectioning at a thickness of 5μm and placement onto a Xenium slide (10x Genomics). After cutting, the slide was incubated at 42°C for 3 hours and then stored in a desiccator at room temperature for 2 weeks before processing for spatial transcriptomics using the Xenium Prime 5k Mouse Pan Tissue and Pathways Panel with the optional Cell Segmentation Staining step (10x Genomics), according to the manufacturer’s instructions. After running on the Xenium Analyser, slides were washed in PBS-T (PBS 0.05% Tween), dehydrated with ethanol, and stored in 30% glycerol (in PBS) at -20°C. The young and the old Xenium datasets were transformed into Seurat objects using Seurat’s LoadXenium function. The objects were filtered for cells with unique feature counts > 50 and < 600 as well as total feature counts < 900.

### Single nucleus RNA and ATAC sequencing data

Total thymic stromal cells (CD45⁻, TER-119⁻) were FACS sorted from young (4 to 6-week-old, n=2, pooled 2 mice per replicate) and aged (68 to 73-week-old, n=2, pooled 2 mice per replicate) female C57BL/6 mice. For a list of antibodies used see Supplementary Table 2. Nuclei were isolated performing an incubation of sorted cells in a 0.1× lysis buffer for 5-mins as per protocol CG000366 (10x Genomics) and then processed for RNA and ATAC sequencing using the Epi Multiome Kit (10x Genomics) and its associated protocol. Libraries were pooled in equimolar concentrations and then sequenced on an Illumina NovaSeq6000. The snRNA and ATAC total stroma sequencing datasets were de-multiplexed and pre-processed using 10x Genomics Cell Ranger ARC (v2.0.0) pipeline and filtered for cells with unique gene counts >1000 and <5500; total gene counts <1.5×10^4; fraction of mitochondrial genes <0.08; and total ATAC reads per cell between 10^3 and 10^4.5. Seurat^82^ (v5.2.0) and Signac^83^ (v1.14.0) were utilized for multimodal integration using a weighted nearest-neighbor approach to obtain a joint RNA/ATAC UMAP projection. Doublets were removed using DoubletFinder (v2.0.4)^84^ assuming a doublet formation rate of 7.5%, 10 principal components (PC), pN = 0.25, and pK = 0.09. Feature plots were generated for the remaining cells, and *Ptprc*-positive clusters were removed. Both resulting age groups were annotated jointly, based on differentially expressed genes in the individual clusters.

### Estimation of mitotic age

Mitotic age was estimated using EpiTrace^50^ (v0.0.2.1), a computational framework that infers the replicative history of individual cells from scATACseq chromatin accessibility profiles by leveraging accessibility at conserved genomic loci. The annotated snRNAseq/scATACseq data was stratified into distinct age groups and subset to relevant cell types. Then, the EpiTraceAge function was applied to the ATAC count matrices using the default “randomized” normalization method prior to mitotic age estimation.

### Pseudo-time trajectories

Pseudo-time trajectories for selected thymocyte subpopulations were inferred using SeuratWrappers (v0.4.0) and monocle3^51^ (v1.4.26). Monocle3 reconstructs cellular trajectories by learning a principal graph within a low-dimensional embedding and orders cells along this graph to estimate their progression through a continuous biological process (pseudo-time). The literature-adapted thymocyte reference dataset^52^ was pre-processed as described above. To orient the trajectory according to thymocyte maturation, the root node was defined within the signaling double-positive T cell (DP Sig.) population, from which pseudo-time values were assigned across the learned trajectory.

### Cross-modal integration of single-cell and spatial transcriptomics data

An annotated CITEseq dataset for intrathymic hematopoietic cells was adapted from literature^52^ and linked to its full transcriptome (NCBI Gene Expression Omnibus (GEO) accession number GSE186078). The original dataset was subsetted to the 5 available wild-type C57BL/6 samples and the raw counts were batch corrected using ComBat-seq^85^ (sva package v3.54.0) while maintaining the original annotations (Extended Data Fig. 2.1e-f). This was analogously conducted for the aforementioned thymic stromal cell snRNAseq dataset. The young and old spatial transcriptomic datasets were then split into two cell classes, namely hematopoietic and stromal-like cells, based on the overlap with differentially expressed genes in their respective, CITEseq or snRNAseq reference datasets. Specifically, differential gene expression between the stromal (snRNAseq) and hematopoietic (CITEseq) reference datasets was computed using Seurat’s FindMarkers function with “min.pct” option = 0.2 and collection of up to the top 50 DEGs for each cell type within each dataset. The resulting DEG-lists were depleted of genes occurring in both lists to assure uniqueness. This allowed the assignment of cell-class (i.e. hematopoietic or stromal-like) to each Xenium cell by querying the cell’s transcriptome for overlap with markers present in either DEG-list using a majority voting scheme (Extended Data Fig. 2.1g). The reference cells profiled via snRNAseq/CITEseq and the query cells profiled by spatial transcriptomics were then separated into their respective ages as well as stromal and hematopoietic cell class categories. The spatial transcriptomics data was normalized in these groups using SCTransform^82^ and integrated against the respective reference dataset. Here, the same CITEseq reference dataset was utilized for both the young and the old tissue section. In Seurat, a canonical correlation analysis (cca) was selected as integration method for the stromal cells while, due to large cell counts, the less computationally demanding reciprocal principal component analysis (rpca) was chosen for the hematopoietic cells. This resulted in joint UMAP representations of both modalities for stromal and hematopoietic cells. Both integrations were performed using Seurat’s sequence of SelectIntegrationFeatures, PrepSCTIntegration, FindIntegrationAnchors and IntegrateData. The integrated datasets of both ages and cell class categories were scaled and subjected to a PCA. Subsequently, for the stromal cells only, the integrated UMAP projection was restricted to only those cells that were highly similar across both Xenium and snRNA/ATACseq using the spectral clustering package DBSCAN (to remove highly dissimilar cells) and the RANN (v2.6.2) package’s nn2 function (to identify nearest neighbors) as well as Seurat’s label transfer annotation score (used as an additional quality metric to discard dissimilar cells in the query dataset). Next, for the remaining stromal and hematopoietic datasets (here illustrated for the hematopoietic cell category of the young dataset in Extended Data Fig. 2.1h), the scaled, integrated transcription profiles of the 3000 most variable genes in each cell were correlated across the two modalities, identifying each reference cell’s most similar spatial transcriptomics counterpart (here demonstrated for the hematopoietic cell category in Extended Data Fig. 2.1i-j). In both cases, the highest-correlating quartile of query cells was assigned molecular information from their single most similar reference cell, including, the full transcriptome, snATACseq accessibility profile or CITEseq protein measurements. The transfer of molecular information from one reference cell to multiple query cells was permitted. In addition to molecular information, previously curated cell type annotations were directly propagated to the top correlated quartile of cells in the same fashion (Extended Data Fig. 2.1k and Supplementary Fig. 2.2a). A differential gene expression analysis for both the hematopoietic and stromal cell category was conducted to quantify the quality of the cross-modal link. This DGE analysis was performed separately in the snRNAseq resp. CITEseq, and spatial transcriptomics datasets using Seurat’s FindMarkers function with logfc.threshold=0.25 and min.pct=0.1 to compare resulting DEGs for identical cell types with more than 30 cells across both modalities (Extended Data Fig. 2.1c-d). Additional quality control included querying the location of Xenium-absent marker gene expression of cell populations with a well-defined spatial location after imputation (Supplementary Fig. 2.2 a,b), as well as a global analysis that confirmed the expected, spatial niche enrichment of cells with imputed cell labels (Fig. 4a, Supplementary Fig. 4c). The remaining cells, namely the lower 3 quartiles that did not show sufficiently high correlation between the reference and the query datasets, were subjected to an additional round of annotation relying on transcriptomic profiles of the spatial transcriptomics data alone. The previously defined annotations assigned by label transfer from the references datasets as described above were used as qualitative anchors to inform the broader annotations of the remaining cells (here illustrated for the stromal cell category (Extended Data Fig. 2.2a-b)). Cells with antigen presenting character such as B cells, dendritic cells and other APCs which initially were assigned to both cell categories received a joint spatial transcriptomics-based annotation to unify their annotation (Extended Data Fig. 2.2c-e). The spatial projection of this additional layer of annotation (i.e. Xenium-level annotation) along with the combined projection of both annotation layers is visible from Supplementary Fig. 2.2b-c.

### Spatial analysis

Spatial analyses of spatial transcriptomic data were performed in Python using MuSpAn (v1.2.4).^56^ Raw spatial transcriptomics outputs along with previously defined cell type annotation layers were converted into MuSpAn domain objects characterized by each cell’s centroid location. The exterior boundary of each tissue domain was estimated using an alpha- shape reconstruction with a radius parameter of 150μm. Medullary regions were further partitioned using a hexagonal lattice with a side length of 50μm to facilitate tissue scale analysis. In addition, a spatial graph was constructed by applying Delaunay triangulation to cell centroids, with edges longer than 20μm removed to restrict connectivity to local cellular interactions.

### Neighborhood analysis

To identify recurrent functional tissue microenvironments, neighborhood cluster analysis was performed using cell type annotations of the transcription data (Supplementary Fig. 2.2b). Following existing workflows, a local cellular composition vector was computed from the proportions of each annotated cell type within its three-hop neighborhood (corresponding to an effective spatial extent of approximately 60μm) for each cell. These neighborhood composition vectors were clustered using k-means clustering, with the number of clusters (k=17) selected by silhouette analysis.^56,86^ The resulting neighborhood clusters were manually annotated according to their characteristic cellular compositions and assigned as cell-level labels within the MuSpAn domain objects using the cluster_neighbourhoods() function. Spatial adjacency of neighborhoods was quantified from the underlying spatial graph by identifying graph edges interfacing distinct neighborhood clusters (Supplementary Fig. 4a). For each neighborhood cluster i, the connectivity to cluster j was defined as the proportion of all interfacial edges originating from i that terminated in j, such that Cᵢ→ⱼ = Eᵢⱼ / Σₖ Eᵢₖ where Eᵢⱼ denotes the number of graph edges connecting cells assigned to neighborhood clusters i and j. This normalization yields a weighted, directed graph describing the relative connectivity between neighborhood regions.

To characterize the local thymocyte microenvironments surrounding cortical thymic epithelial cells (cTEC), a second neighborhood clustering analysis was performed using cTEC as the focal cells. In contrast to the graph-based neighborhood definition described above, fixed-radius neighborhoods were defined, with local composition calculated as the proportions of thymocyte populations within 30μm of each cTEC centroid. This distance-based definition was adopted because the morphology of cTEC is not well represented by centroid-derived graph connectivity. The resulting neighborhood composition vectors were clustered using k-means clustering, with the number of clusters (k=6) selected by silhouette analysis. The resulting neighborhood classes were assigned as cell-level annotations using the cluster_neighbourhoods function in MuSpAn.^56^

### Local Getis-Ord Gi* statistic

Spatial clustering of mimetic cell populations within medullary hexagonal lattice regions was quantified using the local Getis–Ord Gi* statistic. For each hexagon, the Gi* statistic compares the weighted sum of mimetic cell counts within its neighborhood, defined by region adjacency, with the value expected under complete spatial randomness, returning a standardized* *z- score.*^87^ Positive *z*-scores indicate local enrichment (hotspots), whereas negative *z*-scores indicate local depletion (coldspots). Analysis was performed for medullary regions exceeding an area of (1 × 10⁵) μm². The number of mimetic cells within each hexagon was used as the input variable, and spatial neighborhoods were defined by direct adjacency between neighboring hexagons. Local Gi* statistics were computed using the getis_ord() function in MuSpAn.^56^

### Quadrat correlation analysis

Spatial co-occurrence between mimetic cell subtypes was quantified using quadrat correlation analysis. This method quantifies the correlation in abundance of pairs of labelled cell populations across a set of spatial regions and compares the observed correlation with a null distribution generated by random permutations of cell labels, yielding a standardized z-score for each pairwise comparison.^88^ Positive z-scores indicate spatial co-enrichment of the corresponding cell populations, whereas negative z-scores indicate spatial segregation. The analysis was performed using the medullary hexagonal lattice as the set of quadrats, with mimetic cell subtype as the cell label. Pairwise quadrat correlation matrices were computed using the quadrat_correlation_matrix() function in MuSpAn with 1,000 cell type label permutations.

### Pair Correlation Function

To assess spatial association of mimetic subpopulations across multiple length scales, we computed the Pair Correlation Function (PCF) for each mimetic cell type. The pair correlation function (PCF), *g(r)*, quantifies the density of point pairs separated by distance *r* relative to the density expected at the same distance for a given spatial null model.^56^ Here we consider the null of complete spatial randomness as a practical and interpretable baseline for assessing structure in biological data.^89,90^ In this reference, values of g(r) > 1 indicate enrichment or co- localization at distance r, whereas values of g(r) < 1 indicate spatial exclusion. PCFs were calculated within medullary regions exceeding an area of (1 × 10⁵) μm². Calculations used an annulus width of 20μm and a maximum distance of 250μm, with automatic boundary correction supported. Mimetic PCFs were estimated using the cross_pair_correlation() function in MuSpAn.^56^

### Normalized corticomedullary junction distance

The Euclidean distance of each cell from the corticomedullary junction (CMJ) was calculated in the young and old spatial transcriptomics dataset. For cells located in the cortex, the Euclidean distance to the capsule boundary was additionally determined. To enable comparison between the cortical and medullary compartments despite differences in their widths, distances were normalized independently within each compartment. Cortical distances were normalized by dividing the distance from the CMJ by the sum of the distances from the CMJ and the capsule. Medullary distances were normalized by the maximum distance from the CMJ within the medullary region. Consequently, the normalized distance ranged from 0 at the CMJ to 1 in the cortex and −1 in the medulla, corresponding to the deepest point within each compartment. These calculations were performed using the get_minimum_distances_boundaries() function in MuSpAn.^56^

### Probability Density Functions

To assess the distribution of the TEC subtypes relative to the CMJ, probability density functions were estimated for each cell type separately across both ages of the spatial transcriptomics experiment. This was achieved using the gaussian_kde function from SciPy’s statistics module.^91^ The densities were then scaled by the number of cells of each cell type in the relevant tissue sample.

### Gene module score estimation

Gene module scores for distinct biological processes were estimated using Seurat’s AddModuleScore function. For each cell, the score was calculated as the average expression of target genes in a predefined module relative to expression-matched control genes. The following gene modules were queried against expressed genes in indicated cell types for each of the two methods investigated. The following gene modules were employed: Senescence: *Cdkn1a, Cdkn2a, Trp53, Rb1, Rbl1, Rbl2*; Inflammaging: *Il1a, Il1b, Il3, Il6, Cxcl1, Cxcl2, Il18, Ccl2, Tnf, Ifng, Cxcl1, Cxcl12*; Senescence-associated secretory pathways (SASP): *Il13, Il15, Cxcl1, Cxcl2, Cxcl3, Ccl8, Ccl2, Ccl7, Ccl3, Ccl20, Ccl16, Ccl26, Ccl17, Cxcl11, Cxcl5, Ccl1, Tgfb2, Tgfb3, Csf2, Csf3, Ifng, Mif, Areg, Ereg, Egf, Fgf2, Hgf, Fgf7, Vegfa, Ang, Kitlg, Cxcl12, Plgf, Ngf, Igfbp2, Igfbp3, Igfbp4, Igfbp6, Igfbp7, Mmp1a, Mmp3, Mmp10, Mmp12, Mmp13, Mmp14, Timp1, Timp2, Serpine1, Serpine2, Plat, Plau, Ctsb, Icam1, Icam3, Tnfrsf11b, Tnfrsf1a, Tnfrsf1b, Tnfrsf10c, Fas, Plaur, Il6st, Egfr, Ptgs2, Nos2, Nos3, Nox4, Cyba, Cybb*. Epithelial-mesenchymal transition (EMT): *Tgfb1, Fgf2, Snai1, Mmp3, Mmp1a, Mmp2, Fn1, Col1a1, Col1a2, Col3a1, S100a4, Vim, Cdh2, Acta2, Snai2, Twist1, Twist2, Zeb1, Zeb2.*^29–32^

### Kernel Density Maps

Spatial distribution of gene expression and gene module scores were visualized as two-dimensional kernel density maps using ggplot’s (v4.0.2) stat_density_2d function. This function performs a two-dimensional kernel density estimation, generating smoothed density contours that highlight regions of locally enriched signal. Prior to visualization, expression floors were applied to exclude low-level background expression.

### Thymic epithelial cell cultures

Thymi were isolated from young (2–3 weeks old) and old (6–38 weeks old) female wild-type C57BL/6 and Cre-control mice. For the mixed-age co-culture experiments, thymi from 15 young mice (2–3 weeks old; average age: 17 days) and 26 old mice (6–26 weeks old; average age: 15.6 weeks) were used. For the age-matched co-culture experiments, thymi from 12 young mice (2–3 weeks old; mean age: 15.5 days) and 28 old mice (11–38 weeks old; mean age: 24 weeks) were used. cTEC (EpCAM+Ly51+UEA-) high and low for MHCII and pre-selection double-positive thymocytes (CD4+CD8+CD69-CD5-) were co-cultured and the induction of CD69 and CD5 was assessed (antibodies specified in Supplementary Table 3). 40k TEC and thymocytes were mixed at a 1:1 ratio in 1.5mL Eppendorf tubes, centrifuged for 1 minute at 6000 rpm, and incubated in IMDM supplemented with 10% fetal calf serum (FCS). The tubes were incubated with their lids left open for 48 hours at 37°C in a humidified incubator maintained at 10% CO_2_.

### Multiplex spatial proteomic imaging using the PhenoCycler Fusion platform

Thymic lobes were harvested from young (4-week-old, n=2) and aged (75 to 77-week-old, n=5) female C57BL/6 wild-type mice, cleaned of adherent fat, embedded in OCT compound (TissueTek), and frozen in a slurry of dry ice and 2-Methylbutan. The blocks were stored at - 80°C before cryosectioning at a thickness of 7μm, and collected on slides, before processing according to the PhenoCycler Fusion User Guide, v2.1.0 (Quanterix). Each antibody (Supplementary Table 4) was either purchased pre-conjugated, or custom-conjugated using a conjugation kit (Quanterix). Complementary oligonucleotides attached to one of three separate, non-overlapping fluorochromes (reporters) were used to detect antibody binding to their respective epitopes in a sequential manner.^92^ During each staining cycle, an image was recorded using the PhenoImagerFusion slide scanner (Quanterix). Images were preprocessed and combined after cycling using the PhenoImagerFusion software (Quanterix). The experiment produced an image stack at 0.5μm per pixel resolution which was processed using a custom MATLAB (R2024b) pipeline. After capsular artefact removal, a background subtraction was performed on selected markers using a rolling ball radius of 40 pixels before performing a global illumination correction by normalizing the fluorescence intensity of each sample to the global mean and standard deviation. Next, a round-cell segmentation mask was computed to identify hematopoietic cells based on DAPI and CD45 using Cellpose.^93^ Outliers were removed, and average fluorescence intensities normalized between 0 and 1. Next, gating approaches were used to annotate cell types (Supplementary Table 5). This led to the identification of joint cell types across both age groups. Cell-cell adjacencies were computed for each phenotype pair and for each tissue sample. Dendritic cells and thymocyte centroids within <7μm distance were considered adjacent, and those instances were collected in a matrix. Phenotype permutation and Z-score normalization was conducted on a per-sample basis to assess whether juxtaposition of cell types occurred at frequencies above chance. Positive z-scores indicate phenotype pairs that occur more frequently than expected by chance (enriched interactions). To identify age-dependent differences in spatial associations, the z-scores for each phenotype pair were compared across the ages using an exact two-sided permutation test, which is a non-parametric test that does not rely on the normality assumption.

### Single-cell and spatial transcriptomics-based profiling of B cell aggregates

The spatial transcriptomics data was pre-processed as outlined above and restricted to the B cell-rich area depicted in Fig. 7ai, guided by spatial *Cd19* expression. The cells within this area were then subset for those that have non-zero expression of *Cd19, Cd83, Cd38* and *Bcl6* to enrich for B cells. Next, we adapted a scRNAseq reference dataset^58^ (NCBI Gene Expression Omnibus (GEO) accession number GSE148805) that was obtained from > 6-week-old, immunized mouse spleens. The raw counts were normalized, clustered using Seurat’s standard clustering workflow and re-annotated employing a similar strategy to the one reported by the original authors. This reference dataset was integrated against the B cell-enriched, spatial transcriptomics query region using an integration workflow analogously to the one described above (Supplementary Fig. 7a-d).

### Immunofluorescence-based profiling of B cell aggregates

Frozen thymus tissue sections (7μm) from 72 to 77-week-old female C57BL/6 mice were air-dried overnight at room temperature, fixed in acetone for 10 min, rehydrated in PBS for 10 min, and blocked with 5% goat serum in PBS for 20 min. Sections were then incubated in blocking buffer with antibodies outlined in Supplementary Table 6. Following PBS washes, sections were counterstained with DAPI (Merck), mounted with Hydromount (National Diagnostics), and imaged using a Leica DMi8 microscope. Image analysis was performed using the OMERO microscopy data management system.

### Ligand:receptor analysis

Cellchat(v1.6.1)^67,94^ was used for the spatially-restricted ligand receptor analysis. The 65- week-old thymus spatial transcriptomics dataset was utilized with imputed full transcriptomic information following the workflow described above. For the B cell analyses, this dataset was subset for the Fibr I and BC aggr neighborhood clusters (Fig. 4a ii), while for crosstalk within the fibrous area only Fibr I was considered. CellChat objects were generated using a conversion factor of 1 and a spatial tolerance of 3μm, corresponding to the approximate radius of a mouse T cell. Both spot.diameter and spot scale factors were set to 10μm. Cell types represented by fewer than 30 cells were excluded prior to communication probability inference, which was performed using a trimmed mean (trim = 0.1), a distance scaling factor of 0.08, and the “Secreted Signaling” interaction database. Expression values were normalized using the “truncatedMean” method.

## Supporting information

Supplementary Data DGE

## Supplementary information

This article is complemented by a set of supplementary files:

- Supplementary_Figures.pdf
- Supplementary_Tables.pdf
- Supplementary_Data_DGE.csv

## Acknowledgments

The authors would like to thank the flow cytometry facilities at the Institute of Developmental and Regenerative Medicine (IDRM), and at the Department of Biomedicine (DBM); the genomics facility at the Department of Biosystems Science and Engineering (D-BSSE); and the sciCORE scientific computing center at the University of Basel for their support. The authors also thank Joanna Hester for her assistance with the experimental acquisition of the spatial transcriptomics datasets. This research was supported by Swiss National Science Foundation grants IZLJZ3_171050 and 310030_184672 (G.A.H.); Medical Research Council grant MR/S036407/1 (G.A.H.); and Wellcome Trust grants 105045/Z/14/Z and 211944/Z/18/Z (G.A.H.). A.T. holds a PhD student fellowship from the Biowise Foundation in Lausanne, Switzerland. F.D. was supported by an NIHR Clinical Lectureship. J.M. was supported by the Cancer Research UK (CRUK) grant CTRQQR-2021/100002, through the Cancer Research UK Oxford Centre. A.E.H. received funding from the MRC grant MR/X022013/1, the NIHR Oxford Health Biomedical Research Centre, and MyAware.

## Ethics declarations Competing interests

The authors declare no competing interests.

## Data availability

The single cell sequencing data supporting this study was deposited in the gene expression omnibus (GEO) in the super series GSE314651. The snRNA/ATACseq data can be found under GSE314650 and the spatial transcriptomics data is deposited under GSE314649.

## Code availability

The codes utilized in the molecular analysis of this project are accessible on Github: https://github.com/andy7132/Thymus-Aging.

## Author contribution

GH, AT, FD designed the study. FD, SZ, AnKu, SM, MD conducted the mouse work. FD, AnKu, SM, MD conducted the snRNA/ATACseq experiments. FD and AT conducted the spatial transcriptomics experiments. AnKu conducted the thymic epithelial cell cultures. SZ conducted the immunohistochemical experiments. FD, AnKu, AmKh analyzed flow cytometric data. AT conducted the clustering and cell type annotation of single cell and spatial transcriptomic data under supervision of GH, FD, SZ, TB. AT designed and implemented the cross-modal integration workflow under supervision of GH and AH. AT conducted molecular analyses involving the single-cell and spatial transcriptomics data under supervision of GH and FD. JM, JG and AT conducted spatial analyses. LT, AT and SZ conducted computational histology analyses. GH, HB, FD, AH supervised computational analyses. AT, GH, FD interpreted the results. AT, GH, FD, prepared figures and wrote the manuscript with input from all authors.

## Extended Data Figures

**Extended Data Fig. 1:**
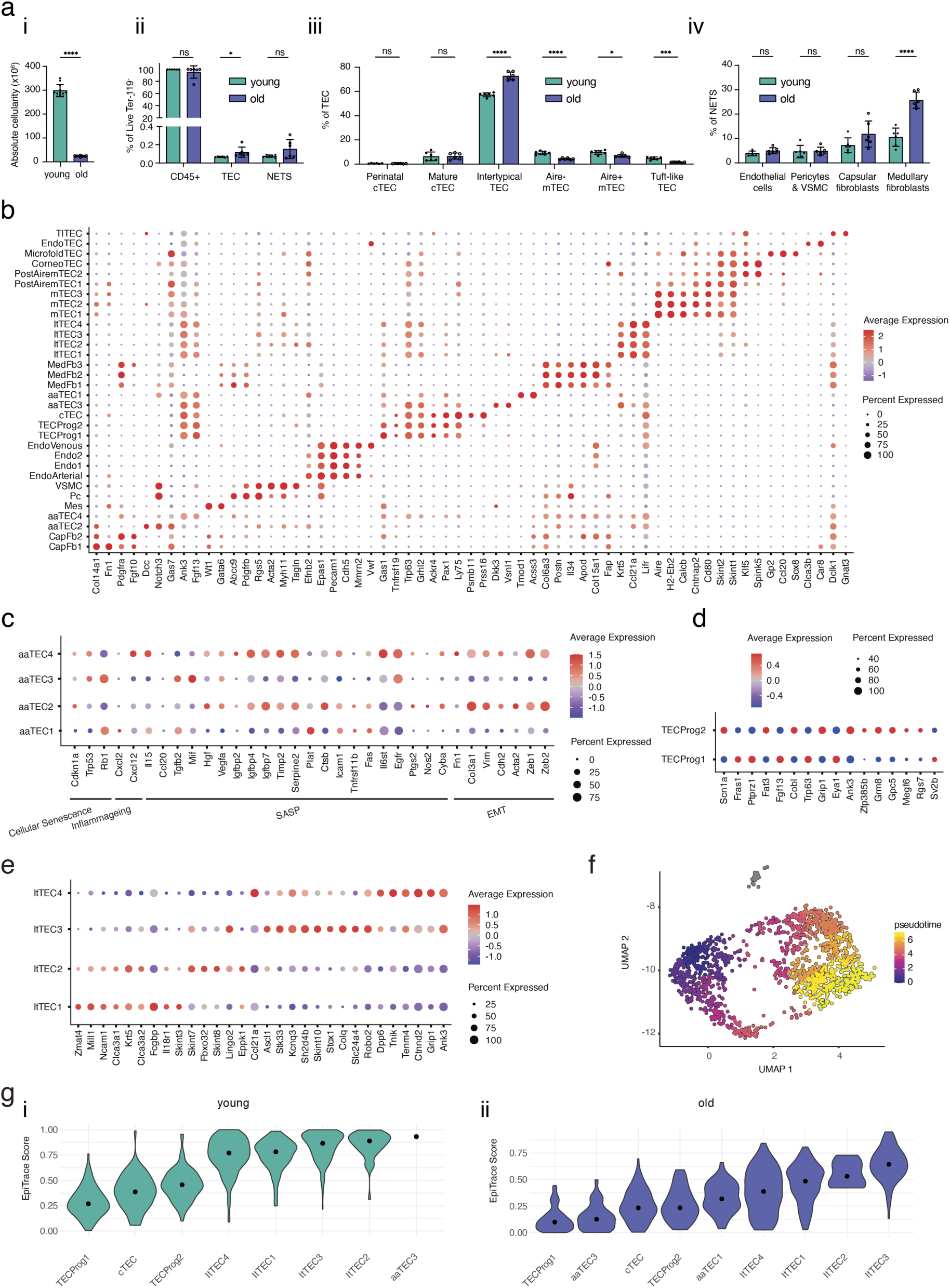
Molecular signatures and age-related changes in frequencies of stromal cells. a) Young and old mice thymi were analyzed and compared for (i) absolute thymic cellularity after enzymatic digestion (n = 12 per group from 4 independent experiments) and (ii-vi) their composition as determined by flow cytometry; n = 6 per group from 2 independent experiments. Gating strategies are outlined in Supplementary Fig. 1.1 and 1.2. (ii) Relative contribution of CD45+ cells, TEC (CD45-EpCAM+) and NETS (CD45-EpCAM-), with prior gating on live, Ter-119-, positive cells. The relative composition of the thymic stroma is shown in (iii) for TEC and (vi) for non-epithelial thymic stroma; Data is represented as mean +/- standard deviation. Statistical analyses were performed using a two-tailed, unpaired student’s t-test for singular comparisons, and multiple unpaired t-tests with Holm-Sidak correction for multiple comparisons. Abbreviations: TEC = thymic epithelial cells; NETS = non-epithelial thymic stroma; cTEC = cortical thymic epithelial cells; mTEC = medullary thymic epithelial cells; VSMC = vascular smooth muscle cells. b) Differentially expressed marker genes for thymic stromal cell populations profiled via snRNA/ATACseq (extended lists are provided in Supplementary Data DGE). c) Gene expression of age-associated (aa) TEC populations for aging-related processes such as cellular senescence, inflammaging, senescence-associated secretory pathways (SASP) and epithelial to mesenchymal transition (EMT).^29–32^ d) Differentially expressed genes in TEC progenitor populations defined against the full stromal cell compartment. e) Differentially expressed genes of distinct intertypical TEC subtypes. f) Joint monocle3-based^51^ pseudo-time trajectory estimation for both ages of the snRNA/ATACseq dataset profiling the early TEC compartment. g) EpiTrace-enabled^50^ estimation of mitotic age score for distinct TEC populations in the young and old snRNA/ATACseq samples.

**Extended Data Fig. 2.1:**
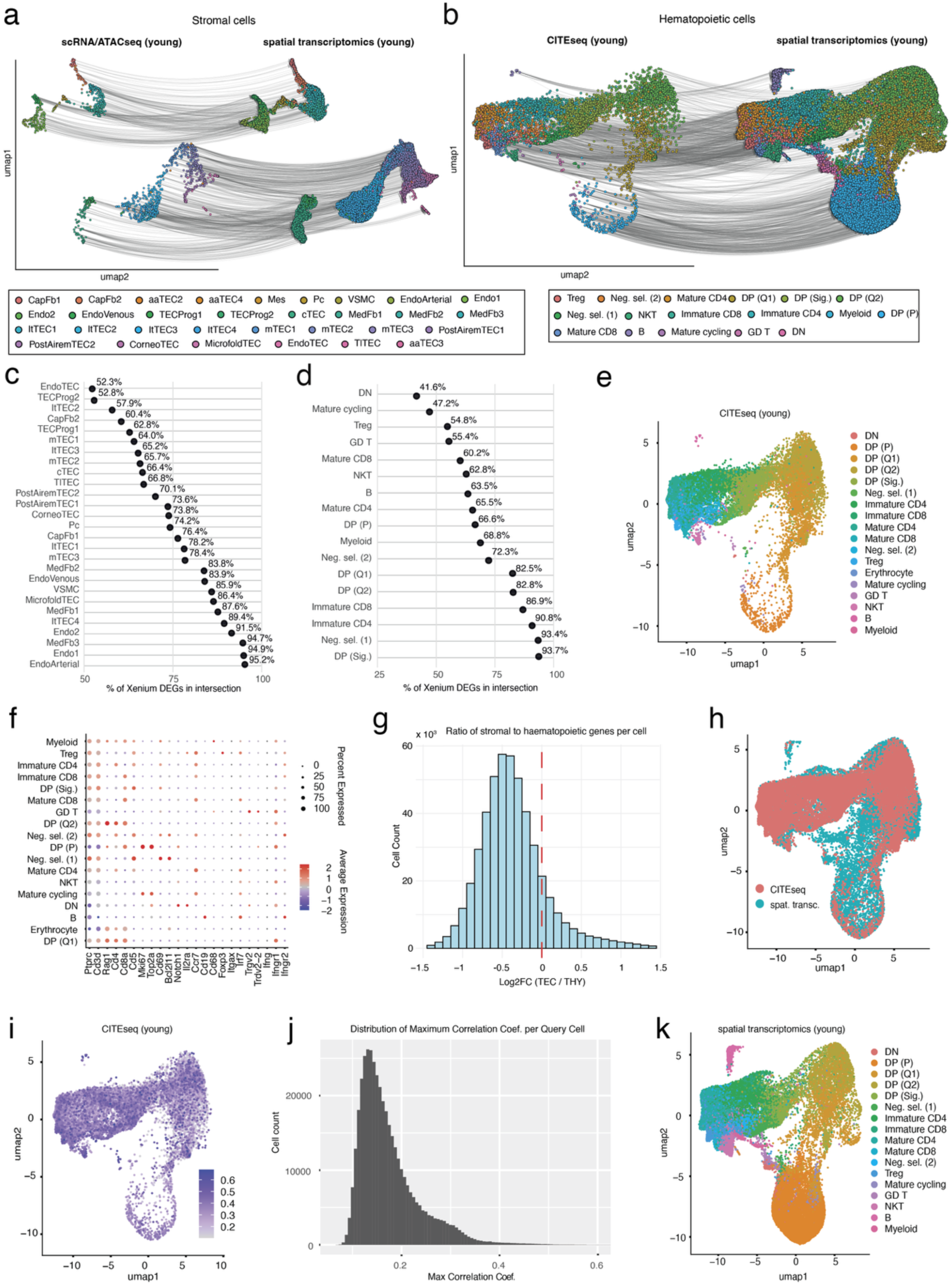
Cross-modal integration and annotation workflow to combine single-cell and spatial transcriptomic data. Sub-sampled spring graph illustrating the propagation of cell type annotations and molecular information between highly similar cells of the a) snRNA/ATACseq, thymic stromal data and b) the adapted CITEseq dataset of thymic hematopoietic cells, and the spatial transcriptomics dataset respectively. c-d) Quantification of shared differentially expressed genes (DEGs) between stromal and hematopoietic cell types. Shown is the cell type-specific proportion of DEGs identified in the spatial transcriptomics dataset that overlap with DEGs detected in differential expression analyses of the corresponding cell types in the dissociated single- cell datasets; Represented are only those cell types with more than 30 cells across both modalities. e) UMAP- projection of the adapted^52^, hematopoietic CITEseq reference dataset along with original annotations. f) Characteristic gene expression signatures for selected genes in the adapted, hematopoietic CITEseq reference dataset. g) Histogram illustrating the initial categorization of cells in the spatial transcriptomics data according to the relative proportion of stromal to hematopoietic genes. Cells with more stromal than hematopoietic genes were assigned to the stromal cell category while all other cells were assigned to the hematopoietic cell category. h) Integrated UMAP projection of the adapted, hematopoietic CITEseq reference dataset and the sub-sampled, hematopoietic-cell-restricted spatial transcriptomics dataset. i) UMAP projection of the hematopoietic CITEseq reference dataset; the color gradient represents the correlation coefficient that is exhibited to cells in the spatial transcriptomics dataset. j) Histogram of recorded correlation coefficient values between expression profiles of hematopoietic cells in the spatial transcriptomic dataset exhibited towards cells in the CITEseq reference dataset. k) Sub-sampled UMAP projection of the hematopoietic spatial transcriptomics dataset along with cell type annotation after label propagation from the CITEseq reference dataset.

**Extended Data Fig. 2.2:**
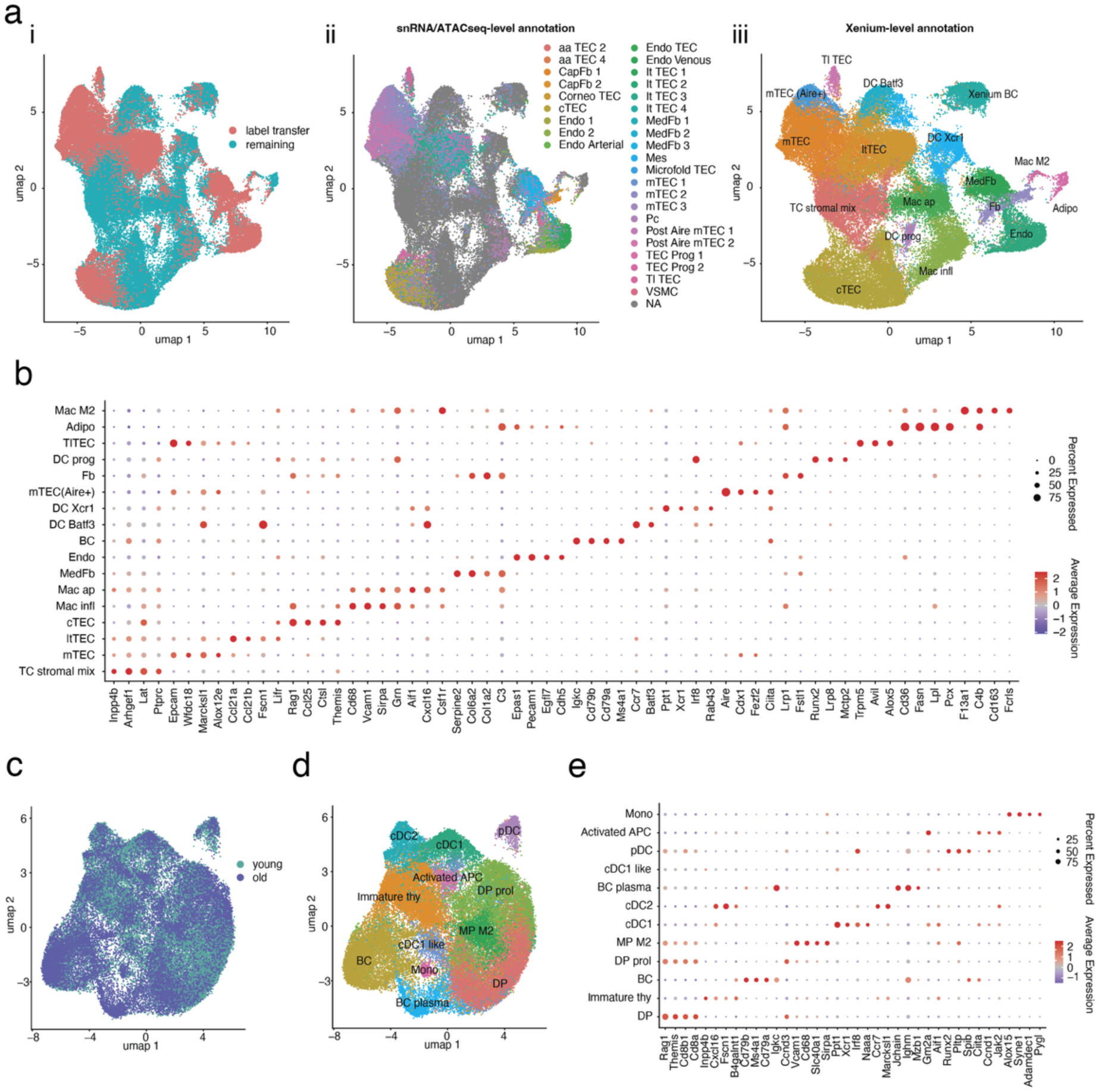
Spatial transcriptomic-based annotation of stromal and hematopoietic cells. a) Annotation workflow employed for the non-reference-annotated, “remaining” stromal cells across both ages. The reference-annotated cells were used as qualitative annotation anchors to facilitate the annotation of the remaining cells. (i) Sub-sampled UMAP projection of the distinct annotation strategies employed for spatial transcriptomic data (ii) UMAP projection of the spatial transcriptomic data of label-transfer-based snRNA/ATACseq-level annotations and (iii) the resulting spatial transcriptomics-level annotations. b) Characteristic marker gene expression of the resulting spatial transcriptomics (or Xenium) -level annotations of the stromal cells. c) Integrated UMAP projection of shared cell types between the stromal and hematopoietic cell categories for the age-pooled spatial transcriptomics datasets. d) Spatial transcriptomics-informed annotation of shared cell types for the age- pooled datasets. e) Characteristic gene expression signatures for spatial transcriptomics-informed annotation of the shared cell types.

**Extended Data Fig. 5:**
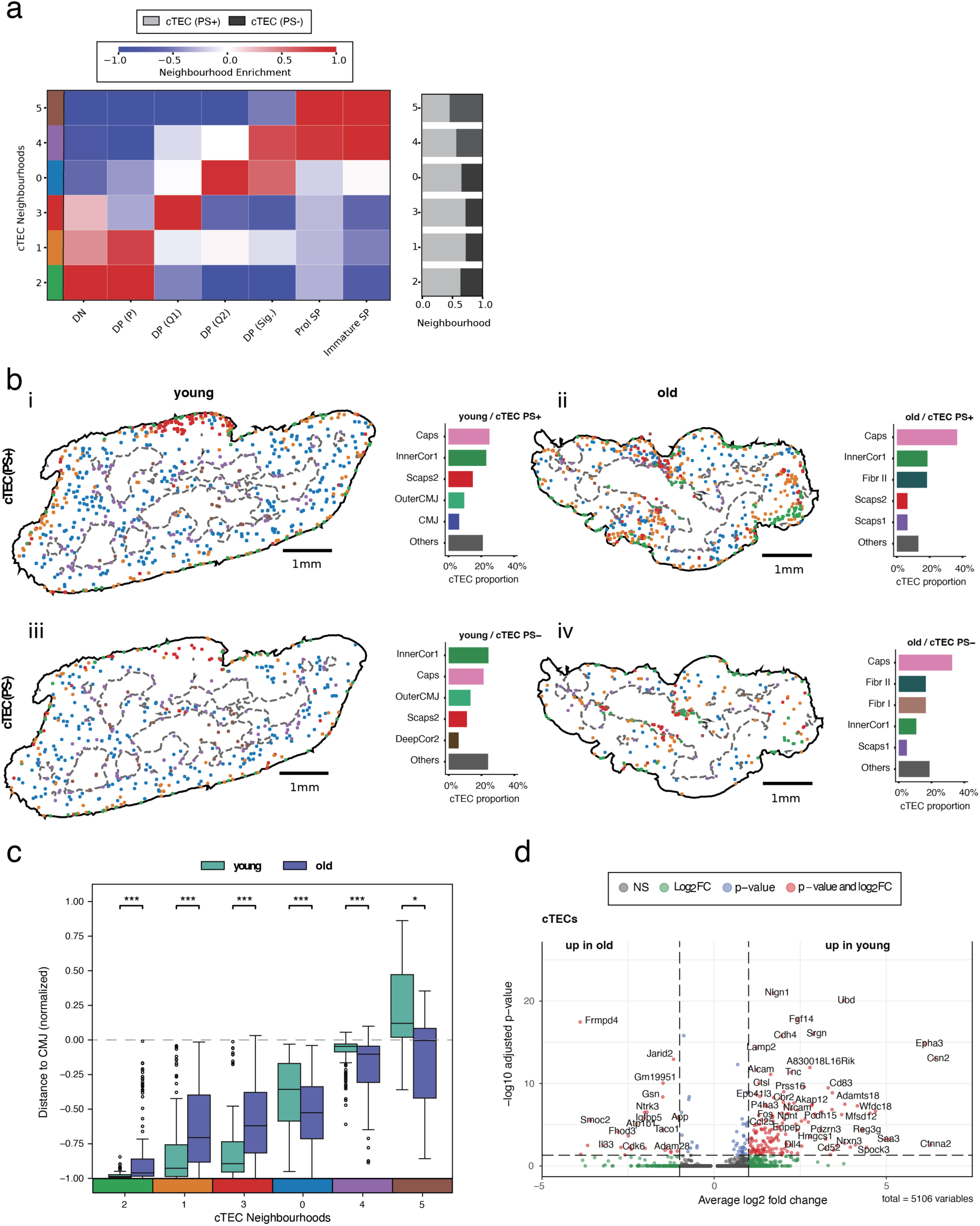
Aging reshapes the molecular identity and spatial distribution of cTEC in the thymus. a) Heatmap representing the local neighborhood enrichment (k=6) of T cell subtypes in proximity of cTEC of both age groups in the spatial transcriptomics data; Horizontal bar charts show the relative abundance of each neighborhood by age; PS+ denotes perinatal cTEC^57^, expressing markers including *Cd83, Cd274, Cd40*, *Tnfrsf14*, and *Icam1*, whereas PS- comprises all other cTEC. b) Spatial distribution of distinct cTEC across the young and old spatial transcriptomic dataset. Cell centroids are colored by their local neighborhood identity; Horizontal bar charts quantify the cluster membership of young and old cTEC subtypes according to the global neighborhood clustering (performed in Fig. 4a). c) Box plot quantifying the normalized spatial distribution of individual cTEC neighborhoods with respect to the CMJ in the young and old spatial transcriptomics data; a two- sided Mann-Whitney U test was employed to compare the resulting distributions with * p ≤0.05, ** p ≤0.01, *** p ≤0.001. d) Volcano plot showing differential gene expression between young and old cTEC of the snRNA/ATACseq data.

**Extended Data Fig. 7:**
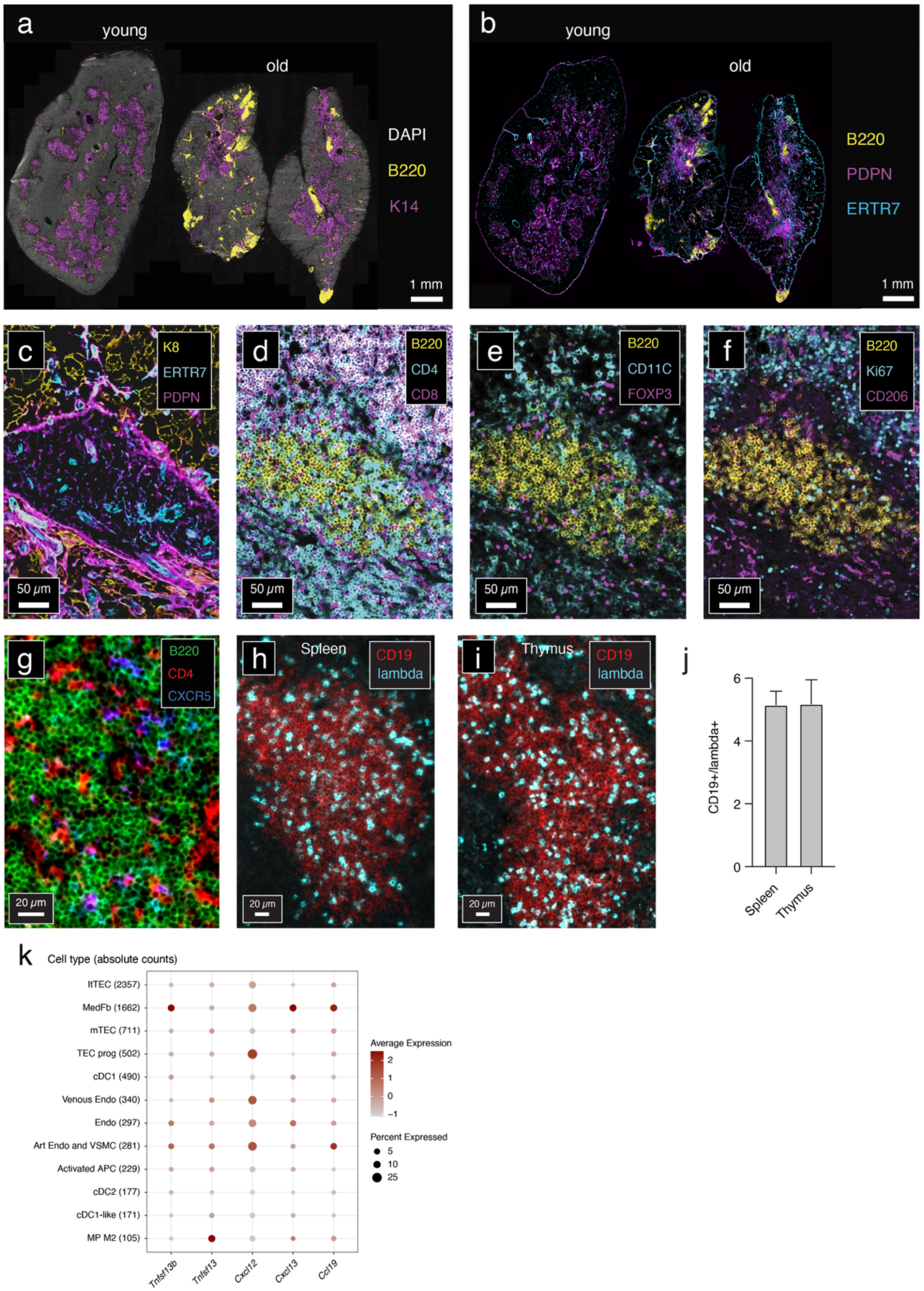
Spatial distribution and molecular characteristics of thymic TLS. a,b) Representative immunofluorescence images of young and old mouse thymic tissue sections showing the localization of cytokeratin 14 (K14)-positive medullary thymic epithelial cells (mTEC), podoplanin (PDPN)- and ERTR7-positive stromal cells, and B220-positive B cells. c-f) Multiplex immunofluorescence images of a representative thymic B cell aggregate in an old mouse thymus showing the localization of B220 expressing B cells alongside thymic stroma and thymocytes positive for CD4 and/or CD8, CD11c+ cells, FOXP3+ regulatory T cells; and CD206+ cells; images are representative of 2 independent experiments. g) Immunofluorescence image of an aged thymic B Cell aggregate stained for B220 (green), CD4 (red), and CXCR5 (blue); the image is representative of 2 independent experiments. (h,i) Immunofluorescence images of CD19 and λ light chain staining representing a splenic and a thymic B cell aggregate of the same aged mouse. j) Quantification of λ:κ light chain ratios in splenic and thymic B cells to assess B cell clonality. The data considers up to 3 B cell aggregates per tissue group and is represented as mean +/- SD. k) Gene expression profiles of selected cell types within the BC aggr and Fibr I regions (Fig. 4a ii) of the aged spatial transcriptomics dataset. Absolute cell numbers are indicated in parentheses.

## References

1. López-Otín, C., Blasco, M.A., Partridge, L., et al., Hallmarks of aging: An expanding universe, Cell (186), 2023, 243–278. 10.1016/j.cell.2022.11.001.

2. Fafián-Labora, J.A., and O’Loghlen, A., Classical and Nonclassical Intercellular Communication in Senescence and Ageing, Trends in Cell Biology (30), 2020, 628–639. 10.1016/j.tcb.2020.05.003.

3. Goyani, P., Christodoulou, R., and Vassiliou, E., Immunosenescence: Aging and Immune System Decline, Vaccines (12), 2024, 1314. 10.3390/vaccines12121314.

4. Lee, K.-A., Flores, R.R., Jang, I.H., et al., Immune Senescence, Immunosenescence and Aging, Frontiers in Aging (3), 2022,. 10.3389/fragi.2022.900028.

5. Liu, Q., Zheng, Y., Goronzy, J.J., et al., T cell aging as a risk factor for autoimmunity, Journal of Autoimmunity (137), 2023, 102947. 10.1016/j.jaut.2022.102947.

6. Shaw, A.C., Joshi, S., Greenwood, H., et al., Aging of the Innate Immune System, Current opinion in immunology (22), 2010, 507–513. 10.1016/j.coi.2010.05.003.

7. Liu, Z., Liang, Q., Ren, Y., et al., Immunosenescence: molecular mechanisms and diseases, Signal Transduction and Targeted Therapy (8), 2023, 1–16. 10.1038/s41392-023-01451-2.

8. Thomas, R., Wang, W., and Su, D.-M., Contributions of Age-Related Thymic Involution to Immunosenescence and Inflammaging, Immunity & Ageing (17), 2020, 2. 10.1186/s12979-020-0173-8.

9. Ferrucci, L., and Fabbri, E., Inflammageing: chronic inflammation in ageing, cardiovascular disease, and frailty, Nature Reviews Cardiology (15), 2018, 505–522. 10.1038/s41569-018-0064-2.

10. Ovadya, Y., Landsberger, T., Leins, H., et al., Impaired immune surveillance accelerates accumulation of senescent cells and aging, Nature Communications (9), 2018, 5435. 10.1038/s41467-018-07825-3.

11. Nikolich-Žugich, J., Ageing and life-long maintenance of T-cell subsets in the face of latent persistent infections, Nature Reviews Immunology (8), 2008, 512–522. 10.1038/nri2318.

12. Elyahu, Y., Hekselman, I., Eizenberg-Magar, I., et al., Aging promotes reorganization of the CD4 T cell landscape toward extreme regulatory and effector phenotypes, Science Advances (5), 2019, eaaw8330. 10.1126/sciadv.aaw8330.

13. Goronzy, J.J., and Weyand, C.M., Immune aging and autoimmunity, Cellular and Molecular Life Sciences (69), 2012, 1615–1623. 10.1007/s00018-012-0970-0.

14. Thapa, P., and Farber, D.L., The Role of the Thymus in the Immune Response, Thoracic Surgery Clinics (29), 2019, 123–131. 10.1016/J.THORSURG.2018.12.001.

15. Cowan, J.E., Takahama, Y., Bhandoola, A., et al., Postnatal Involution and Counter- Involution of the Thymus, Frontiers in Immunology (11), 2020, 1–11. 10.3389/fimmu.2020.00897.

16. Hale, L.P., Histologic and molecular assessment of human thymus, Annals of Diagnostic Pathology (8), 2004, 50–60. 10.1016/j.anndiagpath.2003.11.006.

17. Baran-Gale, J., Morgan, M.D., Maio, S., et al., Ageing compromises mouse thymus function and remodels epithelial cell differentiation, eLife (9), 2020, 1–27. 10.7554/ELIFE.56221.

18. Liang, Z., Dong, X., Zhang, Z., et al., Age-related thymic involution: Mechanisms and functional impact, Aging Cell (21), 2022, e13671. 10.1111/acel.13671.

19. Klein, L., and Petrozziello, E., Antigen presentation for central tolerance induction, Nature Reviews Immunology (25), 2025, 57–72. 10.1038/s41577-024-01076-8.

20. Ashby, K.M., and Hogquist, K.A., A guide to thymic selection of T cells, Nature Reviews Immunology (24), 2024, 103–117. 10.1038/s41577-023-00911-8.

21. Daley, S.R., Teh, C., Hu, D.Y., et al., Cell death and thymic tolerance, Immunological Reviews (277), 2017, 9–20. 10.1111/imr.12532.

22. Palmer, D.B., The Effect of Age on Thymic Function, Frontiers in Immunology (4), 2013,. 10.3389/fimmu.2013.00316.

23. Kousa, A.I., Jahn, L., Zhao, K., et al., Age-related epithelial defects limit thymic function and regeneration, Nature Immunology (25), 2024, 1593–1606. 10.1038/s41590-024-01915-9.

24. Ye, P., and Kirschner, D.E., Reevaluation of T Cell Receptor Excision Circles as a Measure of Human Recent Thymic Emigrants1, The Journal of Immunology (168), 2002, 4968–4979. 10.4049/jimmunol.168.10.4968.

25. Coder, B., and Su, D.-M., Thymic involution beyond T-cell insufficiency, Oncotarget (6), 2015, 21777–21778. 10.18632/oncotarget.4970.

26. Srinivasan, J., Lancaster, J.N., Singarapu, N., et al., Age-Related Changes in Thymic Central Tolerance, Frontiers in Immunology (12), 2021,. 10.3389/fimmu.2021.676236.

27. Hidalgo, Y., Núñez, S., Fuenzalida, M.J., et al., Thymic B Cells Promote Germinal Center-Like Structures and the Expansion of Follicular Helper T Cells in Lupus-Prone Mice, Frontiers in Immunology (11), 2020,. 10.3389/fimmu.2020.00696.

28. Wedemeyer, S.A., and Griffith, A.V., Thymic B cells in aging and autoimmune disease, Frontiers in Immunology (16), 2025, 1595805. 10.3389/fimmu.2025.1595805.

29. Gorgoulis, V., Adams, P.D., Alimonti, A., et al., Cellular Senescence: Defining a Path Forward, Cell (179), 2019, 813–827. 10.1016/j.cell.2019.10.005.

30. Li, X., Li, C., Zhang, W., et al., Inflammation and aging: signaling pathways and intervention therapies, Signal Transduction and Targeted Therapy (8), 2023, 239. 10.1038/s41392-023-01502-8.

31. Li, M., Luan, F., Zhao, Y., et al., Epithelial-mesenchymal transition: An emerging target in tissue fibrosis, Experimental Biology and Medicine (241), 2016, 1–13. 10.1177/1535370215597194.

32. Yang, J., Liu, J., Liang, J., et al., Epithelial-mesenchymal transition in age-associated thymic involution: Mechanisms and therapeutic implications, Ageing Research Reviews (92), 2023, 102115. 10.1016/j.arr.2023.102115.

33. Xu, P.-X., Zheng, W., Laclef, C., et al., *Eya1* is required for the morphogenesis of mammalian thymus, parathyroid and thyroid, Development (129), 2002, 3033–3044. 10.1242/dev.129.13.3033.

34. Ghosh, A., and Fowler, V.M., Tropomodulins, Current Biology (31), 2021, R501–R503. 10.1016/j.cub.2021.01.055.

35. Li, Y., Pan, X., Gong, Z., et al., Tropomodulin-1—From the actin slow-growing end to multifunctional roles, FEBS Letters (600), 2026, 10–19. 10.1002/1873-3468.70176.

36. Zhang, J., Di, S., Li, M., et al., FAM107A as a tumor suppressor in esophageal squamous carcinoma inhibits growth and metastasis, Pathology - Research and Practice (252), 2023, 154945. 10.1016/j.prp.2023.154945.

37. Ma, W., Lin, J., Ma, R., et al., Insights into Structure and Function of Growth Arrest Specific 2 (GAS2), Journal of Cancer (16), 2025, 146–156. 10.7150/jca.102893.

38. Lehembre, F., Yilmaz, M., Wicki, A., et al., NCAM-induced focal adhesion assembly: a functional switch upon loss of E-cadherin, The EMBO Journal (27), 2008, 2603–2615. 10.1038/emboj.2008.178.

39. Shelly, M., Cancedda, L., Lim, B.K., et al., Semaphorin3A Regulates Neuronal Polarization by Suppressing Axon Formation and Promoting Dendrite Growth, Neuron, 2011, 433–446. 10.1016/j.neuron.2011.06.041.

40. Egea-Zorrilla, A., Blasco-Iturri, Z., Saez, B., et al., The Notch3 Pathway in Organ Fibrosis, Fibrosis (2), 2024, 10007–10007. 10.70322/fibrosis.2024.10007.

41. Zhang, Y., Xie, K., and Jiang, T., Roles and regulation of δ-catenin in tumorigenesis and neuronal diseases, Frontiers in Cell and Developmental Biology (13), 2025, 1559059. 10.3389/fcell.2025.1559059.

42. Buechler, M.B., Pradhan, R.N., Krishnamurty, A.T., et al., Cross-tissue organization of the fibroblast lineage, Nature (593), 2021, 575–579. 10.1038/s41586-021-03549-5.

43. Granadier, D., Acenas, D., and Dudakov, J.A., Endogenous thymic regeneration: restoring T cell production following injury, Nature Reviews Immunology (25), 2025, 407–424. 10.1038/s41577-024-01119-0.

44. Hauri-Hohl, M.M., Zuklys, S., Keller, M.P., et al., TGF-â signaling in thymic epithelial cells regulates thymic involution and postirradiation reconstitution, Blood (112), 2008, 626–634. 10.1182/blood-2007-10-115618.

45. Kumar, R., Langer, J.C., and Snoeck, H.-W., Transforming growth factor-β2 is involved in quantitative genetic variation in thymic involution, Blood (107), 2006, 1974–1979. 10.1182/blood-2005-04-1495.

46. Garfin, P.M., Min, D., Bryson, J.L., et al., Inactivation of the RB family prevents thymus involution and promotes thymic function by direct control of Foxn1 expression, Journal of Experimental Medicine (210), 2013, 1087–1097. 10.1084/jem.20121716.

47. Alessio, N., Squillaro, T., Di Bernardo, G., et al., Increase of circulating IGFBP-4 following genotoxic stress and its implication for senescence, eLife (9), 2020, e54523. 10.7554/eLife.54523.

48. Siraj, Y., Aprile, D., Alessio, N., et al., IGFBP7 is a key component of the senescence-associated secretory phenotype (SASP) that induces senescence in healthy cells by modulating the insulin, IGF, and activin A pathways, Cell Communication and Signaling (22), 2024, 540. 10.1186/s12964-024-01921-2.

49. Nusser, A., Sagar, Swann, J.B., et al., Developmental dynamics of two bipotent thymic epithelial progenitor types, Nature 2022 606:7912 (606), 2022, 165–171. 10.1038/s41586-022-04752-8.

50. Xiao, Y., Jin, W., Ju, L., et al., Tracking single-cell evolution using clock-like chromatin accessibility loci, Nature Biotechnology (43), 2025, 784–798. 10.1038/s41587-024-02241-z.

51. Cao, J., Spielmann, M., Qiu, X., et al., The single-cell transcriptional landscape of mammalian organogenesis, Nature (566), 2019, 496–502. 10.1038/s41586-019-0969-x.

52. Steier, Z., Aylard, D.A., McIntyre, L.L., et al., Single-cell multiomic analysis of thymocyte development reveals drivers of CD4+ T cell and CD8+ T cell lineage commitment, Nature Immunology (24), 2023, 1579–1590. 10.1038/s41590-023-01584-0.

53. Givony, T., Leshkowitz, D., Del Castillo, D., et al., Thymic mimetic cells function beyond self-tolerance, Nature (622), 2023, 164–172. 10.1038/s41586-023-06512-8.

54. Nusser, A., Thomas, O.S., Zhang, G., et al., Developmental trajectory and evolutionary origin of thymic mimetic cells, Nature, 2025, 1–10. 10.1038/s41586-025-09148-y.

55. Sansom, S.N., Shikama-Dorn, N., Zhanybekova, S., et al., Population and single-cell genomics reveal the Aire dependency, relief from Polycomb silencing, and distribution of self-antigen expression in thymic epithelia, Genome Research (24), 2014, 1918–1931. 10.1101/gr.171645.113.

56. Bull, J.A., Moore, J.W., Mulholland, E.J., et al., MuSpAn: A Toolbox for Multiscale Spatial Analysis,, 2025, 2024.12.06.627195. 10.1038/s41467-026-75649-7.

57. Klein, F., Veiga-Villauriz, C., Börsch, A., et al., Combined multidimensional single- cell protein and RNA profiling dissects the cellular and functional heterogeneity of thymic epithelial cells, Nature Communications (14), 2023, 1–16. 10.1038/s41467-023-39722-9.

58. Laidlaw, B.J., Duan, L., Xu, Y., et al., The transcription factor Hhex cooperates with the corepressor Tle3 to promote memory B cell development, Nature Immunology (21), 2020, 1082–1093. 10.1038/s41590-020-0713-6.

59. Chaudhuri, J., Evans, T., Kumar, R., et al., Biological function of activation-induced cytidine deaminase (AID), Biomedical Journal (37), 2014, 269. 10.4103/2319-4170.128734.

60. Ochiai, K., Maienschein-Cline, M., Simonetti, G., et al., Transcriptional Regulation of Germinal Center B and Plasma Cell Fates by Dynamical Control of IRF4, Immunity (38), 2013, 918–929. 10.1016/j.immuni.2013.04.009.

61. Harrer, C., Otto, F., Radlberger, R.F., et al., The CXCL13/CXCR5 Immune Axis in Health and Disease—Implications for Intrathecal B Cell Activities in Neuroinflammation, Cells (11), 2022, 2649. 10.3390/cells11172649.

62. Zhao, L., Jin, S., Wang, S., et al., Tertiary lymphoid structures in diseases: immune mechanisms and therapeutic advances, Signal Transduction and Targeted Therapy (9), 2024, 225. 10.1038/s41392-024-01947-5.

63. Mackay, F., and Schneider, P., Cracking the BAFF code, Nature Reviews Immunology (9), 2009, 491–502. 10.1038/nri2572.

64. Allen, C.D.C., Okada, T., and Cyster, J.G., Germinal-Center Organization and Cellular Dynamics, Immunity (27), 2007, 190–202. 10.1016/j.immuni.2007.07.009.

65. Carroll, M.C., The complement system in regulation of adaptive immunity, Nature Immunology (5), 2004, 981–986. 10.1038/ni1113.

66. Bouloudani, T., and Sautès-Fridman, C., Fibroblasts hold the key to TLS formation, Immunity (59), 2026, 1490–1492. 10.1016/j.immuni.2026.05.009.

67. Jin, S., Plikus, M.V., and Nie, Q., CellChat for systematic analysis of cell–cell communication from single-cell transcriptomics, Nature Protocols (20), 2025, 180–219. 10.1038/s41596-024-01045-4.

68. Gordon, J., and Manley, N.R., Mechanisms of thymus organogenesis and morphogenesis, Development (138), 2011, 3865–3878. 10.1242/dev.059998.

69. Gordon, J., Patel, S.R., Mishina, Y., et al., Evidence for an early role for BMP4 signaling in thymus and parathyroid morphogenesis, Developmental Biology (339), 2010, 141–154. 10.1016/j.ydbio.2009.12.026.

70. Rossi, S.W., Jeker, L.T., Ueno, T., et al., Keratinocyte growth factor (KGF) enhances postnatal T-cell development via enhancements in proliferation and function of thymic epithelial cells, Blood (109), 2007, 3803–3811. 10.1182/BLOOD-2006-10-049767.

71. Czarkwiani, A., Lobo, M., Castro, L.A.B., et al., Molecular basis for de novo thymus regeneration in a vertebrate, the axolotl, SCienCe immunology, 2025,.

72. Deuel, T.F., Zhang, N., Yeh, H.-J., et al., Pleiotrophin: A Cytokine with Diverse Functions and a Novel Signaling Pathway, Archives of Biochemistry and Biophysics (397), 2002, 162–171. 10.1006/abbi.2001.2705.

73. Chen, F., Lyu, L., Xing, C., et al., The pivotal role of TGF-β/Smad pathway in fibrosis pathogenesis and treatment, Frontiers in Oncology (15), 2025, 1649179. 10.3389/fonc.2025.1649179.

74. Tam, K.J., Hui, D.H.F., Lee, W.W., et al., Semaphorin 3 C drives epithelial-to- mesenchymal transition, invasiveness, and stem-like characteristics in prostate cells, Scientific Reports (7), 2017, 11501. 10.1038/s41598-017-11914-6.

75. Deng, Y., Peng, Z., Ming, K., et al., Single-cell analysis of human thymus and peripheral blood unveils the dynamics of T cell development and aging, Nature Aging, 2025,. 10.1038/s43587-025-00990-3.

76. Sato, K., Kato, A., Sekai, M., et al., Physiologic Thymic Involution Underlies Age- Dependent Accumulation of Senescence-Associated CD4+ T Cells, The Journal of Immunology (199), 2017, 138–148. 10.4049/jimmunol.1602005.

77. Zhu, X., Gui, J., Dohkan, J., et al., Lymphohematopoietic progenitors do not have a synchronized defect with age-related thymic involution, Aging Cell (6), 2007, 663–672. 10.1111/j.1474-9726.2007.00325.x.

78. Sun, L., Brown, R., Chen, S., et al., Aging induced decline in T-lymphopoiesis is primarily dependent on status of progenitor niches in the bone marrow and thymus, Aging (4), 2012, 606–619. 10.18632/aging.100487.

79. Pinto, A.I., Smith, J., Kissack, M.R., et al., Thymic B Cell-Mediated Attack of Thymic Stroma Precedes Type 1 Diabetes Development, Frontiers in Immunology (9), 2018,. 10.3389/fimmu.2018.01281.

80. Castañeda, J., Hidalgo, Y., Sauma, D., et al., The Multifaceted Roles of B Cells in the Thymus: From Immune Tolerance to Autoimmunity, Frontiers in Immunology (12), 2021,. 10.3389/fimmu.2021.766698.

81. Nuñez, S., Moore, C., Gao, B., et al., The human thymus perivascular space is a functional niche for viral-specific plasma cells, Science Immunology (1), 2016, eaah4447. 10.1126/sciimmunol.aah4447.

82. Stuart, T., Butler, A., Hoffman, P., et al., Comprehensive Integration of Single-Cell Data, Cell (177), 2019, 1888–1902. 10.1016/j.cell.2019.05.031.

83. Stuart, T., Srivastava, A., Madad, S., et al., Single-cell chromatin state analysis with Signac, Nature Methods (18), 2021, 1333–1341. 10.1038/s41592-021-01282-5.

84. McGinnis, C.S., Murrow, L.M., and Gartner, Z.J., DoubletFinder: Doublet Detection in Single-Cell RNA Sequencing Data Using Artificial Nearest Neighbors, Cell Systems (8), 2019, 329–337.e4. 10.1016/j.cels.2019.03.003.

85. Zhang, Y., Parmigiani, G., and Johnson, W.E., ComBat-seq: Batch effect adjustment for RNA-seq count data, NAR Genomics and Bioinformatics (2), 2020,. 10.1093/nargab/lqaa078.

86. Mulholland-Illingworth, E.J., Moore, J.W., Lin, M., et al., Therapeutic manipulation and spatial quantification of the tumor microenvironment in colorectal cancer, iScience (29), 2026, 115193. 10.1016/j.isci.2026.115193.

87. Getis, A., and Ord, J.K., The Analysis of Spatial Association by Use of Distance Statistics, Geographical Analysis (24), 1992, 189–206. 10.1111/j.1538-4632.1992.tb00261.x.

88. Morueta-Holme, N., Blonder, B., Sandel, B., et al., A network approach for inferring species associations from co-occurrence data, Ecography (39), 2016, 1139–1150. 10.1111/ecog.01892.

89. Moore, J.W., Bull, J.A., and Byrne, H.M., netPCF: Geometry-aware pair correlation functions for spatial biology, .

90. Bull, J.A., Mulholland, E.J., Leedham, S.J., et al., Extended correlation functions for spatial analysis of multiplex imaging data, Biological Imaging (4), 2024, e2. 10.1017/S2633903X24000011.

91. Virtanen, P., Gommers, R., Oliphant, T.E., et al., SciPy 1.0: fundamental algorithms for scientific computing in Python, Nature Methods (17), 2020, 261–272. 10.1038/s41592-019-0686-2.

92. Goltsev, Y., Samusik, N., Kennedy-Darling, J., et al., Deep Profiling of Mouse Splenic Architecture with CODEX Multiplexed Imaging, Cell (174), 2018, 968–981. 10.1016/J.CELL.2018.07.010.

93. Stringer, C., Wang, T., Michaelos, M., et al., Cellpose: a generalist algorithm for cellular segmentation, Nature Methods (18), 2020, 100–106. 10.1038/s41592-020-01018-x.

94. Jin, S., Guerrero-Juarez, C.F., Zhang, L., et al., Inference and analysis of cell-cell communication using CellChat, Nature Communications (12), 2021, 1088. 10.1038/s41467-021-21246-9.

