## Supplementary Figures for "Spatial remodeling of the thymic stroma with age disrupts niches for T cell selection and tolerance"

**Supplementary Figure 1.1**

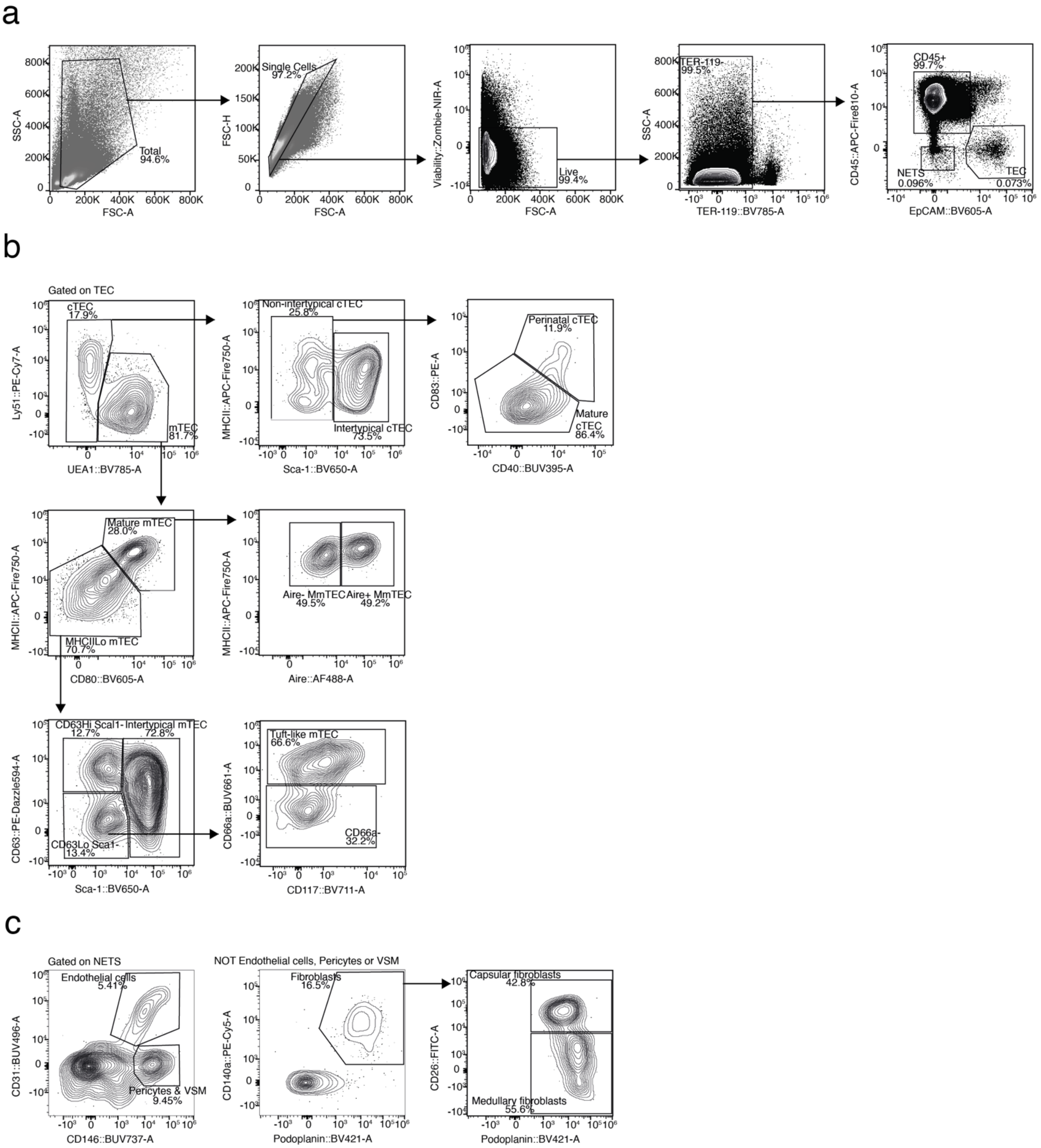


**Supplementary Fig. 1.1:** Flow cytometry gating strategies for a) intrathymic CD45+ cells, thymic epithelial cells (TEC) and non-epithelial thymic stromal cells (NETS) after enzymatic digestion. b) TEC subpopulations and c) NETS subpopulations. Shown are representative plots from a young thymus; (M) mTEC = (Mature) medullary thymic epithelial cells, VSM = vascular smooth muscle cells.

**Supplementary Figure 1.2**

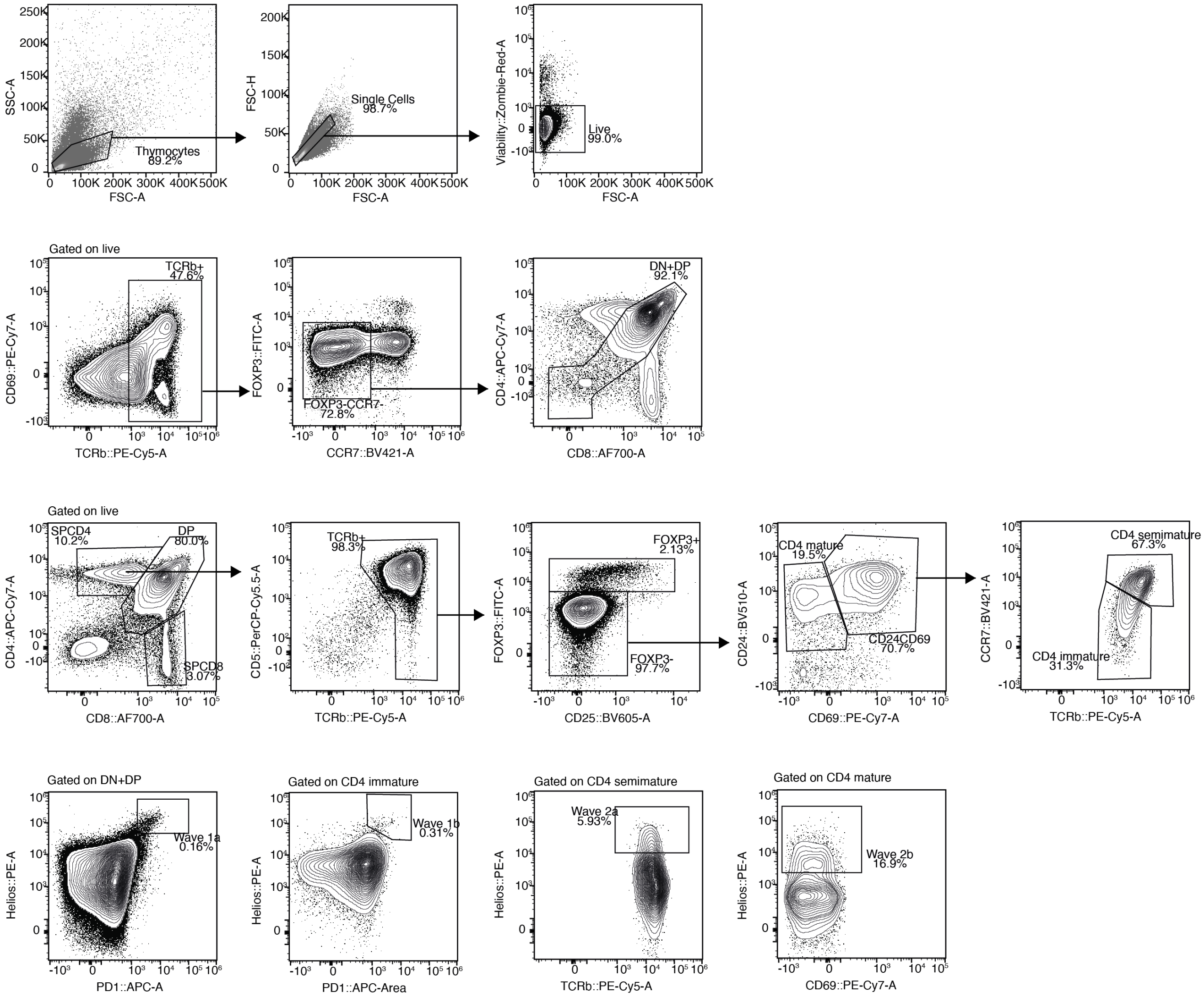


**Supplementary Fig. 1.2:** Flow cytometry gating strategies for thymocytes following mechanical dissociation of thymus lobes. For post-selectional maturation stages (single positive immature, semimature and mature) gating for the CD4 lineage only is shown as gating for CD8 lineage followed the same principles with the exception of not requiring exclusion of FOXP3 expressing cells. Plots in lower row show gating strategy for thymocytes undergoing cortical (wave 1a and b) and medullary (waves 2a and b) negative selection.[1]; DN = double negative, DP = double positive, SP = single positive.

**Supplementary Figure 1.3**

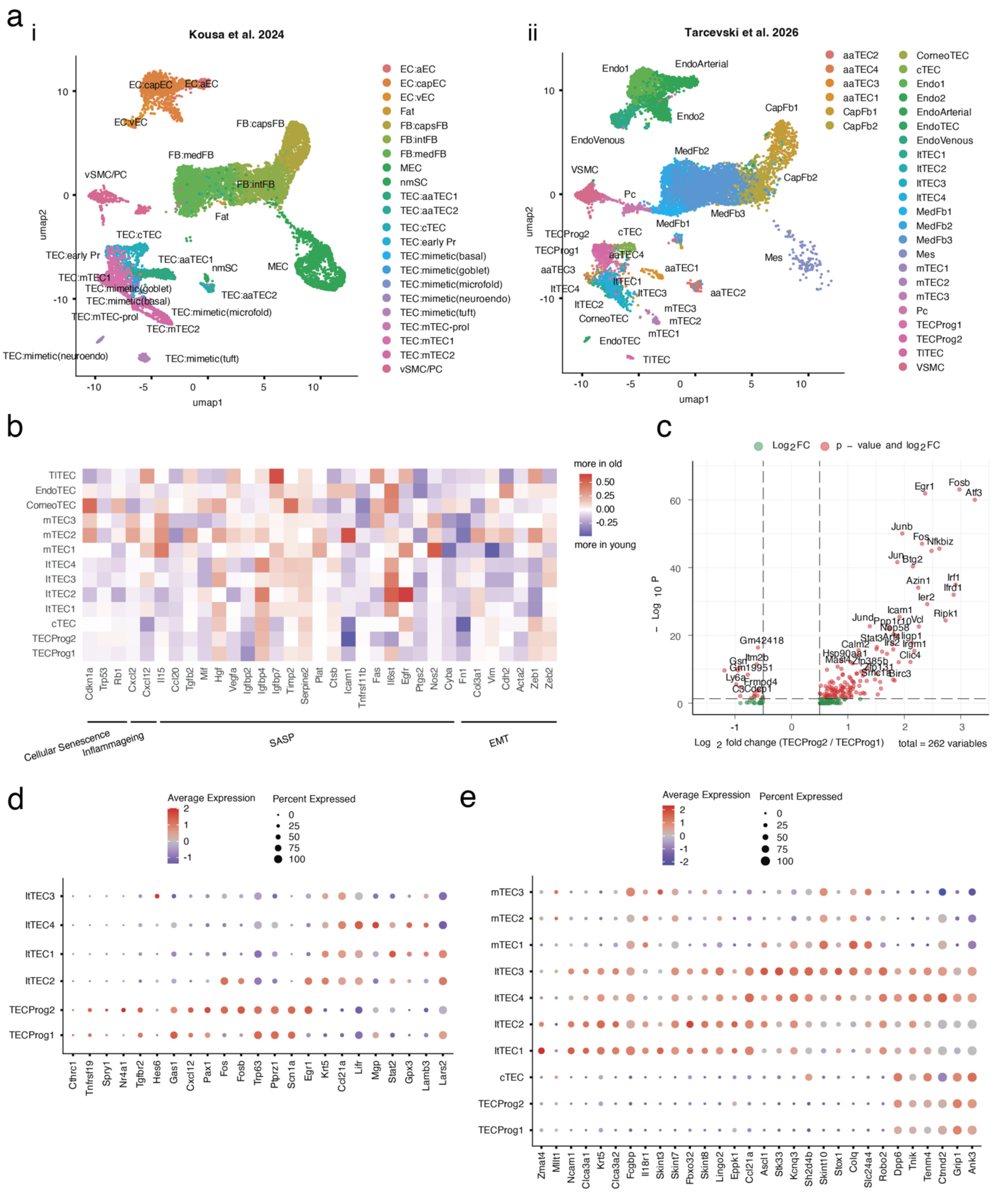


**Supplementary Fig. 1.3:** a) Integration of a published scRNAseq dataset[2] with our stromal snRNAseq dataset to align annotations of age-associated thymic epithelial cells (aaTECs). b) Expression of genes associated with aging-related processes, including cellular senescence, inflammaging, senescence-associated secretory pathway (SASP), and epithelial-to-mesenchymal transition (EMT).[3–6] c) Volcano plot showing differential gene expression between the two TEC progenitor populations. d)

Literature-adapted[7] gene lists to illustrate the similarity of our TEC progenitor and intertypical TEC (ItTEC) populations to previously-described embryonic and postnatal TEC progenitor populations. e) Expression of differentially expressed intertypical TEC markers highlighting the transcriptional similarities between TEC progenitors, ItTECs, cortical TECs (cTECs), and medullary TECs (mTECs).

**Supplementary Figure 2.1**

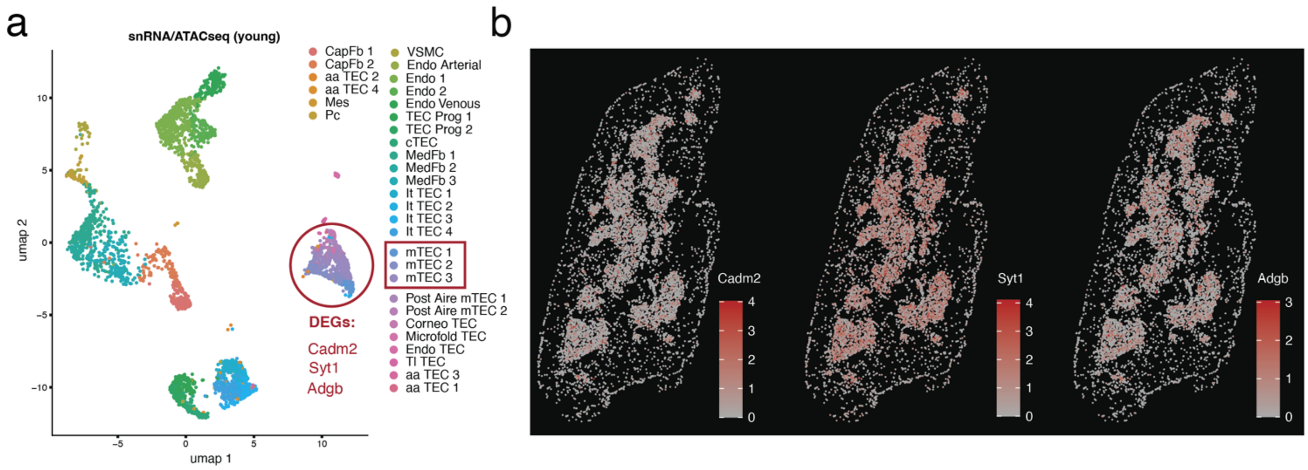


**Supplementary Fig. 2.1:** a) UMAP-projection of the annotated, snRNA/ATACseq stromal dataset for the young thymus. A DGE analysis identified target genes absent in the spatial transcriptomics panel across Aire-positive mTEC populations (1-3) including *Cadm2, Syt1,* and *Adgb*. b) Spatial gene expression of cross-modally integrated datasets after propagation of molecular information from the snRNA/ATACseq reference to the spatial transcriptomic dataset. The panel-absent, differentially expressed genes are highly expressed at their expected medullary location.

**Supplementary Figure 2.2**

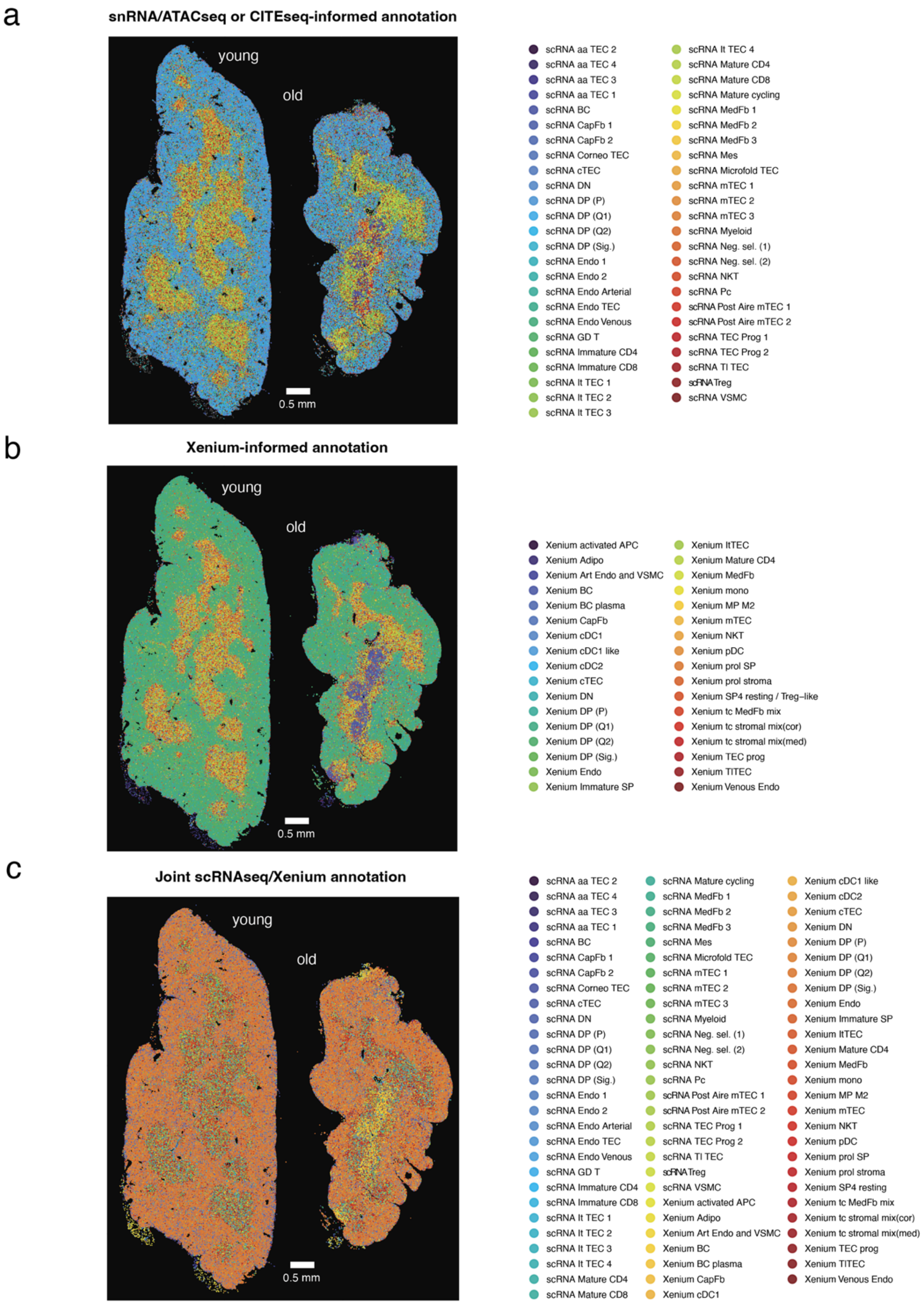


**Supplementary Fig. 2.2:** a-c) Cell type annotation and spatial localization of individual cells in the spatial transcriptomics dataset using three complementary annotation strategies. a) Cell type annotations obtained by label transfer from the snRNA/ATACseq and CITEseq reference datasets. b) Cell type annotations generated relying on the spatial transcriptomics data. c) Final integrated annotation combining both approaches. Where available, high-resolution annotations (scRNA) transferred from the snRNA/ATACseq and CITEseq references were retained in preference to the broader spatial transcriptomics-based annotation (Xenium).

**Supplementary Figure 2.3**

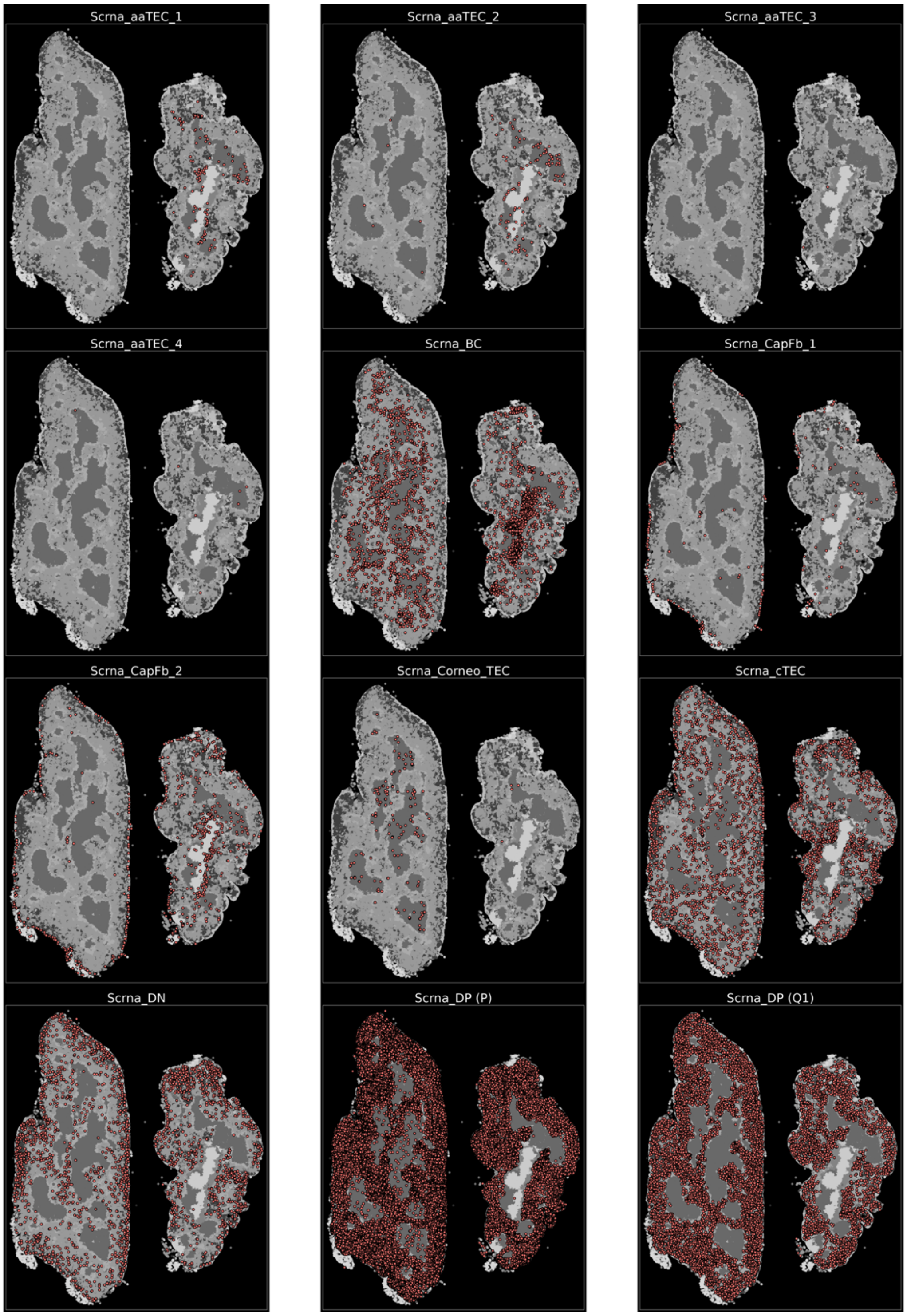


**Supplementary Fig. 2.3:** Overview of the spatial distribution of individual cell types following cross-modal integration in the young (left) and old (right) thymus profiled by spatial transcriptomics.

**Supplementary Figure 2.4**

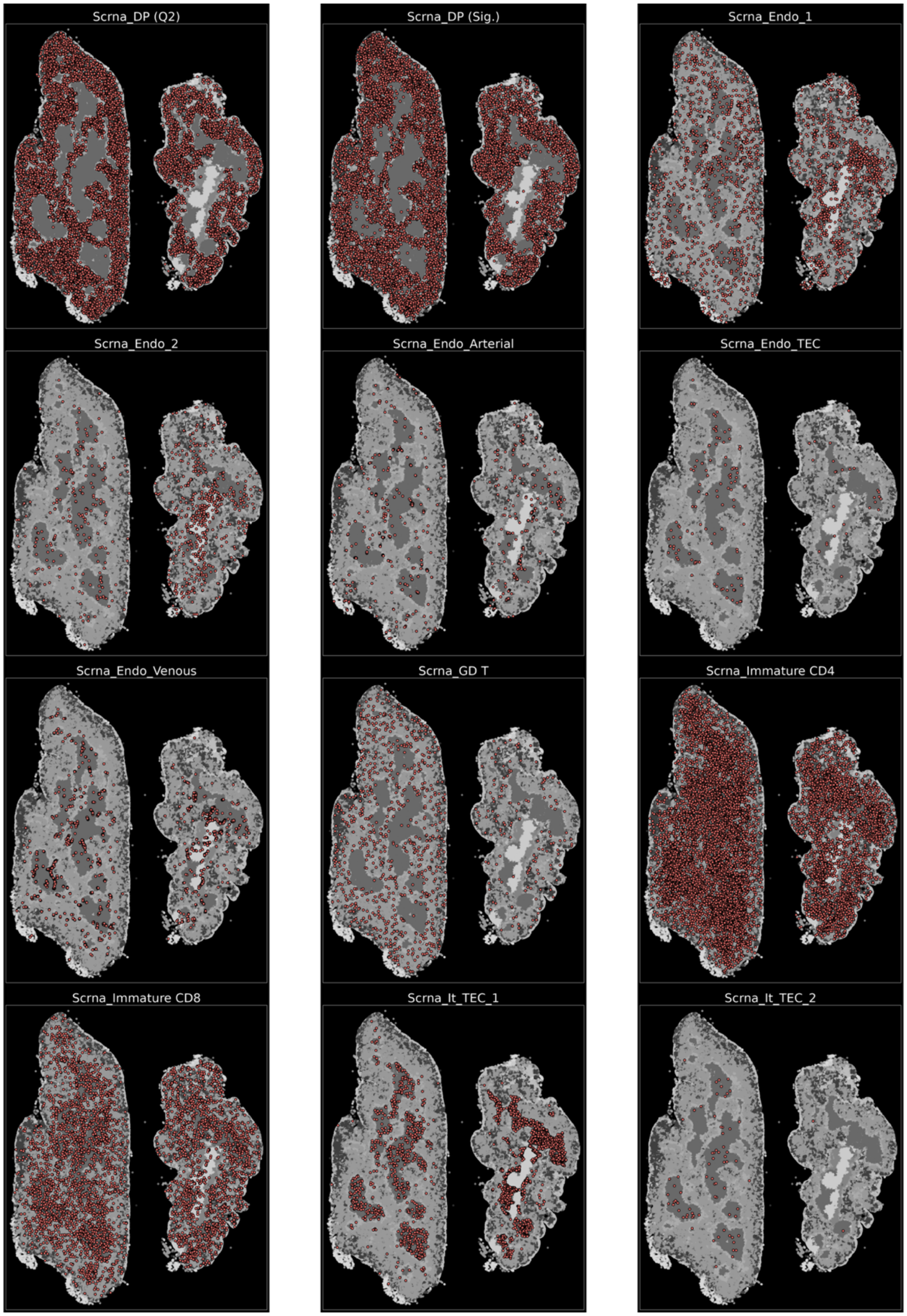


**Supplementary Fig. 2.4:** Overview of the spatial distribution of individual cell types following cross-modal integration in the young (left) and old (right) thymus profiled by spatial transcriptomics.

**Supplementary Figure 2.5**

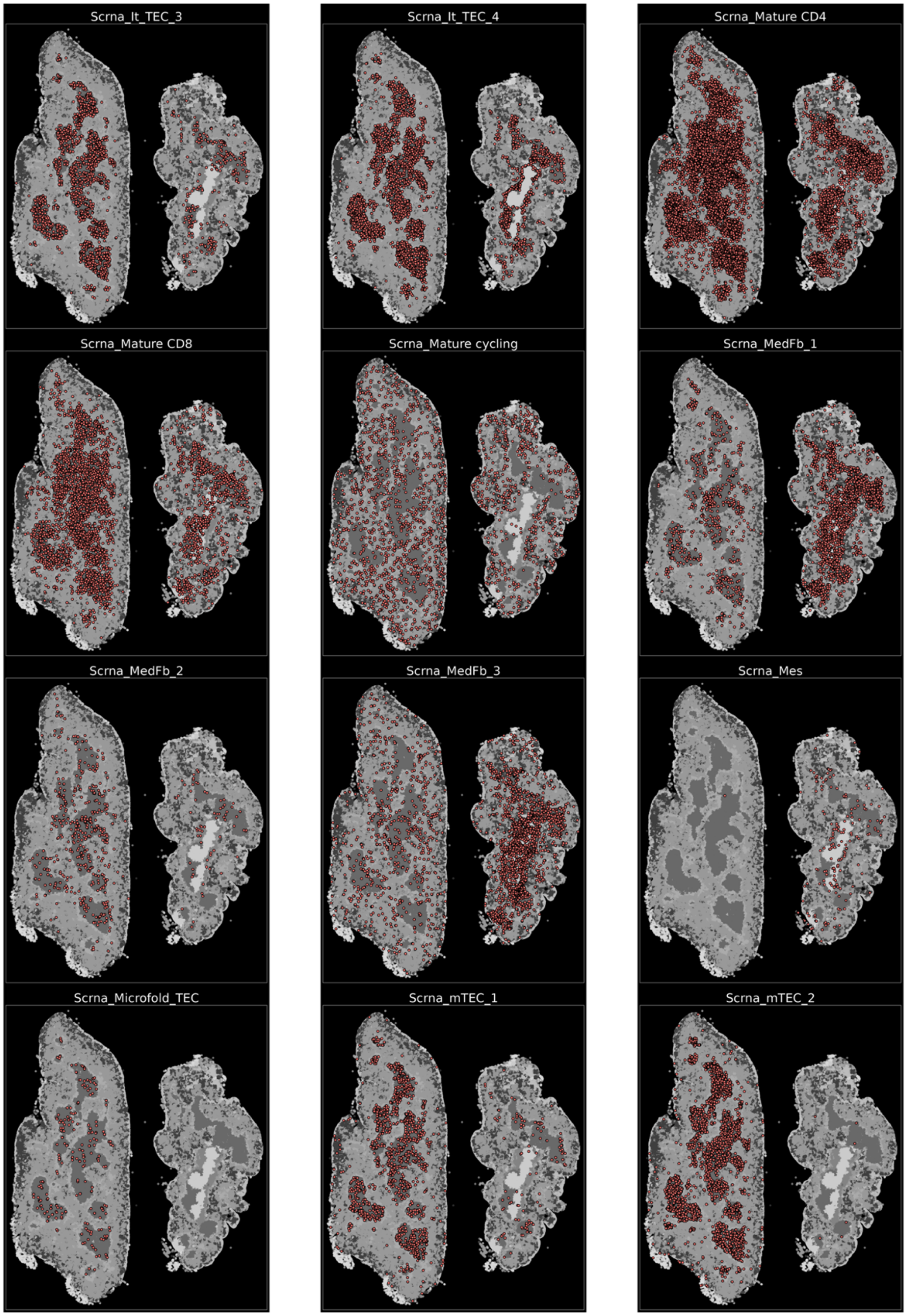


**Supplementary Fig. 2.5:** Overview of the spatial distribution of individual cell types following cross-modal integration in the young (left) and old (right) thymus profiled by spatial transcriptomics.

**Supplementary Figure 2.6**

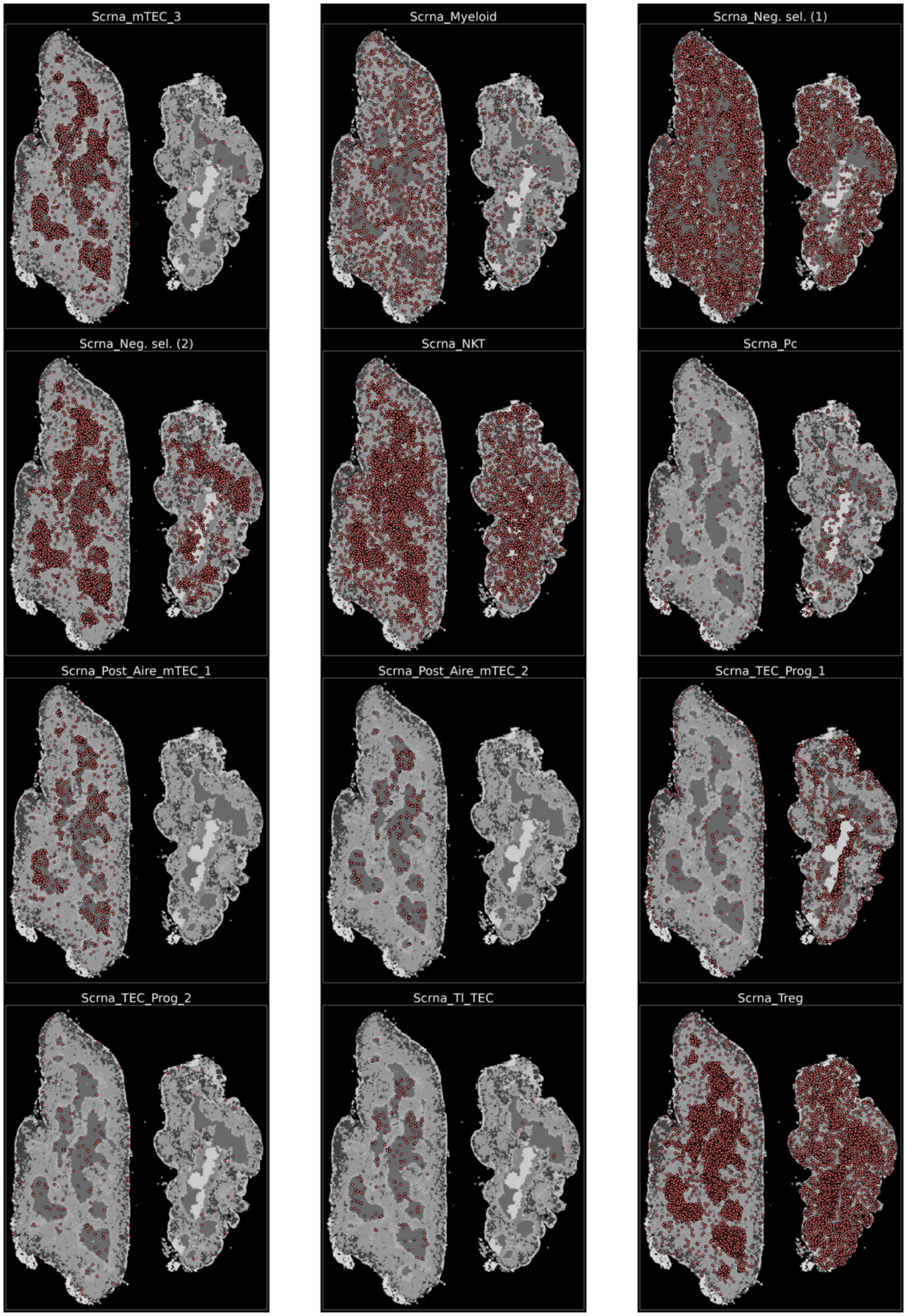


**Supplementary Fig. 2.6:** Overview of the spatial distribution of individual cell types following cross-modal integration in the young (left) and old (right) thymus profiled by spatial transcriptomics.

**Supplementary Figure 2.7**

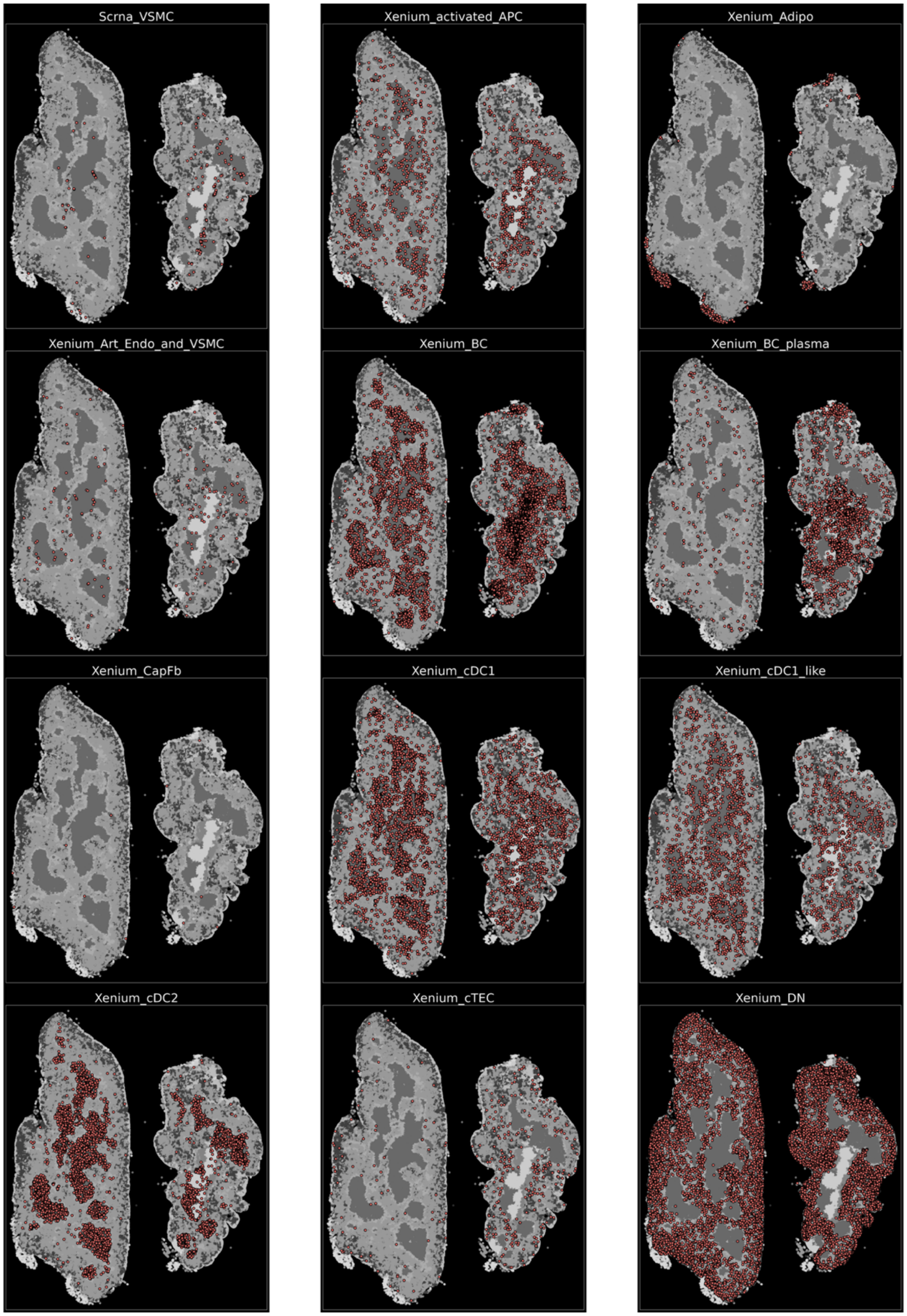


**Supplementary Fig. 2.7:** Overview of the spatial distribution of individual cell types following cross-modal integration in the young (left) and old (right) thymus profiled by spatial transcriptomics.

**Supplementary Figure 2.8**

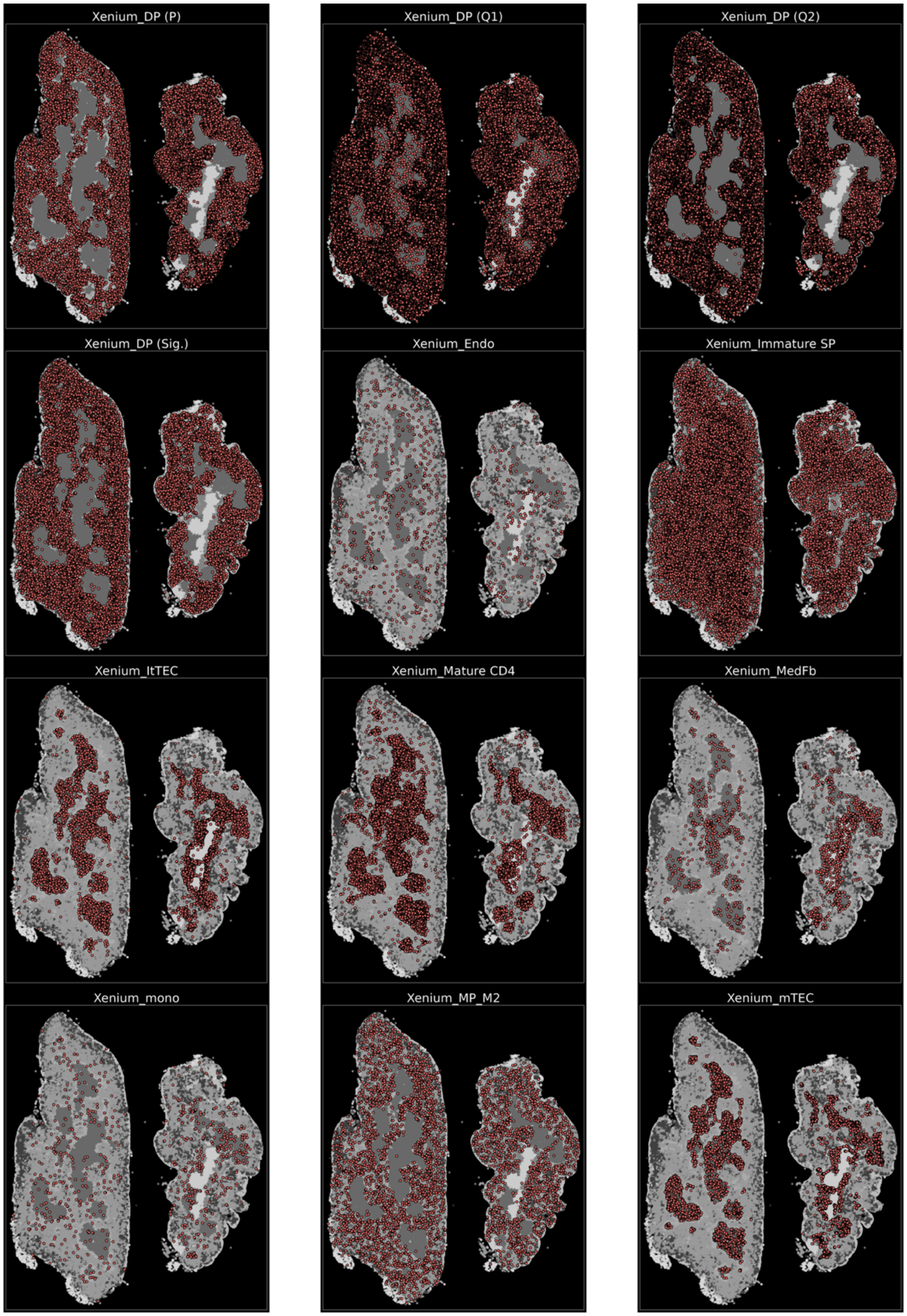


**Supplementary Fig. 2.8:** Overview of the spatial distribution of individual cell types following cross-modal integration in the young (left) and old (right) thymus profiled by spatial transcriptomics.

**Supplementary Figure 2.9**

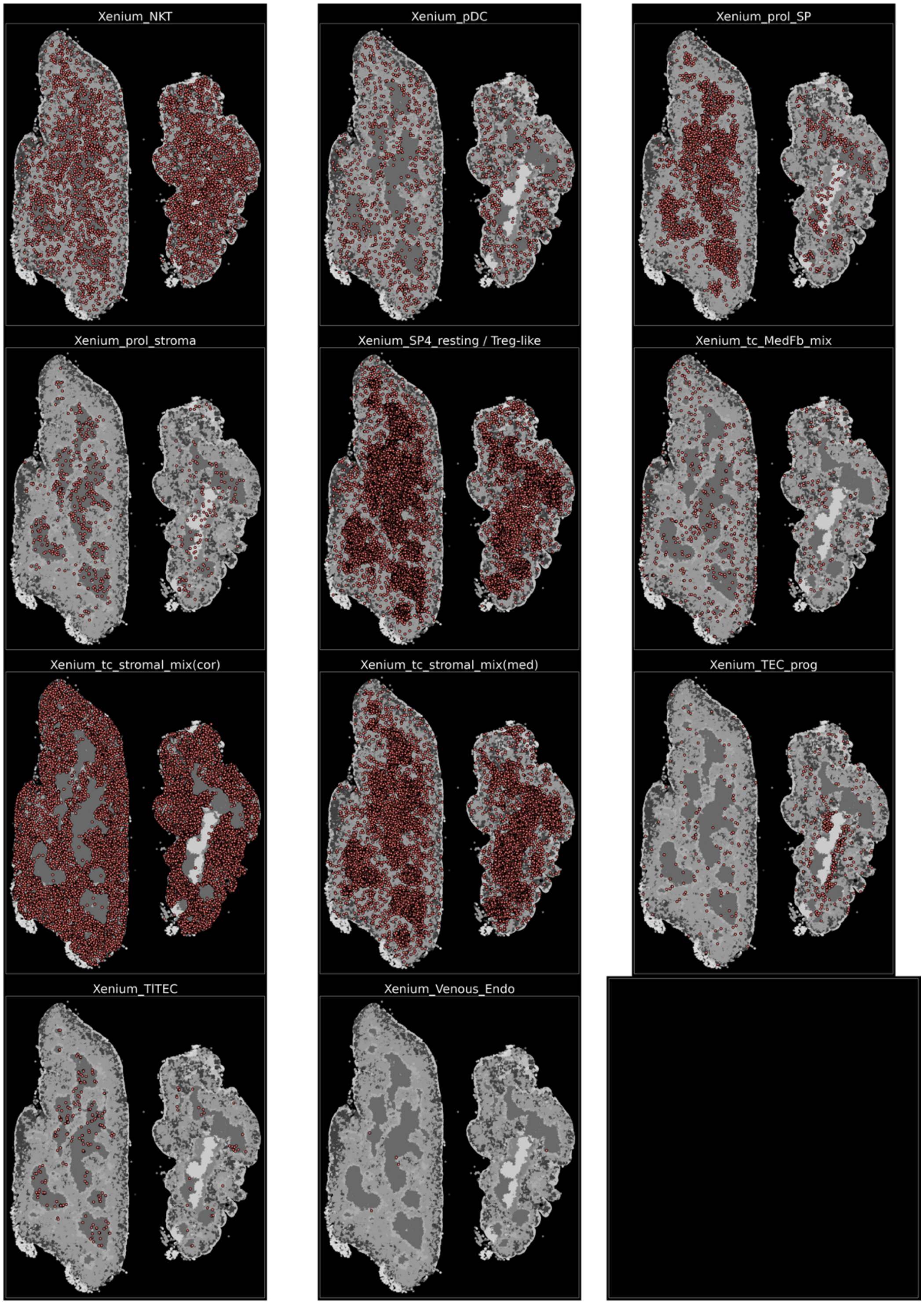


**Supplementary Fig. 2.9:** Overview of the spatial distribution of individual cell types following cross-modal integration in the young (left) and old (right) thymus profiled by spatial transcriptomics.

**Supplementary Figure 3**

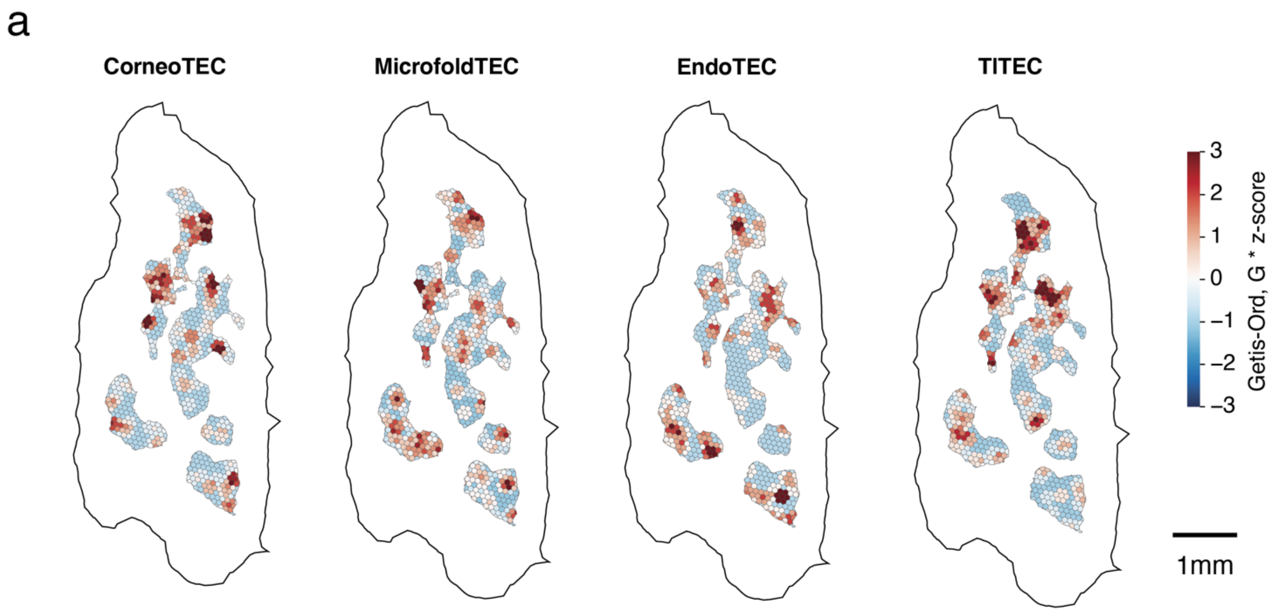


**Supplementary Fig. 3:** a) Spatial cell type enrichment of distinct mimetic TEC subtypes across hexagonal bins in the young spatial transcriptomics dataset.

**Supplementary Figure 4**

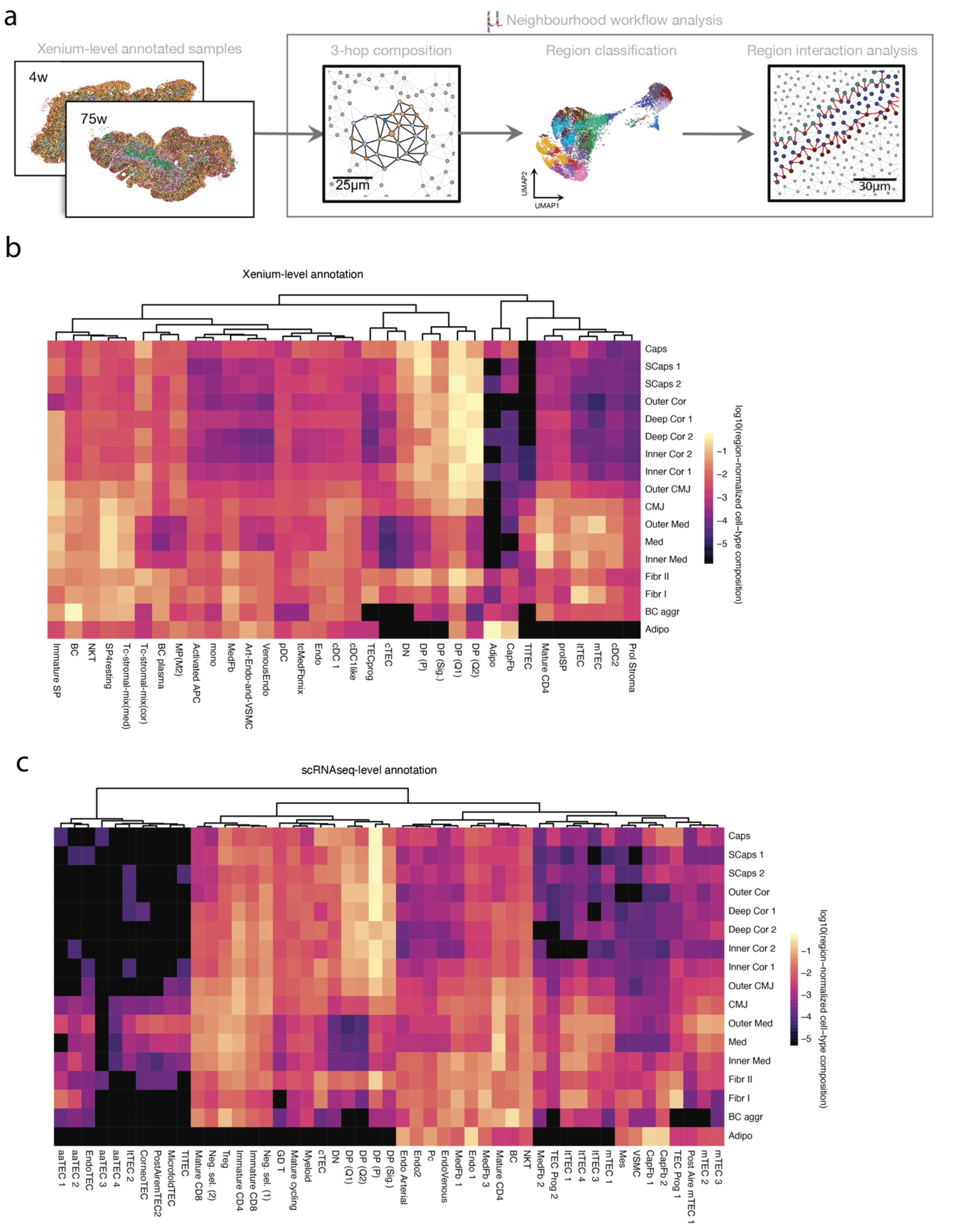


**Supplementary Fig. 4:** a) Schematic MuSpAn[8] spatial analysis workflow used to define global tissue neighborhoods in the young and old spatial transcriptomics datasets. Cell annotations from the spatial transcriptomics dataset were subjected to threehop neighborhood clustering, resulting in the identification of 17 functional regions that were subsequently analyzed for region-region interactions. b) Region-normalized cell type composition of the 17 functional regions based on the spatial transcriptomics (Xenium)-derived annotations. c) Region-normalized cell type composition of the same functional regions for the snRNA-seq/ATAC-seq and CITE-seq-level annotations obtained following label transfer.

**Supplementary Figure 7**

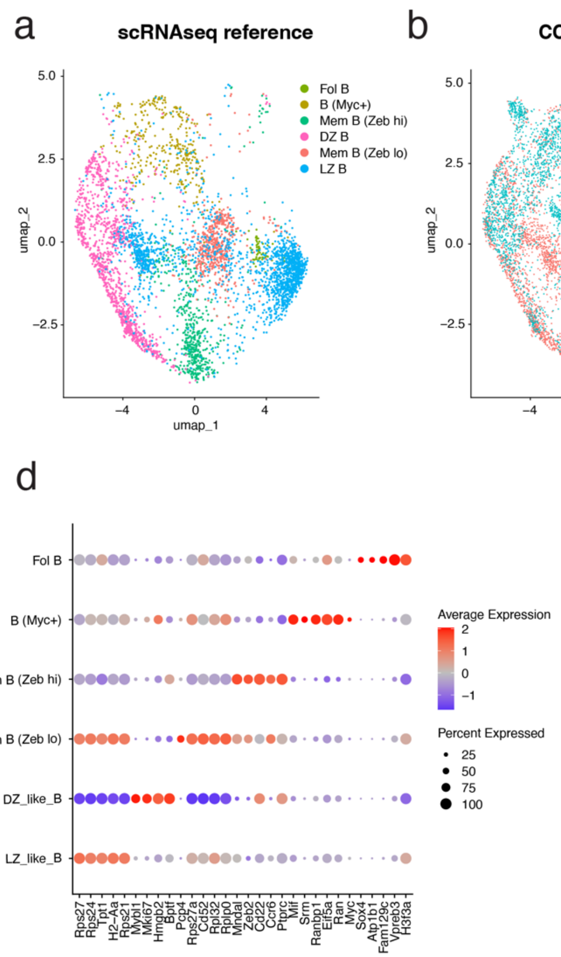


**Supplementary Fig. 7:** a) Re-annotated UMAP projection of the adapted splenic scRNA-seq reference dataset.[9] b) Integrated UMAP projection of the scRNA-seq reference and the central B-cell aggregate (Fig. 7a(i,ii)) from the aged spatial transcriptomics dataset. c) UMAP projection showing the cell-type annotations obtained by label transfer to B cells in the spatial transcriptomics dataset. d) Expression of representative marker genes defining the identified B-cell subtypes across the spatial transcriptomics data.

**Supplementary Figure 8**

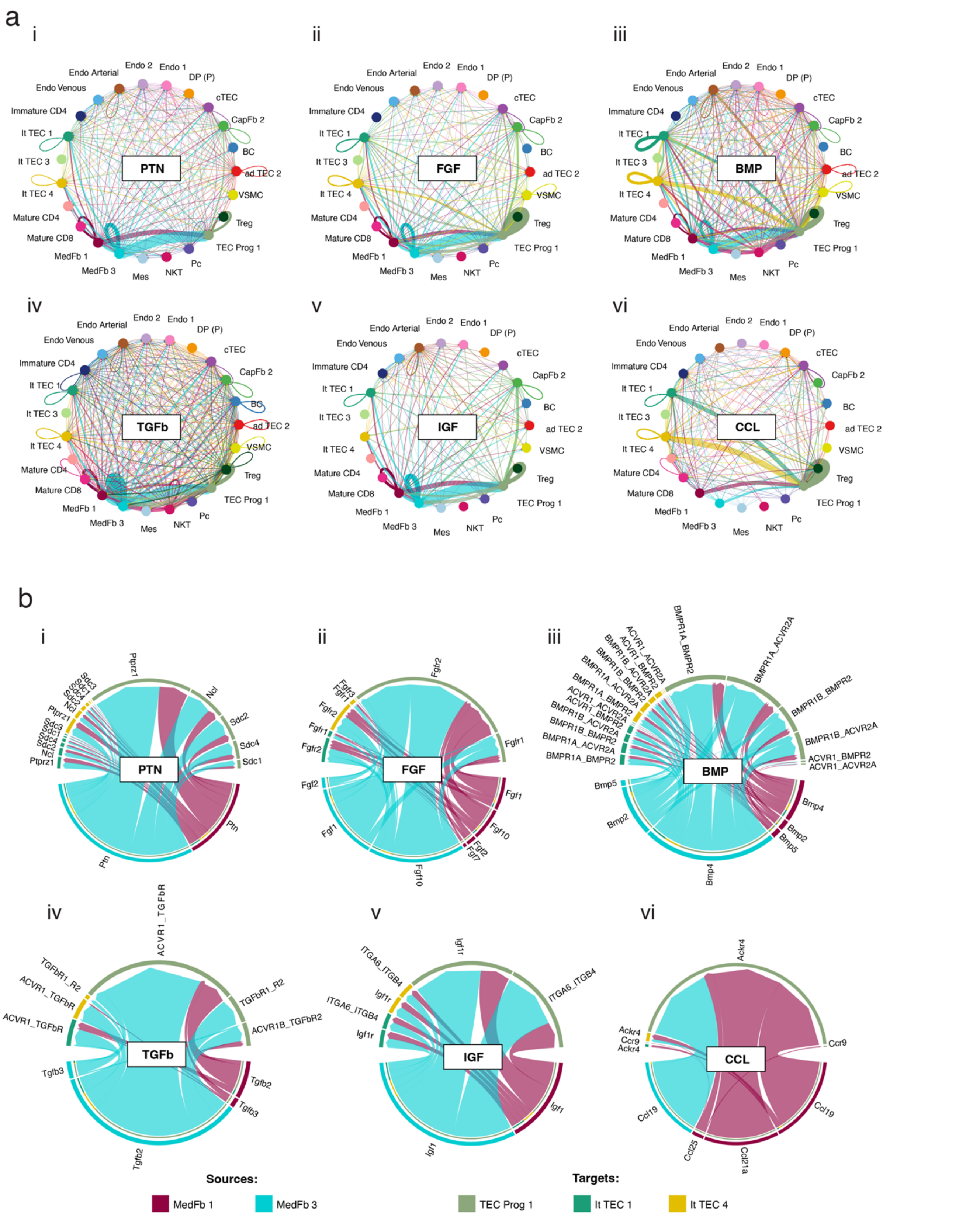


**Supplementary Fig. 8:** a) Weighted interaction networks showing the predicted strengths of paracrine signaling through the pleiotrophin (PTN), fibroblast growth factor (FGF), bone morphogenetic protein (BMP), transforming growth factor-β (TGFβ), insulin-like growth factor (IGF), and C-C motif chemokine ligand (CCL) pathways within the fibroblast-rich region of the old spatial transcriptomics dataset (Fibr I, 16; Fig. 4a(ii)). b) Chord plots illustrating individual ligand-receptor interactions within these signaling pathways between the selected source populations (medullary fibroblasts 1 and 3) and target populations (TEC progenitors and intertypical TECs) of the same region in the old spatial transcriptomics dataset.

### References


1. Daley, S.R., Hu, D.Y., and Goodnow, C.C., Helios marks strongly autoreactive CD4+ T cells in two major waves of thymic deletion distinguished by induction of PD-1 or NF-κB, *Journal of Experimental Medicine* (210) ,**2013**, 269–285. https://doi.org/10.1084/jem.20121458.

2. Steier, Z., Aylard, D.A., McIntyre, L.L., et al., Single-cell multiomic analysis of thymocyte development reveals drivers of CD4+ T cell and CD8+ T cell lineage commitment, *Nature Immunology* (24) ,**2023**, 1579–1590. https://doi.org/10.1038/s41590-023-01584-0.

3. Gorgoulis, V., Adams, P.D., Alimonti, A., et al., Cellular Senescence: Defining a Path Forward, *Cell* (179) ,**2019**, 813–827. https://doi.org/10.1016/j.cell.2019.10.005.

4. Li, X., Li, C., Zhang, W., et al., Inflammation and aging: signaling pathways and intervention therapies, *Signal Transduction and Targeted Therapy* (8) ,**2023**, 239. https://doi.org/10.1038/s41392-023-01502-8.

5. Li, M., Luan, F., Zhao, Y., et al., Epithelial-mesenchymal transition: An emerging target in tissue fibrosis, *Experimental Biology and Medicine* (241) ,**2016**, 1–13. https://doi.org/10.1177/1535370215597194.

6. Yang, J., Liu, J., Liang, J., et al., Epithelial-mesenchymal transition in age-associated thymic involution: Mechanisms and therapeutic implications, *Ageing Research Reviews* (92) ,**2023**, 102115. https://doi.org/10.1016/j.arr.2023.102115.

7. Nusser, A., Sagar, Swann, J.B., et al., Developmental dynamics of two bipotent thymic epithelial progenitor types, *Nature 2022 606:7912* (606) ,**2022**, 165–171. https://doi.org/10.1038/s41586-022-04752-8.

8. Bull, J.A., Moore, J.W., Mulholland, E.J., et al., MuSpAn: A Toolbox for Multiscale Spatial Analysis, ,**2025**, 2024.12.06.627195. https://doi.org/10.1038/s41467-026-75649-7.

9. Laidlaw, B.J., Duan, L., Xu, Y., et al., The transcription factor Hhex cooperates with the corepressor Tle3 to promote memory B cell development, *Nature Immunology* (21) ,**2020**, 1082–1093. https://doi.org/10.1038/s41590-020-0713-6.
