## Supplementary Tables for "Spatial remodeling of the thymic stroma with age disrupts niches for T cell selection and tolerance"

**Supplementary Table 1:** Antibody panel used for the flow cytometric analysis of the young and old thymi.

| Marker | Host species | Clone | Vendor | Catalogue number | Fluorophore | Dilution |
|---|---|---|---|---|---|---|
| AIRE | Rt | 5H12 | Invitrogen | 53-5934-82 | AF488 | 1:200 |
| CCR7 | Rt | 4B12 | BioLegend | 120120 | BV421 | 1:100 |
| CD117 | Rt | 2B8 | BioLegend | 105835 | BV711 | 1:200 |
| CD140a | Rt | APA5 | BioLegend | 135919 | PE-Cy5 | 1:400 |
| CD146 | Rt | ME-9F1 | BD OptiBuild | 741899 | BUV737 | 1:400 |
| CD24 | Rt | M1/69 | BD OptiBuild | 747717 | BV510 | 1:1000 |
| CD25 | Rt | PC61 | BioLegend | 102036 | BV605 | 1:1000 |
| CD26 | Rt | H194-112 | BioLegend | 137806 | FITC | 1:200 |
| CD31 | Rt | 390 | BD OptiBuild | 741084 | BUV496 | 1:500 |
| CD4 | Rt | GK1.5 | BioLegend | 100414 | APC-Cy7 | 1:400 |
| CD40 | Rt | 3/23 | BD OptiBuild | 745697 | BUV395 | 1:400 |
| CD45 | Rt | 30-F11 | BioLegend | 103174 | APC-Fire810 | 1:400 |
| CD5 | Rt | 53-7.3 | BioLegend | 100624 | PerCP-Cy5.5 | 1:400 |
| CD63 | Rt | NVG-2 | BioLegend | 143914 | PE-Dazzle594 | 1:400 |
| CD66a | Ms | CC1 | BD OptiBuild | 750946 | BUV661 | 1:400 |
| CD69 | Ar Hms | H1.2F3 | BioLegend | 104512 | PE-Cy7 | 1:400 |
| CD8a | Rt | 53-6.7 | BioLegend | 100730 | AF700 | 1:400 |
| CD80 | Ar Hms | 16-10A1 | BioLegend | BioLegend | BV605 | 1:400 |
| CD83 | Rt | Michel-19 | BioLegend | 121508 | PE | 1:400 |
| EpCAM | Rt | G8.8 | BioLegend | 118227 | BV605 | 1: |
| EpCAM | Rt | G8.8 | BioLegend | 118220 | PerCP-Cy5.5 | 1:500 |
| FOXP3 | Rt | FJK-16s | Invitrogen | 11-5773-82 | FITC | 1:200 |

| Marker | Host species | Clone | Vendor | Catalogue number | Fluorophore | Dilution |
|---|---|---|---|---|---|---|
| Helios | Ar Hms | 22F6 | BioLegend | 137216 | PE | 1:100 |
| Ly51 | Rt | 6C3 | BioLegend | 108314 | PE-Cy7 | 1:500 |
| MHCII | Rt | M5/114.15.2 | BioLegend | 107652 | APC-Fire750 | 1:1000 |
| PD1 | Rt | RMP1-30 | BioLegend | 109112 | APC | 1:200 |
| Podoplanin | Syr Hms | 8.1.1 | BioLegend | 127423 | BV421 | 1:200 |
| Sca-1 | Rt | D7 | BioLegend | 108143 | BV650 | 1:1000 |
| Streptavidin | - | - | BioLegend | 405249 | BV785 | 1:500 |
| TCRb | Ar Hms | H57-597 | BioLegend | 109210 | PE-Cy5 | 1:1000 |
| TER-119 | Rt | TER-119 | BioLegend | 116204 | Biotin | 1:400 |
| UEA-1 | - | - | Vector Laboratories | B-1065 | Biotin | 1:400 |

**Supplementary Table 2:** Antibodies used for the flow cytometric sort of young and old thymic stroma prior to snRNA/ATACseq (10x multiomics).

| Marker | Host species | Clone | Vendor | Catalogue number | Fluorophore | Dilution |
|---|---|---|---|---|---|---|
| CD45 | Rt | 30-F11 | BioLegend | **103128** | AF700 | 1:400 |
| CD19 | Rt | 6D5 | BioLegend | **115508** | PE | 1:500 |
| TER119 | Rt | TER-119 | BioLegend | **116206** | FITC | 1:500 |

**Supplementary Table 3:** Antibodies used for the thymic epithelial cell cultures.

| Marker | Host species | Clone | Vendor | Catalogue number | Fluorophore | Dilution |
|---|---|---|---|---|---|---|
| CD4 | Rt | RM4-5 | BioLegend | 100546 | BV650 | 1:500 |
| CD45 | Rt | 30-F11 | Invitrogen | MCD4517 | PE-TexasRed | 1:250 |
| CD5 | Rt | 53-7.3 | BioLegend | 100622 | PE-Cy7 | 1:500 |
| CD5 | Rt | 53-7.3 | BioLegend | 100624 | PerCP-Cy5.5 | 1:500 |

| Marker | Host species | Clone | Vendor | Catalogue number | Fluorophore | Dilution |
|---|---|---|---|---|---|---|
| CD8a | Rt | 53-6.7 | BioLegend | 100730 | AF700 | 1:400 |
| CD69 | Ar Hms | H1.2F3 | BioLegend | 104506 | FITC | 1:500 |
| CD83 | Rt | Michel-19 | BioLegend | 121508 | PE | 1:200 |
| EpCAM | Rt | G8.8 | BioLegend | 118220 | PerCP-Cy5.5 | 1:500 |
| EpCAM | Rt | G8.8 | BioLegend | 118227 | BV605 | 1:500 |
| Ly51 | Rt | 6C3 | BioLegend | 108314 | PE-Cy7 | 1:500 |
| MHCII | Rt | M5/114.15.2 | BioLegend | 107628 | APC-Cy7 | 1:1000 |
| Sca-1 | Rt | D7 | BioLegend | 108118 | AF647 | 1:1000 |
| Sca-1 | Rt | D7 | BioLegend | 108139 | BV785 | 1:1000 |
| TCRb | Ar Hms | H57-597 | BioLegend | 109208 | PE | 1:500 |
| UEA-1 | - | - | Vector Laboratories | FL-1061-2 | Fluorescein | 1:1000 |
| UEA-1 | - | - | Vector Laboratories; self-labelled | L-1060-2 | Cy5 | 1:1000 |

**Supplementary Table 4:** mIHC antibody panel, including host species, clone, vendor, catalogue number, fluorophore, dilution, and exposure time. DAPI was imaged in every cycle. Markers are listed in imaging order and were acquired in groups of three (see Methods); Ar Hms, Armenian hamster; Ms, mouse; Rb, rabbit; Rt, rat; Sy Hms, Syrian hamster.

| Marker | Host species | Clone | Vendor | Catalogue number | Fluorophore | Dilution | Exposure time [ms] |
|---|---|---|---|---|---|---|---|
| DAPI | - | - | - | - | - | 1:1000 | 150 |
| B220 | Rt | Ra3-6B2 | Akoya | 4150006 | AF488 | 1:200 | 200 |
| CD8A | Rt | 53-6.7 | Akoya | 4250017 | ATTO550 | 1:200 | 150 |
| CD69 | Ar Hms | H1.2F3 | Invitrogen | 14-0691-82 | ATTO550 | 1:200 | 300 |
| CD11C | Ar Hms | N418 | Akoya | 4350013 | Cy5 | 1:200 | 300 |
| CD45 | Rt | 30-F11 | Akoya | 4150002 | AF488 | 1:200 | 150 |
| XCR1 | Ms | ZET | BioLegend | 148202 | Cy5 | 1:100 | 300 |

| Marker | Host species | Clone | Vendor | Catalogue number | Fluorophore | Dilution | Exposure time [ms] |
|---|---|---|---|---|---|---|---|
| PDCA1 | Rt | eBio927 | Invitrogen | 16-3172-81 | AF488 | 1:200 | 300 |
| SIRPA | Rt | P84 | Invitrogen | 16-1721-81 | ATTO550 | 1:100 | 300 |
| CD5 | Rt | 53-7.3 | Akoya | 4250007 | ATTO550 | 1:200 | 250 |
| ERTR7 | Rt | ER-TR7 | Bio-Rad | MCA2402 | Cy5 | 1:400 | 80 |
| CD4 | Rt | RM4-5 | Akoya | 4250016 | ATTO550 | 1:200 | 150 |
| PDPN | Sy Hms | 8.1.1 | Invitrogen | MA5-18054 | AF488 | 1:200 | 300 |
| K8 | Rt | TROMA-1 | EMD Millipore | MABT329 | Cy5 | 1:200 | 80 |
| K14 | Rb | polyclonal | Invitrogen | PA5-16722 | AF488 | 1:200 | 80 |

**Supplementary Table 5:** Table of mIHC phenotype definitions, including the marker thresholds, distance thresholds, and regional filters used to identify each cell phenotype. "&" indicates a logical "AND" while "|" denotes a logical "OR" operator.

| Number | Phenotype name | Corresponding cell type | Defining markers and their normalized expression thresholds | Distance thresholds [pixels] | Region filter |
|---|---|---|---|---|---|
| 9 | SP8 (CD69-) | SP8 T-cells not expressing CD69 | (CD8a > 0.3) & (CD4 < 0.2) & ((TCRb > 0.15) \| (CD5 > 0.15)) & (CD69 <0.25) | - | Medulla |
| 11 | BC | B cells | B220 > 0.1 | 4 for BC clusters, 6 for interspersed BC | - |
| 13 | pDC (PDCA1+) | Plasmacytoid dendritic cells expressing PDCA1 | (CD11c > 0.25) & (PDCA1 > 0.25) | 11 | - |
| 14 | cDC1 (XCR1+) | Conventional dendritic cells expressing XCR1 | (CD11c > 0.25) & (XCR1 > 0.35) | 11 | - |
| 15 | cDC2 (SIRPa+) | Conventional dendritic cells expressing SIRPa | (CD11c > 0.25) & (SIRPa > 0.25) | 11 | - |
| 18 | SP4 (CD69-) | SP4 T-cells not expressing CD69 | (CD8a < 0.2) & (CD4 > 0.2) & ((TCRb > 0.2) \| (CD5 > 0.15)) & (CD69 <0.25) | - | Medulla |

**Supplementary Table 6:** IHC antibody panel used for TLS profiling, including clone and vendor information.

| Marker | Clone | Vendor |
|---|---|---|
| B220 | RA3-6B2 | BioLegend |
| ERTR7 | ERTR7 | Bio-Rad |
| PDPN | 8.1.1 | BioLegend |
| K8 | TROMA-1 | NICHD Hybridoma Bank |
| K14 | Poly19053 | BioLegend |
| CD21 | 7E9 | BioLegend |
| CD68 | FA-11 | BioLegend |
| GL7 | GL7 | BioLegend |
| PNAd | MECA-79 | BioLegend |
| CD19 | 6D5 | BioLegend |
| AID | mAID-2 | Thermo Fisher Scientific |
| CD138 | 281-2 | BioLegend |
| IgG | poly4060 | BioLegend |
| IgM | II/41 | eBioscience |
| CXCR5 | L138D7 | BioLegend |
| CCL21 | Polyclonal | LSBio |
| FOXP3 | FJK-16s | Invitrogen |
